# Evo2HiC: a multimodal foundation model for integrative analysis of genome sequence and architecture

**DOI:** 10.1101/2025.11.18.689171

**Authors:** Tangqi Fang, Yongkang Li, Xiao Wang, Zhiping Xiao, Shengqi Hang, Ghulam Murtaza, Junwei Yang, Hanwen Xu, Anupama Jha, William Noble, Sheng Wang

**Author notes:** These authors contributed equally (Tangqi Fang, Yongkang Li, Xiao Wang).

## Abstract

Understanding how genome sequences shape three-dimensional (3D) genome architecture is fundamental to interpreting diverse biological processes. Although previous studies have shown that sequence information can predict 3D genome architecture, these models fall short in capturing cell type–specific structures because they are trained solely on sequence inputs. The widely available Hi-C data, which contains rich structural information across biosamples, can provide complementary features to sequence data for studying cell type–specific architectures. Recently, DNA foundation models have demonstrated encouraging performance in capturing long-range genomic dependencies, holding promise for modeling chromatin interactions. However, the extremely high computational cost of running these models limits their applicability to Hi-C analysis, which requires genome-wide sequence embeddings. Here, we present Evo2HiC, a multimodal foundation model that jointly models genome sequences and structures to study cell type-specific chromatin structure. The key idea of Evo2HiC is to distill a large-scale DNA foundation model, Evo 2 (7B), into a compact encoder, while guiding the distillation with Hi-C data to preserve genomic features critical for 3D genome analysis. The model supports two types of encoders, one that operates directly on DNA sequences, and a second that additionally takes as input corresponding Hi-C data. Using the DNA-only encoder and predicting raw Hi-C contact matrices, Evo2HiC improved distance-stratified Spearman correlation by 10.9% over Orca. Moreover, by jointly embedding Hi-C and sequence information Evo2HiC achieved the best overall Pearson correlation when predicting five representative epigenomic assays. Interpretation analysis of Evo2HiC revealed its ability to identify cell type–specific sequence motifs that explain changes in epigenomic signals. Finally, we demonstrated the cross-species generalizability of Evo2HiC on 177 species from the DNA Zoo dataset for Hi-C resolution enhancement. In summary, Evo2HiC is a foundation model that integrates genome sequences and 3D chromatin structure information, substantially reduces computational cost while maintaining state-of-the-art accuracy on predicting various epigenomic signals and genome architecture, enables the identification of cell type-specific motifs, and demonstrates robust generalizability across species.

## Introduction

Understanding how genetic information flows from the DNA sequence to downstream biological processes is fundamental across diverse biological contexts, including development [1, 2, 3], evolution [4, 5, 6], and disease [7, 8, 9, 10, 11, 12]. Within the nucleus, this genetic information is organized into a three-dimensional structure that profoundly influences essential genome functions, including DNA replication, transcription, cell differentiation, and cellular senescence [13]. To characterize this spatial organization, a variety of experimental techniques, such as Hi-C [14], ChiA-PET [15], GAM [16], SPRITE [17], and HiChIP [18], have been developed to map chromatin architecture and DNA–DNA interactions across the genome, generating a large number of contact matrices in which the axes represent genomic loci and the matrix values reflect their spatial proximity in three dimensions.

Meanwhile, genomic sequence encodes features such as CTCF binding motifs and CpG islands that act as key determinants of chromatin loops, domains, and compartments, thereby shaping the three-dimensional genome architecture [19, 20]. Building on this principle, several computational methods, including Akita [21], DeepC [22], and Orca [23], have been developed to predict chromatin organization directly from DNA sequences, revealing the underlying relationships between chromatin sequences and structure.

However, because current approaches are trained solely using sequences as inputs, they need to be retrained when analyzing a new cell type. This failure to generalization across cell types makes it challenging for these models to characterize cell type-specific chromatin interaction patterns. One potential solution is to jointly train a model using genome sequences and Hi-C data. In contrast to genome sequences, which allow us to extract high-resolution 1D features, a Hi-C contact matrix directly delivers 2D patterns but is typically analyzed at coarse resolution (e.g., 10 kb [14]) due to sequencing depth constraints. Therefore, high-resolution 1D features derived from genome sequences and 2D features from low-resolution contact matrices provide complementary information for deciphering cell type-specific chromatin structures.

Jointly analyzing Hi-C data and genome sequences requires a model to process millions of bases as input, posing substantial challenges for capturing long-range dependencies. Recently, a series of DNA sequence foundation models have demonstrated strong performance across diverse genomic tasks [24, 25, 26, 27] due to their ability to model long-range dependencies. For instance, the recently developed DNA foundation model Evo 2 can process sequence windows up to one million base pairs [28], demonstrating its capacity to capture distal genomic interactions. Despite this potential, applying DNA foundation models to chromatin structure prediction remains computationally demanding for several reasons. First, Hi-C data represent pairwise genomic interactions, requiring simultaneous analysis of two DNA sequences, which doubles computational cost. Second, the dense attention mechanisms and expansive context window in these foundation models create massive computational redundancy, especially when downstream tasks focus on specific sub-matrices. Third, models such as Evo 2 require high-end GPUs, such as NVIDIA H100s, to support efficient inference, further limiting accessibility for many experimental laboratories lacking such computational resources.

To this end, we propose Evo2HiC, a multimodal foundation model for jointly analyzing DNA sequence and Hi-C contact matrices. The key idea is to distill the powerful but expensive DNA foundation model Evo 2 into a much smaller model that maintains the long-range modeling ability of Evo 2 but requires fewer computational resources. Instead of directly distilling Evo 2 to a small model, we jointly project Evo 2 and our model to a shared Hi-C embedding space. This approach allows the distillation to focus on features that are critical to the chromatin structure. The resulting embedding space can be used as an encoder for downstream applications that involve Hi-C data and DNA sequence data. This distilled encoder reduces the time and memory requirements for downstream applications by a factor of 500 compared to Evo 2.

We demonstrate the versatility, effectiveness, and interpretability of Evo2HiC for studying cell type-specific chromatin architecture. We observe a strong consistency between Evo 2–based embeddings and those produced by Evo2HiC. On raw Hi-C contact matrices predicted directly from sequence, Evo2HiC improves the distance-stratified Spearman correlation by 10.9% over Orca [29]. Moreover, Evo2HiC achieves state-of-the-art performance in predicting five representative types of epigenomic signals, including DNase, CTCF, H3K27ac, H3K27me3, and H3K4me3. Interpretation of the model further reveals cell type–specific sequence motifs that explain observed changes in epigenomic activity. The distilled encoder in Evo2HiC also exhibits strong cross-species generalizability when using only Evo 2 human embeddings. By jointly leveraging genomic sequences and low-coverage contact matrices, Evo2HiC achieves the best performance over four existing methods for Hi-C resolution enhancement across 177 species.

In summary, Evo2HiC is a multimodal foundation model that jointly models genome sequences and 3D chromatin structures. By distilling large-scale DNA language models into a compact architecture grounded in Hi-C data, Evo2HiC substantially reduces computational cost while maintaining high predictive accuracy across diverse tasks, including predicting the Hi-C contact matrix from DNA sequence, epigenomic signal prediction, and Hi-C resolution enhancement, and further demonstrates cell type-specific interpretability and cross-species generalizability.

## Results

### Overview of Evo2HiC

Evo2HiC aims to learn a lightweight encoder capable of capturing megabase-scale long-range interactions that are critical for Hi-C data analysis (**Fig. 1**). To achieve this, we distill the 7 billion-parameter Evo 2 model into a compact CNN-based genome sequence encoder consisting of seven layers and totaling 3.6 million parameters, trained on 1.2 million 2 kb human genomic bins. Given the nearly 2000-fold reduction in model size, it is infeasible for the CNN encoder to retain all information from Evo 2. Therefore, we encourage the model to focus on genomic features most relevant to the chromatin structure. To this end, we leverage Hi-C data to guide the distillation process. Specifically, we optimize two contrastive learning objectives: one aligns the sequence embeddings from the CNN encoder with those from Evo 2, and the other aligns the CNN-based embeddings with Hi-C patch embeddings produced by a Hi-C encoder. We deliberately avoid direct alignment between Evo 2 and the Hi-C encoder to mitigate the computational burden arising from large-scale 2D contact matrices and genome-wide embeddings. Following these two alignment steps, the resulting CNN-based module serves as our genomic sequence encoder. The co-embedded representation, which concatenates embeddings from both Hi-C contact maps and genomic sequences across the entire human genome, further enables downstream applications, such as Hi-C resolution enhancement and epigenomic signal prediction.

**Figure 1:**
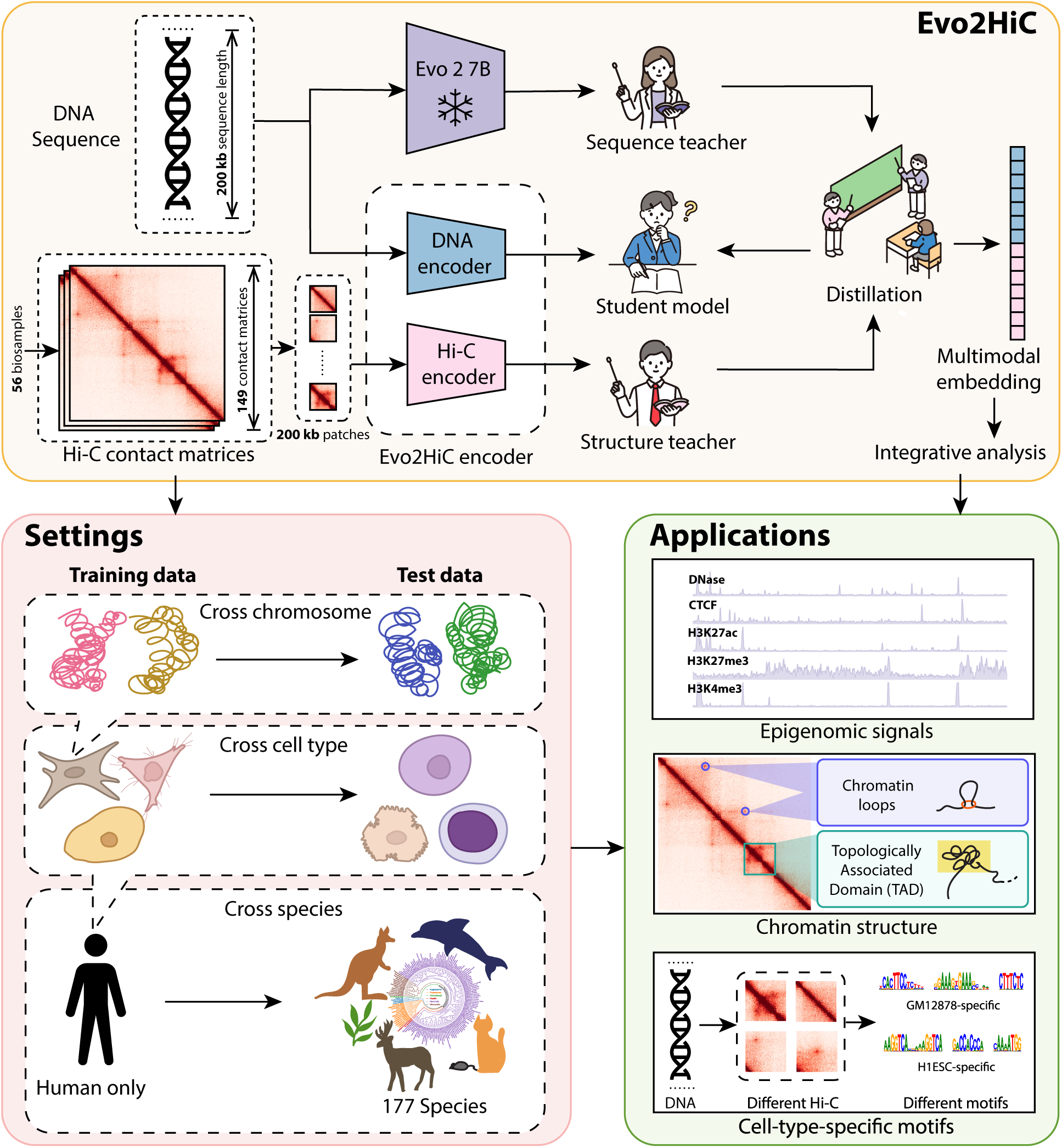
Overview of Evo2HiC. Evo2HiC is a multimodal foundation model that integrates Hi-C contact map and DNA sequence at 2kb resolution. It exploits distillation to learn a compact CNN-based sequence encoder from large-scale DNA foundation model Evo 2, while encouraging this distillation to focus on genomic features that are more critical to the chromatin structure. By integrating sequence and contact matrix, Evo2HiC allows cross cell-type and cross-species generalizability, further improving the performance on predicting epigenomic signals, chromatin structure analysis, and cell type-specific motif identification.

### Evo2HiC preserves the genomic sequence features from Evo 2

We first examined whether the distilled CNN-based encoder can preserve genomic sequence features from Evo 2. To this end, we compared pairwise similarities between 2 kb human genomic sequences embedded by Evo 2 and by Evo2HiC (**Fig. 2a,b**). The embeddings showed a strong correlation of 0.63, markedly higher than the 0.13 correlation achieved by the same CNN architecture trained without Evo 2 distillation (see **Methods**). Given that Evo 2 was pretrained on 128,411 species, we next asked whether the distilled encoder trained only on human data could generalize to another species, such as mouse. Indeed, when evaluated on 78,255 2 kb mouse genomic sequences, Evo2HiC achieved substantially higher correlation with Evo 2 compared to the non-distilled CNN encoder (**Fig. 2c,d**), indicating successful transfer of cross-species genomic representations.

**Figure 2:**
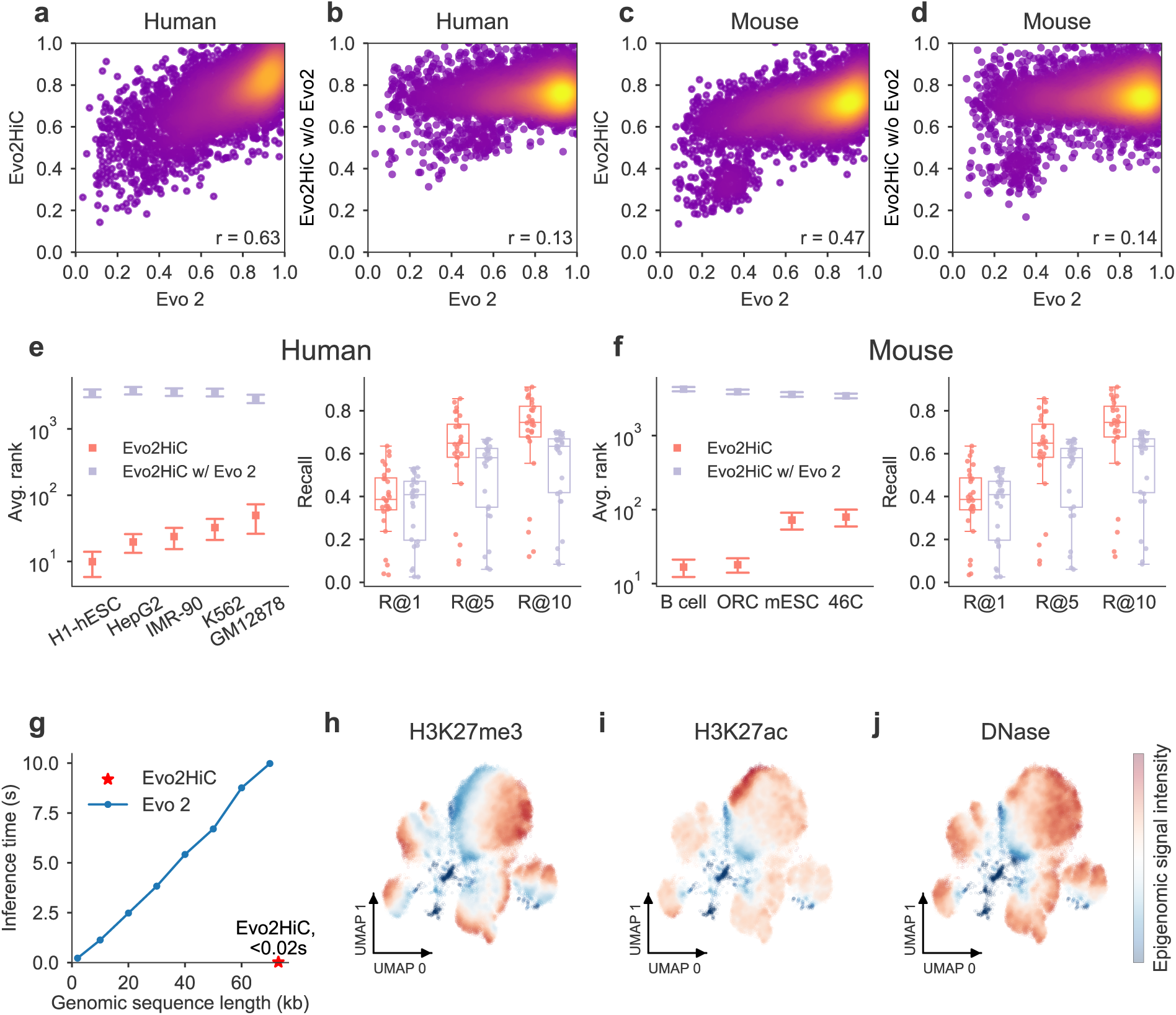
Examining the embedding space of Evo2HiC. a,c, Embedding-space alignment between Evo2HiC and Evo 2. Pairs of 2 kb sequences are randomly sampled: the x-axis is the cosine similarity of the pair in Evo 2 embedding space; the y-axis is the same pair’s cosine similarity in Evo2HiC’s embedding space. The strong correlation (*r*) indicates that Evo2HiC preserves Evo 2’s pairwise similarity structure. **b,d,** The same comparison for an Evo2HiC variant without the Evo 2 distillation. The decrease in *r* relative to panels a/c quantifies the contribution of the Evo 2 distillation objective. **e,f,** Comparison between Evo2HiC and the Evo2HiC without Evo 2 on retrieving the most similar Hi-C patch using the corresponding genomic sequences in terms of average rank and recall on human (**e**) and on mouse (**f** ). In the plot, boxes represent mean rank, with an error bar of 10*×*SE (standard error). For recall, the boxes represent the 25–75 percentile range with an error bar of 1.5*×*IQR (interquartile range) **g,** Comparison of inference cost for genomic sequences of different lengths between Evo 2 and Evo2HiC. The Evo 2 cost increases with larger genomic sequence length, whereas Evo2HiC (red star) has a low inference cost. **h-j,** UMAP visualization of the embeddings of genomic sequences by Evo2HiC, colored by epigenomic signals of H3K27me3 (**h**), H3K27ac (**i**), DNase (**j**).

Having confirmed that our model retains Evo 2–derived genomic features, we next assessed whether these features benefit Hi-C analysis. On a held-out set of genomic sequence and Hi-C patch pairs, we compared the performance of identifying the corresponding Hi-C patch using sequence embeddings distilled from Evo 2 (**Fig. 2e-f, Extended Data Fig. 1**). Our embeddings achieved higher recall and lower average rank than those from the encoder trained without Evo 2 on both human and mouse datasets, demonstrating the relevance of Evo 2 features for capturing chromatin organization. The performance gain was more pronounced in mouse, whose sequence data were not explicitly included during training, suggesting improved cross-species generalizability likely stemming from Evo 2’s multi-species training.

Beyond accuracy, Evo2HiC offers substantial computational efficiency. In contrast to Evo 2’s computationally expensive inference, Evo2HiC employs an efficient CNN-based architecture, achieving up to 500-fold faster inference at a genomic sequence length of 70 kb (**Fig. 2g, Supplementary Figure 1**).

Finally, we explored whether the co-embedding space learned from genomic sequences and Hi-C contact matrices could capture functional genomic activity. Visualization of the epigenomic scores from different assays (H3K27me3, H3K27ac, DNase, CTCF, H3K4me3) revealed clear spatial patterns in the co-embedding space (**Fig. 2h-j, Supplementary Figure 2**), highlighting its potential for predicting diverse epigenomic signals.

### Evo2HiC predicts Hi-C contact matrix using genome sequences

Before jointly analyzing genome sequence and Hi-C data, we first verified that the sequence encoder in Evo2HiC is structure-aware, retaining sequence-derived information to recover 3D contact patterns (**Fig. 3a–h, Extended Data Fig. 2, Supplementary Figure 4**). Specifically, a U-Net–based decoder was trained to generate Hi-C contact patches from the learned sequence embeddings. We compared Evo2HiC with Orca, which employs a hierarchical sequence encoder to predict contact matrices, and with Evo 2, which uses the same U-Net–based decoder as our model. The latter comparison serves as an ablation to assess the intrinsic quality of our sequence encoder.

**Figure 3:**
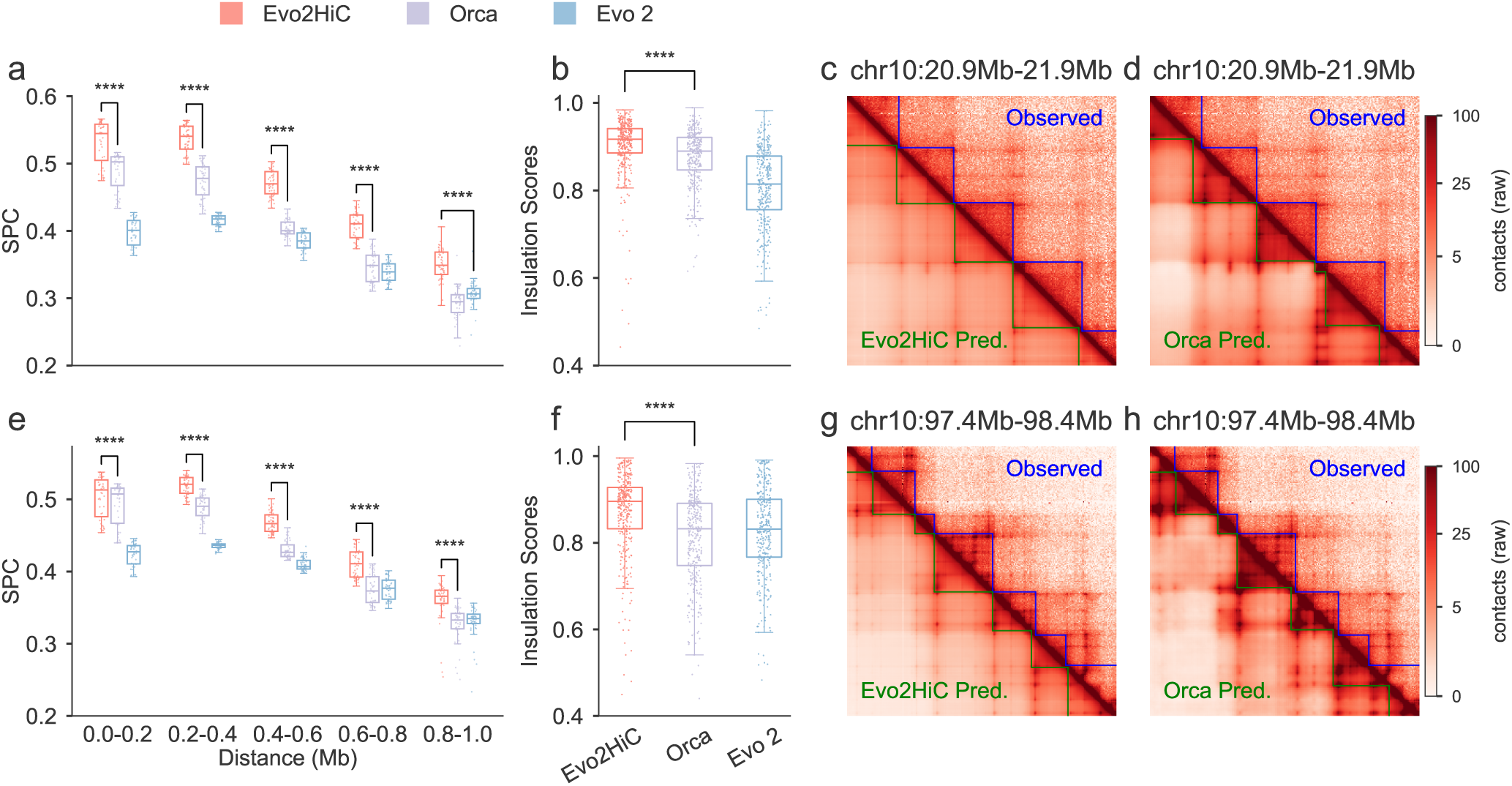
Genome structure prediction from sequence. a–h, Comparison among Evo2HiC, Orca, and Evo 2 for predicting Hi-C contact maps from genomic sequence input. Metrics are Spearman correlation on Hi-C contact counts **(a, e)** and Pearson correlation on insulation score **(b, f)**. In each box plot the central line marks the median, the box spans the 25th–75th percentile, and whiskers extend to 1.5*×*IQR, with individual values overlaid as jittered points. **(c, d, g, h)** Each matrix is split along the main diagonal: the upper triangle shows the observed Hi-C contact map, and the lower triangle shows the corresponding model prediction; **blue segments** outline TAD boundaries called on the observed (upper) matrix; **green segments** outline boundaries called on the predicted (lower) matrix. **a–d** are results on the H1ESC cell line and **e–h** are results on the HFF cell line.

Across two human cell lines, H1ESC and HFF, Evo2HiC outperformed both Orca and Evo 2 in terms of Spearman correlation coefficient (SPC) on the Hi-C contact counts (**Fig. 3a, e, Extended Data Fig. 2a–d**). Specifically, we tiled held-out test chromosomes (chr 9 and chr 10) with 1 Mb windows at 4 kb resolution using a 500 kb step size, computed distance-stratified SPC within each window, and achieved a 10.9% improvement over Orca (see **Methods** for details). When evaluated by insulation score (**Fig. 3b, f, Extended Data Fig. 2e,f**), which reflects the organization of topologically associating domains (TADs), Evo2HiC also achieved the highest performance, demonstrating its capacity to capture fine-grained 3D chromatin features. Taken together, these results indicate that the distilled sequence encoder is topology-aware and preserves the 3D contact information necessary for the downstream analyses described in the following sections.

We further compared Evo2HiC with Orca in Orca’s original setting. For a fair comparison, we trained Evo2HiC on the same Micro-C target representation and the same auxiliary epigenomic peak sets that Orca uses, and evaluated both models in Orca’s own setting: its 417 held-out 1 Mb windows on chromosomes 9 and 10, its target matrices and its validity mask (see **Methods**). On Orca’s reported metric, the pooled correlation of log(observed/expected) contacts, Evo2HiC was comparable to Orca in both cell lines (**Extended Data Table 1**), while being fine-tuned from its pretrained encoder on an order of magnitude fewer training samples. On insulation score correlation, observed/expected correlation and mean squared error [30], Evo2HiC was significantly more accurate than Orca in both cell lines (for example, insulation score correlation 0.940 versus 0.887 in H1ESC), indicating that it better recovers domain-level organization at comparable pixel-level accuracy.

### Evo2HiC predicts epigenomic profiles

Next, we leveraged the joint embedding of Hi-C and genomic sequences learned by Evo2HiC to predict epigenomic signals using a transformer-based decoder trained on five representative assays: DNase accessibility and ChIP-seq for CTCF, H3K27ac, H3K27me3, and H3K4me3. In the cross-chromosome evaluation, Evo2HiC substantially outperformed the Hi-C–only baseline and Evo 2, both of which used the same decoder as Evo2HiC but took only Hi-C data alone or Evo 2 embeddings as input, achieving an average improvement of 34.7% over the Hi-C–only baseline and 26.2% over Evo 2 across all assays in terms of Pearson correlation coefficient (PCC) (**Fig. 4a,b, Supplementary Figure 5a,b**). Under the more challenging cross-cell line and cross-chromosome setting, Evo2HiC also surpassed both models in three out of five assays (**Fig. 4c, Supplementary Figure 5c**). We also benchmarked Evo2HiC against the state-of-the-art sequence-to-epigenome foundation models AlphaGenome [31] and Borzoi [32] on genomic regions shared by their respective test sets and Evo2HiC’s test set (**Extended Data Fig. 3**). Although AlphaGenome and Borzoi use fragment-level rather than cross-chromosome data splits, Evo2HiC was evaluated under a stricter cross-chromosome split and achieved comparable performance, outperforming Borzoi by 5.4% in PCC while trailing AlphaGenome by only 0.87%. Evo2HiC was also more accurate than HiCFoundation [33], a Hi-C foundation model with an epigenomic prediction head whose training included our test chromosomes, in 14 of the 15 cell line and assay combinations, including the held-out K562 cell line (**Extended Data Table 2a**). To evaluate the contribution of our knowledge distillation strategy, we performed an ablation study comparing Evo2HiC against both a from-scratch training baseline (no-distillation) and a direct combination of raw Evo 2 embeddings with Hi-C data (Evo 2 + HiC) (**Extended Data Fig. 4**). Under the cross-chromosome setting, the distillation approach consistently enhanced predictive accuracy, outperforming the no-distillation variant and Evo 2 + HiC baselines by 2.0% and 3.8%, respectively. This advantage also generalized to the more challenging cross-cell-type and cross-chromosome setting, where Evo2HiC maintained a 2.9% and 3.1% improvement over the no-distillation variant and Evo 2 + HiC models, respectively.

**Figure 4:**
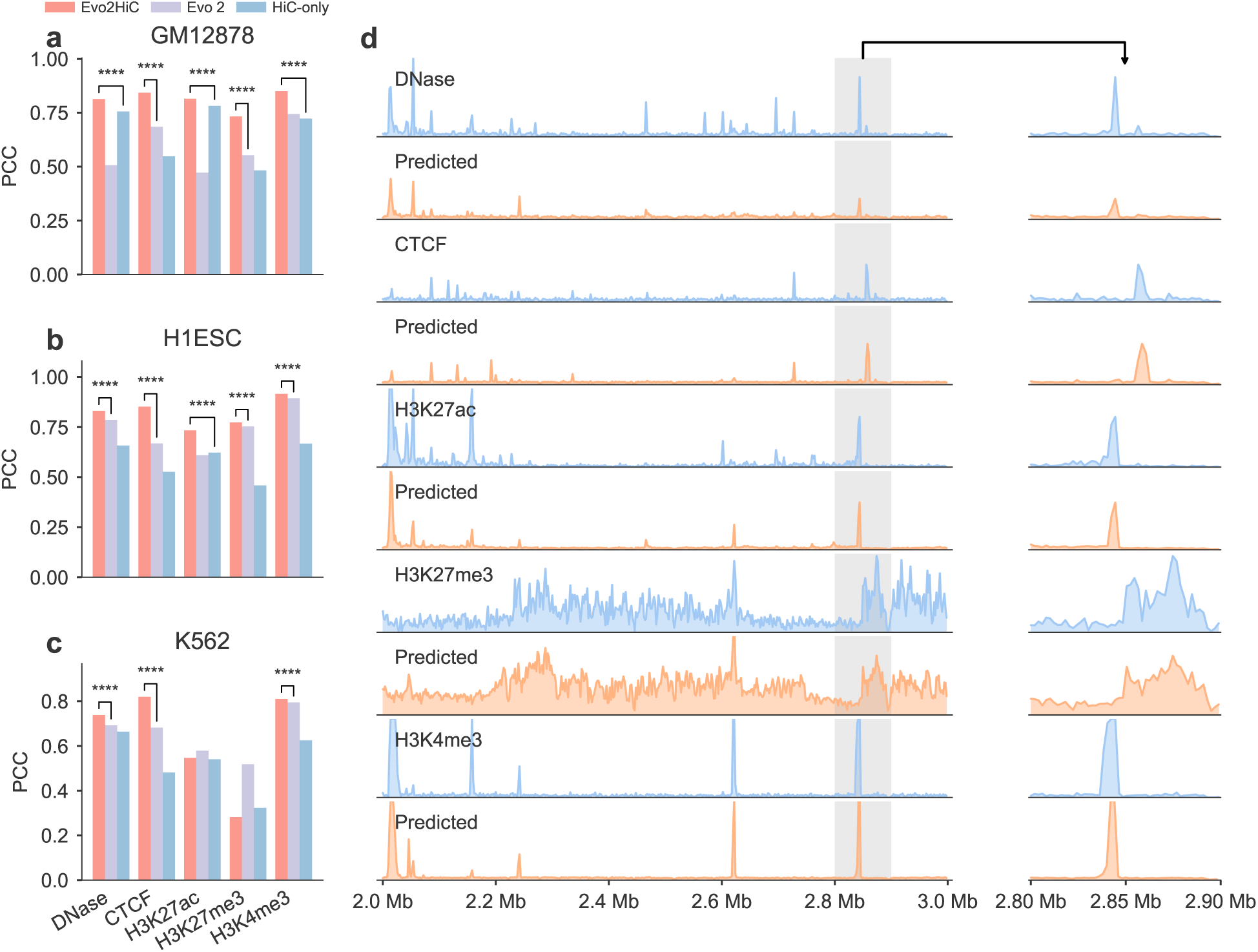
Comparison on predicting epigenomic signals. a,b, Comparison on predicting epigenomic signals in the cross chromosome setting on GM12878 (**a**) and H1ESC (**b**). **c,** Comparison on predicting epigenomic signals in the cross cell line and cross chromosome setting on K562. For each assay, a statistical test is only performed for the performance gain between Evo2HiC and the best baseline. **d,** Comparison between the predicted epigenomic signal by Evo2HiC and the observed epigenomic signals in the GM12878 cell line (chr9:2 Mb - 3 Mb).

Further analysis reveals that Evo2HiC not only captures fine-grained genomic activity (**Fig. 4d**) but also accurately resolves cell type-specific regulatory variations. To evaluate this cross-cell type capability, we constructed a cross-cell type prediction matrix (**Extended Data Fig. 5a**) and assessed the PCC between the differential ground-truth signals and the corresponding differential predictions across different cell lines (**Extended Data Fig. 5b, c**). Evo2HiC substantially improved the prediction of cell type-specific differences for CTCF, H3K4me3 and DNase, most markedly for CTCF (PCC 0.366 versus 0.195 for the strongest baseline), with confidence intervals of the improvement well above zero for all three assays (**Supplementary Table 13**). These gains indicate that the model dynamically leverages cell type-specific information rather than memorizing cell type-invariant sequence features, reflecting its ability to integrate sequence representation and structural information. For H3K27ac, Evo2HiC performed on par with the Hi-C–only baseline, and for H3K27me3 it was below the sequence-only baseline; representative loci for both successful and unsuccessful assays are shown in **Supplementary Figure 15**.

Beyond signal prediction, the joint profiling of Hi-C, DNase, and ChIP-seq enabled by Evo2HiC can be directly leveraged to nominate enhancer-gene regulatory links. By integrating these predicted modalities, we computed the Activity-by-Contact (ABC) score [34] and evaluated it against CRISPRi perturbation data in K562, a cell line held out from both pretraining and fine-tuning. To avoid evaluating on chromosomes used for training, we used two genome-wide CRISPRi screens [35, 36] restricted to the held-out chromosomes 9 and 10. For distal elements (≥ 50 kb from the transcription start site; 18 validated enhancers), ABC scores computed from Evo2HiC’s predicted tracks ranked validated enhancers markedly higher than genomic distance or Hi-C contact alone (AUPRC 0.221 versus 0.039 for distance; **Extended Data Table 3a**). An example locus (*PPIF* in GM12878) is shown in **Supplementary Figure 7**. These results illustrate how Hi-C and epigenomic profiles predicted by Evo2HiC can be used to nominate distal enhancers in an unseen cell type.

The combination of Hi-C and predicted ChIP-seq further enables study of cellular regulatory networks. From the Hi-C contact maps, we identified chromatin loops using HiCCUPS [14] and classified their anchors according to the predicted epigenomic signals, including in the held-out K562 cell line. Using the ENCODE registry of candidate cis-regulatory elements [37] as the reference classes, anchor classifications derived from Evo2HiC’s predicted profiles agreed with those derived from measured signals better than those of the Hi-C–only and sequence-only baselines in all three cell lines, including the held-out K562 (**Extended Data Table 3b**). Two readouts that use no measured epigenomic data further support these classifications (**Extended Data Table 3c**). First, anchors that Evo2HiC predicted to be CTCF-bound were enriched for CTCF motifs identified from sequence alone [38, 39], on par with measured CTCF ChIP-seq, and loops whose two anchors were both predicted CTCF-bound were mostly convergently oriented. Second, genes whose promoters lay in anchors predicted to be promoter-like were more highly expressed than genes in other anchors. We further highlight representative CTCF, active, and repressive loops, whose anchors display distinct epigenomic signatures that align closely with established biological expectations (**Supplementary Figure 9**). Under a rule-based classification that assigns H3K27me3-marked anchors as repressive first, the advantage in K562 was reduced, reflecting the weaker held-out prediction of H3K27me3 (**Supplementary Figure 8, Supplementary Table 14**).

### Evo2HiC identifies interpretable cell type-specific motifs

Identifying motifs that drive epigenomic signal variation is essential for understanding genome function. Sequence-only models such as Orca and Akita can be interpreted by applying attribution methods such as SHAP [40] to their predictions; however, such interpretability is tied to a fixed downstream model head trained for a specific cell line, so obtaining cell type–specific motifs requires training and analyzing a separate model for each cell type. Evo2HiC offers a complementary, multimodal alternative: by taking Hi-C as part of the input, applying the same attribution method under different Hi-C contexts yields distinct attribution distributions over the same DNA sequence within a single model. We exploit this property to compute attribution scores for each base pair conditioned on the supplied Hi-C contact map, and to identify motifs with enriched attribution scores for each cell type (see **Methods** for details).

We applied our cell type–specific interpretability framework to two human cell lines: H1ESC and GM12878 (**Fig. 5a**). By using independent gene expression data from the ENCODE project [41], we computed a differential expression score (Δ log(expression)) of each motif between two cell lines. We observed that cell type–specific motifs exhibited substantially higher differential expression compared to shared motifs (**Fig. 5b,c**). As an additional validation, our framework identified the CTCF motif as a shared motif across both cell types, consistent with its well-established role in chromatin organization (**Fig. 5d–f**).

**Figure 5:**
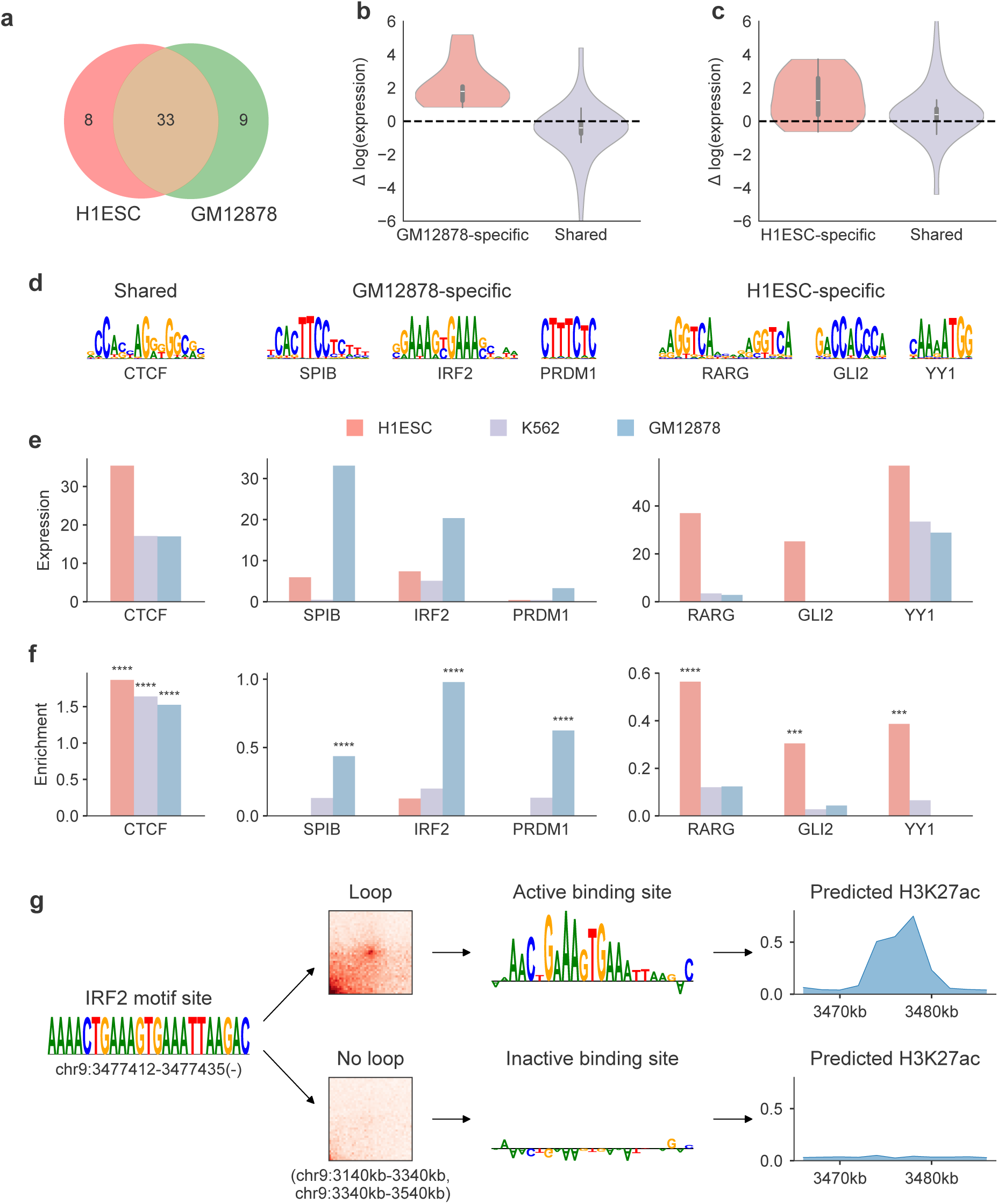
Identifying cell type-specific motifs for interpreting epigenomic signal changes. a, Number of shared and cell type-specific motifs identified for GM12878 and K562 by Evo2HiC. **b,c,** Log expression change between cell type-specific motifs and shared motifs on GM12878 (**b**) and H1ESC (**c**). **d,** Shared and cell type-specific motifs identified for GM12878 and H1ESC by Evo2HiC. **e,f,** Gene expression (**e**) and attribution enrichment (**f** ) of different motifs across two cell types. Attribution enrichment is calculated by the SHAP attribution score of that motif against random sequences. CTCF is a shared motif. SPIB, IRF2, PRDM1 are identified GM12878-specific motifs. RARG, GLI2, YY1 are identified H1ESC-specific motifs. **g,** Evo2HiC reveals the active binding site for a cell type-specific motif for more fine-grained interpretation of epigenomic signal changes.

For cell type–specific motifs, we further observed higher expression levels in their corresponding cell lines. To further quantitatively assess motif relevance, we introduced an attribution enrichment score (**Fig. 5c**), which measures the proportion of motif-associated regions receiving high attribution scores relative to background regions. All cell type–specific motifs exhibited strong attribution enrichment, demonstrating Evo2HiC ’s ability to precisely identify regulatory motifs that are both structurally and transcriptionally relevant in a cell type–specific manner. As a control, the K562 cell line showed neither high expression nor elevated attribution scores for motifs specific to H1ESC and GM12878, confirming the specificity of our findings.

Moreover, the discovered cell type–specific motifs align well with previous studies. For instance, IRF family motifs were specifically enriched in GM12878, consistent with their established role in mediating cell type–specific transcription factor binding [42]. RARG binding motifs were identified in H1ESC, consistent with prior work linking RARG to retinoic acid–induced enhancer activation and increased H3K27ac in embryonic stem cells [43]. These observations support the biological relevance of the motifs recovered by our framework.

Beyond cell type–specific motif identification, the multimodal formulation of Evo2HiC also enables direct analysis of how 3D contacts shape regulatory activity, providing more fine-grained interpretability. By contrasting epigenomic signal predictions between contact matrices with and without the chromatin loop, we can distinguish active from inactive binding sites, further elucidating the context-dependent activity of regulatory motifs (**Fig. 5g**).

### Evo2HiC demonstrates strong generalizability across 177 species

As Evo 2 is pretrained on multiple species, we hypothesized that the genomic sequence encoder distilled from Evo 2 using only human sequences might generalize to other species. To verify this, we evaluated Evo2HiC on the Hi-C resolution enhancement task, which takes genomic sequences and low-coverage Hi-C matrices as input and aims to reconstruct the corresponding high-resolution contact matrices. In this setting, a U-Net–based decoder was used to generate Hi-C contact patches from the combined embeddings of genomic sequences and Hi-C patches.

We first evaluated performance on 14 human contact maps and observed that Evo2HiC achieved the best results across all four evaluation metrics (**Fig. 6a**), demonstrating the effectiveness of jointly leveraging sequence and contact information for Hi-C resolution enhancement. To assess cross-species generalization, we next applied Evo2HiC to 11 mouse contact maps, where the mouse genomic sequence embeddings were generated without access to Evo 2 mouse embeddings (**Fig. 6b, Supplementary Figure 10**). Nevertheless, Evo2HiC still achieved the best performance, confirming its ability to generalize beyond the species used for distillation. We further compared Evo2HiC with the recent Hi-C foundation model HiCFoundation [33]. Because HiCFoundation holds out different chromosomes from ours, our test chromosomes were part of its training data; even so, Evo2HiC outperformed it on SPC, SSIM and PSNR on the human maps and on SSIM and PSNR on the mouse maps (**Extended Data Table 2b**). To confirm that this performance boost was driven by the learned sequence representations, we performed an ablation study removing the DNA input. The Evo2HiC (Hi-C only) baseline exhibited a significant performance drop on both human and mouse datasets (**Extended Data Fig. 6**), confirming that the distilled sequence encoder extracted informative features for both human and mouse.

**Figure 6:**
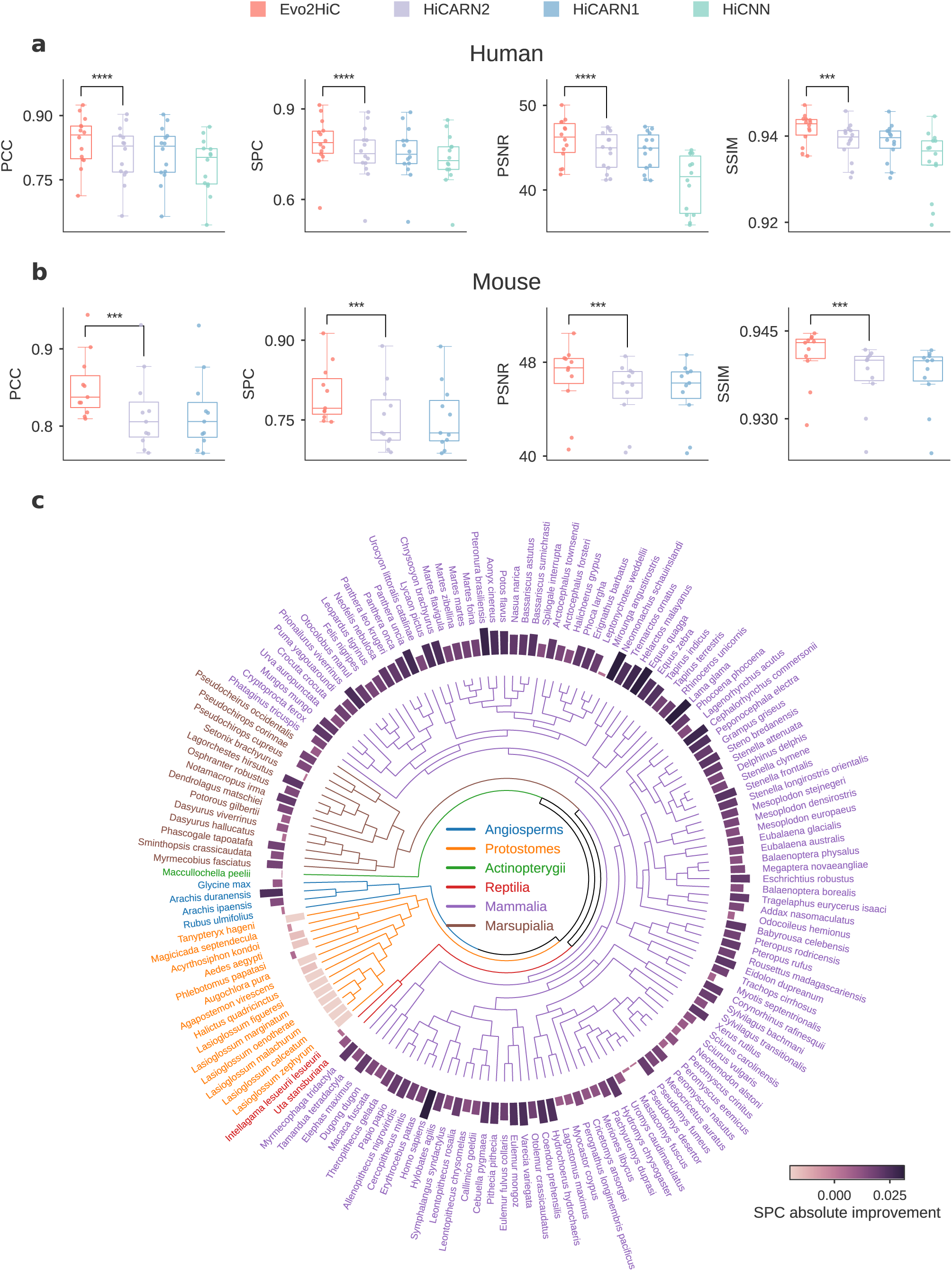
Evo2HiC enables cross-species analysis on the DNA Zoo data. a,b, Comparison on resolution enhancement in human (**a**) and mouse (**b**) across four metrics. **c,** Comparison on resolution enhancement on 177 species using DNA Zoo. Bar heights indicate the relative improvement over the best baseline HiCARN2 by Evo2HiC in terms of Spearman correlation.

We extended the evaluation to the DNA Zoo dataset by collecting genomic sequences and Hi-C contact matrices from 177 species (**Fig. 6c, Supplementary Figure 11**). Without utilizing Evo 2 embeddings for these species, Evo2HiC still outperformed the representative baseline in 157 out of 177 species in terms of SPC, and outperformed HiCFoundation in pixel-level PCC in 127 of 177 species (**Extended Data Table 2b**). At the level of TAD boundaries, evaluated with the same protocol as in the analyses above, Evo2HiC also achieved a higher TAD-boundary F1 than HiCARN2 (123 of 177 species) and HiCFoundation (133 of 177 species; **Supplementary Table 15**).

Moreover, the biodiversity of the DNA Zoo dataset allowed us to investigate where the sequence prior learned from human data contributes. Compared with the same model run on random DNA and with the Hi-C–only ablation, the genome sequence improved pixel-level accuracy in nearly all placental mammals and marsupials, with a gain that decreased with evolutionary distance from human (**Extended Data Fig. 7, Supplementary Table 15**). In these species, the sequence also shifted predicted domain boundaries towards motifs of CTCF, a well-established vertebrate boundary factor, whereas no such shift was observed in protostomes and angiosperms (**Supplementary Table 16, Supplementary Figures 12 and 16**). Thus, the sequence prior learned from human data generalizes across mammals, consistent with their shared determinants of 3D genome organization such as CTCF, whereas its contribution to more distant clades, which lack these shared factors, remains to be established.

## Discussion

We have presented Evo2HiC, a lightweight and effective computational framework for jointly analyzing Hi-C contact matrices and genomic sequences at 2 kb resolution. The key idea of Evo2HiC is to distill a large-scale DNA foundation model (Evo 2) into a much smaller encoder, while guiding the distillation with Hi-C data to retain genomic features critical for three-dimensional genome analysis. We systematically evaluated Evo2HiC across three major applications: (1) predicting Hi-C contact matrix from genomic sequences, (2) predicting epigenomic profiles from joint Hi-C and sequence embeddings, and (3) enhancing Hi-C resolution by integrating Hi-C and sequence. The superior performance across all tasks confirms the effectiveness of our joint modeling framework. Moreover, Evo2HiC enables interpretable analysis by identifying cell type–specific sequence motifs that explain the observed epigenomic profiles, offering mechanistic insights into how 3D genome architecture regulates essential genomic functions.

Despite the promising results of Evo2HiC, several limitations remain that we aim to address in future work. First, beyond Evo 2, a growing number of DNA foundation models [24, 25, 26] have demonstrated strong performance across diverse genomic tasks. We plan to investigate how distilling a compact encoder from multiple foundation models could further enhance the robustness and generalizability of our model. Second, while our results on predicting epigenomic signals has demonstrated potential for studying gene ex-pression, we intend to extend it to directly predict RNA-seq profiles [44], which could provide deeper insights into how 3D chromatin organization influences gene expression. Third, in the current work we extracted embeddings from Evo 2 by freezing its parameters, as the model’s size makes fine-tuning computationally challenging. We plan to develop efficient fine-tuning strategies [45, 46] that enable adapting Evo 2 with Hi-C data, potentially unlocking its full capacity for chromatin structure analysis. Fourth, the joint mod-eling of sequence and chromatin context positions Evo2HiC as a natural starting point for predicting the impact of structural variants (SVs) on 3D genome organization. Extending Evo2HiC to population-scale SV-effect prediction, for instance on the UK Biobank[47], represents a promising direction for translating sequence–structure modeling into genomic insights.

## Methods

### Overview of Evo2HiC

Evo2HiC has three main components: the Hi-C encoder, the DNA encoder, and task-specific heads for downstream applications.

The Hi-C encoder employs a CNN-based architecture, comprising an initial cross-embedding layer and two residual blocks. The cross-embedding layer applies three convolutional operations to the Hi-C matrices with kernel sizes of 3, 7, and 15, producing feature maps of dimensions 64, 32, and 32, respectively. These are concatenated to form a 128-dimensional Hi-C embedding. Subsequently, two residual blocks are incorporated to enhance the encoder’s representational capacity. In addition to standard residual blocks, ours are modified to adapt to multiple resolutions of Hi-C, allowing for the capture of chromatin architecture at different scales: the input embedding is pooled at multiple scales (with pooling sizes of 1, 2, 4, and 5) before applying 3×3 convolutional layers. The resulting multi-resolution embeddings are then concatenated and projected back to a 128-dimensional representation.

The DNA encoder adopts a two-stage architecture. In the first stage, a 1D convolutional encoder processes the one-hot encoded DNA sequence together with the DNA mappability track to generate 1D embeddings. The architecture of this stage is adapted from Orca [29] and consists of seven stacked 1D residual blocks with kernel sizes of 9 and channel dimensions of 64, 96, 128, 128, 128, 128, and 128, respectively. Each residual block is preceded by a max-pooling layer with pooling sizes of 1, 4, 4, 5, 5, 5, and 1, progressively downsampling the input sequence to produce embeddings at an effective resolution of 2 kb. In the second stage of the DNA encoder, the 1D embeddings corresponding to the row and column indices of the Hi-C matrix are combined by summation and passed through a multi-resolution module identical to that used in the Hi-C encoder. The resulting representations are then projected back into a 128-dimensional embedding space.

### Evo 2 embeddings

To reduce computational cost, we precomputed Evo 2 embeddings on reference genomes (hg38 for training and evaluation, and mm10 for evaluation only). Specifically, we performed inference using the official Evo 2-7B checkpoint, extracted embeddings from layer 27, and averaged them within non-overlapping 2 kb bins, following the protocol described in the Evo 2 paper. To further improve efficiency, inference was conducted on 60 kb subsequences with a 50 kb stride along the genome. Because Evo 2 was designed with a causal mask (i.e., each token can only see tokens before it), we discard the initial 10 kb of each subsequence to avoid edge effects. Inference was performed on both DNA strands, and the resulting strand-specific embeddings were concatenated to obtain the final Evo 2 representations of dimension 8192.

### Hi-C data

For pretraining, we collected high read-depth Hi-C contact maps from a wide range of sources. The dataset comprises 149 Hi-C maps curated from ENCODE [41] and 4DN [48] (Supplementary Data 1), including experiments based on *in situ* Hi-C, intact Hi-C, and Micro-C protocols. We performed training, validation, and testing splits across species, cell lines, and chromosomes. In summary, the dataset includes 138 human Hi-C maps from 52 cell lines, of which 124 maps from 46 cell lines were used for training and 14 maps from 6 cell lines were held out for testing. Additionally, 11 mouse Hi-C maps from 4 cell lines were used exclusively for testing. For all contact maps, chromosome 8 was reserved for validation, while chromosomes 9 and 10 were left out for testing. The training pool was assembled without any per-cell-line or per-tissue quota, and its composition mainly reflects the availability of public Hi-C data. To characterize this distribution, we summarize the per-cell-line map counts in **Supplementary Table 12**. We further include an analysis quantifying how representation in the pretraining data affects downstream performance gains (**Supplementary Fig. 14**), where no positive correlation is found.

All contact matrices were subject to matrix balancing. For maps stored in .hic format, we applied Square Root Vanilla Coverage (SCALE) normalization [49] as implemented in Juicer; for files in which a SCALE vector was unavailable (certain 4DN-released maps), we fell back to Knight-Ruiz (KR) balancing [50]. For maps stored in .cool format, we used Iterative Correction and Eigenvector decomposition (ICE) balancing [51] via the cooler library [52]. Genomic bins whose normalization weights were non-finite were masked and excluded from all downstream analyses.

To prepare balanced contact counts as model inputs and training targets, we applied a mixed piece-wise normalization: counts below a threshold *d* were linearly scaled, while counts at or above *d* were log-transformed as

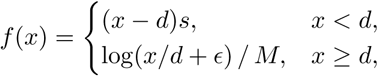

where *d*=10, *ɛ*=10^−5^, *C*_max_=5,000, *M* = log(*C*_max_*/d*), and *s*= log(*d*)*/*((*d*−1)*M* ). Here, *d* defines the transition between the low-count linear branch and the log-compressed branch; *s* is the slope of the low-count branch and approximates the secant slope of the log transform between *x*=1 and *x*=*d*, keeping small contact counts on the same scale as the log transform while avoiding unstable logarithmic behavior near zero; and *C*_max_ sets the upper reference scale for the log-transformed branch. This transformation is designed to compress the range of contact counts and reduce the influence of low-count noise. At inference time, the inverse transform was applied to map model predictions back to raw contact-count space.

### Pretraining

During pretraining, two complementary distillation processes were jointly applied to inject prior knowledge into the encoders. The first process is sequence distillation, where Evo 2 acts as the sequence teacher and the DNA encoder as the student, guiding the encoder to capture sequence-level patterns such as transcription factor motifs. The second process is structure distillation, in which the Hi-C encoder extracts structural knowledge and serves as the structure teacher, transferring knowledge of 3D chromatin organization to help the DNA encoder learn structurally relevant representations.

In sequence distillation, we adopt a contrastive learning framework to inject Evo 2’s knowledge into the DNA encoder. Specifically, embeddings derived from the same genomic sequence by Evo 2 and the DNA encoder are treated as positive pairs, while embeddings from different sequences are treated as negative pairs. This setup encourages the DNA encoder to learn an embedding space consistent with that of Evo 2. At each training step on a single GPU, we randomly sample six 200 kb regions from the human genome training chromosomes. The DNA encoder produces 1D DNA embeddings for every non-overlapping 2 kb bin within these regions, with 20 kb trimmed from both edges to avoid boundary artifacts, yielding 480 embeddings of dimension 128. The corresponding pre-computed Evo 2 embeddings for these 2 kb bins are then retrieved. To enrich the pool of negative samples, we additionally sample 20,480 bins across the genome and obtain their Evo 2 embeddings. This procedure results in 10,060,800 sample pairs in total, of which 480 are positive. To perform the contrastive alignment, additional linear projection layers are appended to both the DNA encoder and Evo 2 embeddings, mapping them into a shared 512-dimensional space. The contrastive distillation loss follows the formulation of SigLIP [53] and is defined as

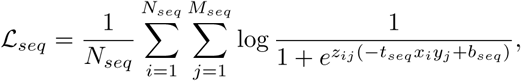

where *N_seq_* and *M_seq_* denote the numbers of DNA encoder and Evo 2 embeddings, respectively; *x_i_* and *y_j_* represent the *i*-th DNA encoder and *j*-th Evo 2 embedding; *z_ij_* equals 1 for positive pairs and −1 for negative pairs; and *t_seq_* and *b_seq_* are learnable temperature and bias parameters that modulate the contrastive scale. In the structure distillation stage, we adopt a similar contrastive learning framework to transfer structural knowledge, such as chromatin loops and topologically associating domains (TADs), from Hi-C contact maps into the DNA encoder. Specifically, for each 2 kb × 2 kb pixel, the Hi-C encoder extracts an embedding from the Hi-C patch centered at that pixel, while the DNA encoder computes a corresponding 2D embedding using the DNA sequences of the row and column bins defining the patch. Within the contrastive framework, Hi-C and 2D DNA embeddings from the same pixel are treated as positive pairs, whereas embeddings from different pixels are treated as negative pairs. At each training step on a single GPU, we randomly sample six 200 kb × 200 kb submatrices at 2 kb resolution from the human Hi-C contact map within a 500 kb diagonal window, excluding the validation and test chromosomes. For each submatrix, eight Hi-C contact maps are randomly selected, and the Hi-C encoder computes embeddings for every pixel across these maps. The embeddings from these eight contact maps are then averaged to obtain the final Hi-C embeddings for pixels. The DNA encoder generates the corresponding 2D DNA embeddings for all pixels, as described in the previous section. To avoid boundary effects, embeddings at both 20 kb borders are discarded. This procedure yields a total of 1,474,560,000 sample pairs, among which 38,400 are positive. Similarly, the contrastive distillation loss follows the formulation of SigLIP and is defined as

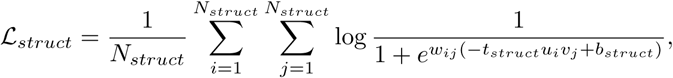

where *N_struct_* denotes the number of Hi-C and 2D DNA embeddings, *u_i_* and *v_j_* represent the *i*-th Hi-C and *j*-th 2D DNA embedding, respectively, *w_ij_* equals 1 for positive pairs and −1 for negative pairs, and *t_stru_* and *b_stru_* are learnable temperature and bias parameters that control the contrastive scaling.

The final pretraining objective is defined as

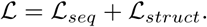

Using this loss, we trained the model on four NVIDIA A100 GPUs with Adam optimizer, an initial learning rate of 10^−4^, and a total of 50,000 training steps, which was determined based on convergence on the validation set.

### Details on measuring embedding space similarity and retrieval

After distilling knowledge from Evo 2 and Hi-C, we evaluated whether the DNA encoder learned an embedding space consistent with Evo 2. To perform this analysis in the original Evo 2 space, we fitted a linear transformation from the 128-dimensional 1D DNA embeddings to the 8192-dimensional Evo 2 embeddings using a least-squares objective on chromosome 9. The learned transformation was then applied to 1D DNA embeddings from chromosome 10 of the human genome (hg38) and chromosome 4 of the mouse genome (mm10). Finally, we quantified the similarity between embedding spaces by comparing pairwise similarities among DNA bins in the Evo 2 space and in our DNA encoder’s transformed space. 5000 pairs of bins were sampled, and the similarity between each pair in the two spaces was computed. The Spearman correlation coefficient (SPC) between these two lists of distances was reported as the final measure of embedding-space similarity.

Next, we assessed whether the encoders successfully captured structural knowledge by evaluating their ability to retrieve Hi-C pixels directly from the corresponding DNA sequences. For computational efficiency, retrieval was restricted to Hi-C pixels located at the same distance from the diagonal within the test chromosomes. We computed the cosine similarity between Hi-C embeddings and 2D DNA embeddings. We reported the top-1, top-5, and top-10 recall (where top-*k* recall indicates that the positive pair appears within the *k* most similar pairs), the average rank of the positive pairs, and the similarity change between positive pairs and negative pairs.

### Predicting Hi-C contact maps

For Hi-C contact map prediction, we used 4 kb–resolution Micro-C data from the H1ESC and HFF cell lines, normalized with SCALE normalization [49]. For both cell lines, chromosome 8 was reserved for validation, and chromosomes 9 and 10 were used for testing. Notably, the H1ESC cell line was entirely held out during pretraining, and the validation and test chromosomes of the HFF cell line were also excluded from the pretraining data.

We designed a U-Net–style architecture [54] as the decoder for this task, which takes 2D DNA embeddings as input and outputs the target Hi-C submatrices. In particular, this U-Net-style architecture is an encoder-decoder architecture with skip connections, where the encoder utilizes three residual blocks to extract features at different granularities, while the decoder employs another three residual blocks to generate the desired output [54]. We fine-tuned Evo2HiC separately for the two cell lines. During fine-tuning, we randomly sampled submatrices of size 1120 kb × 1120 kb along the diagonal of the Hi-C matrix. Each GPU processed a batch of seven submatrices, resulting in a total batch size of 28 across four NVIDIA A100 GPUs. The model was trained using the mean squared error (MSE) loss:

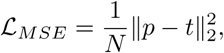

where *p* and *t* denote the predicted and target Hi-C submatrices, respectively, and *N* is the submatrix size. Data augmentation was applied by randomly flipping both the Hi-C submatrices and the corresponding DNA sequences with a probability of 0.5, where flipping refers to reversing the DNA sequences to their reverse-complement strand and mirroring the Hi-C submatrix along both row and column axes. In addition, the DNA sequences were randomly shifted by 100 bp. The model was trained using the Adam optimizer with an initial learning rate of 10^−4^ for a total of 20,000 steps. For evaluation, we applied the model to predict 1 Mb × 1 Mb sliding windows along the diagonal of the Hi-C matrices with a stride of 0.5 Mb. Because the model was trained to predict 1120 kb × 1120 kb submatrices, we trimmed 80 kb from the top and left borders and 40 kb from the bottom and right borders to match the target window size.

We next prepared Orca and Evo 2 as baselines for Hi-C contact map prediction. For Orca, since our evaluation uses the same data sources, we directly downloaded the Orca predictions from the official repository and followed the provided instructions to convert them back into Hi-C contact maps. For Evo 2, we replaced the 2D DNA embeddings with the pre-computed Evo 2 embeddings and used the same decoder architecture and fine-tuning objective to predict Hi-C maps from Evo 2 embeddings.

To evaluate model performance, we followed a widely used scheme based on distance-stratified correlation and insulation score correlation [30]. For distance-stratified correlation, pixels in the Hi-C matrices were grouped by their distance to the diagonal. Specifically, we tiled held-out test chromosomes (chr 9 and chr 10) with 1 Mb windows at 4 kb resolution using a 500 kb step size. For each diagonal offset *d* (*d* = 0, 1*, . . . ,* 249, corresponding to genomic distances 0, 4*, . . . ,* 996 kb), we concatenated all pixels at that offset across the 417 test windows and compute correlation in the raw count space. We report distance-stratified correlation because it isolates each model’s ability to explain contact variation within each genomic-distance bin, rather than summarizing performance across all distances into a single correlation. This per-distance view more directly captures the biological features of interest in this work, such as chromatin loops and TAD boundaries, which manifest at specific distance ranges and can be diluted under a single global metric. For insulation score correlation, we used cooltools [55] to calculate the insulation score for each bin in the submatrices and reported the Pearson correlation coefficient (PCC) and Peak signal-to-noise ratio (PSNR) between the predicted and target insulation profiles.

### Comparison in Orca’s setting

To compare with Orca under its original training target and evaluation protocol, we additionally fine-tuned Evo2HiC in Orca’s setting. The training target was the Micro-C contact matrix of each cell line in Orca’s representation, i.e., ICE-balanced 4 kb contacts processed with cooltools adaptive coarse-graining [55], as in Orca’s preprocessing. As in Orca, the cell-matched Roadmap Epigenomics peak sets (32 tracks for H1ESC and 22 for HFF) were used as auxiliary supervision through a binary cross-entropy loss on the one-dimensional sequence embeddings. The DNA encoder was initialized from the pretrained Evo2HiC encoder, and the auxiliary loss was added to the Hi-C MSE loss with a weight of 0.05. The model was trained with the Adam optimizer at an effective batch size of 40 (bf16 mixed precision, synchronized batch normalization across four GPUs), first at a learning rate of 5 × 10^−4^ after a 1,000-step warm-up, for at least 10,000 steps, and then at 10^−4^ from the best checkpoint; each stage was stopped when the chromosome-8 validation MSE had not improved for five consecutive checkpoints (saved every 1,000 steps). Because batch-normalization running statistics lag behind the weights at a high learning rate, and the resulting validation scores can misrank checkpoints, every checkpoint was evaluated after recomputing its batch-normalization statistics on 50 training batches. The reported model is the checkpoint with the lowest recalibrated chromosome-8 validation MSE (0.68 million training windows for H1ESC and 0.16 million for HFF, compared with approximately 7.7 million for Orca’s 1 Mb model), and the test windows were used only once, for its final evaluation. Both models were evaluated on Orca’s 417 test windows (1 Mb at 4 kb resolution) on chromosomes 9 and 10, against Orca’s target matrices and with Orca’s validity mask. We report Orca’s metric, the Pearson and Spearman correlations of log(observed/expected) contacts pooled over all valid pixels of all windows, together with four complementary metrics [30]: observed/expected correlation, mean squared error and distance-stratified correlation, computed with the published implementation of ref. [30] on log-transformed, distance-normalized maps, and insulation score correlation, computed with cooltools [55] (100 kb window) as in Fig. 3b,f. Recomputing Orca’s released predictions in this pipeline reproduced its published values (H1ESC 0.776/0.761, HFF 0.737/0.716 for PCC/SPC, against the published 0.776/0.762 and 0.740/0.718). Differences between models were tested with paired tests across windows.

### Predicting epigenomic profiles

For epigenomic signal prediction, we collected DNase-seq, ChIP-seq CTCF, ChIP-seq H3K27ac, ChIP-seq H3K27me3, and ChIP-seq H3K4me3 data for the GM12878, H1ESC, and K562 cell lines from ENCODE (accession codes in **Supplementary Table 1**). As in pretraining, chromosome 8 was reserved for validation, and chromosomes 9 and 10 were used for testing. The entire K562 cell line was additionally held out for testing to assess the model’s ability to generalize to unseen cell types. All epigenomic signals were binned at 2 kb resolution. To normalize signal scales across cell lines, we computed the 90th percentile value (*q*) of the signal distribution and used 20 × *q* as a cutoff threshold, normalizing signal intensities to the range [0,1]. We confirmed that in every cell line, fewer than 1% of bins were truncated by this procedure, indicating that nearly all information was preserved.

We next designed a transformer-based decoder to predict these epigenomic signals from multimodal embeddings. The decoder takes 1D DNA embeddings and Hi-C embeddings as input to predict epigenomic signals. Specifically, the Hi-C embeddings were computed with the Hi-C encoder for a 320 kb × 2 Mb region centered along the diagonal and then averaged over the 2 Mb dimension to obtain 1D representations. These 1D Hi-C embeddings were concatenated with the 1D DNA embeddings to form the input to the transformer decoder. The decoder consists of 11 transformer encoder layers with a base hidden dimension of 256 and employs Rotary Position Embedding (RoPE) [56] to encode positional information.

For the finetuning process, the model predicts epigenomic signals within a 320 kb window, using as input a 320 kb DNA sequence together with a 320 kb × 2 Mb Hi-C submatrix centered along the diagonal. The fine-tuning loss combines cosine similarity and mean squared error (MSE), defined as

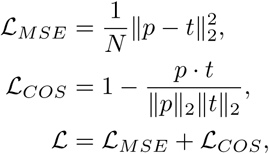

where *p* and *t* denote the prediction and target, respectively, and *N* is the size of target sample. The model was fine-tuned with a total batch size of 16 across four GPUs using the Adam optimizer, an initial learning rate of 10^−4^, and a total of 50,000 steps.

We evaluated Evo2HiC against several baseline models. The Hi-C–only model uses the same architecture as Evo2HiC but excludes DNA sequences from the input. The Evo 2–only model employs the same decoder but replaces DNA sequences with pre-computed Evo 2 embeddings and removes Hi-C input. Because the Evo 2–only model lacks cell line–specific structural information, we trained separate models for each cell line, including the leave-out cell line. A no-distillation Evo2HiC ablation as well as a Hi-C ridge regression model are also included in supplementary experiments. Model performance was evaluated using Pearson and Spearman correlation coefficients (PCC and SPC) between the predicted and target epigenomic signals over the test chromosomes.

For the cross-cell type analysis, uncertainty of the per-assay improvement over the strongest baseline was estimated with a paired block bootstrap over genomic bins (100 kb blocks, 2,000 resamples), averaged over the three pairs of cell lines.

We further compared Evo2HiC with HiCFoundation [33], using its released epigenomic prediction weights without retraining, on the same raw cell-line Hi-C maps (1 kb input) and evaluated with the same code on the same test chromosomes. HiCFoundation holds out chromosomes 4, 5, 11 and 14 rather than 9 and 10, so our test chromosomes were part of its training, which makes the comparison conservative for Evo2HiC.

To locate putative enhancers for a gene of interest, we computed the Activity-by-Contact (ABC) score following Fulco et al. [34]. For a candidate element *i* and the target gene promoter *j*, the score is defined as

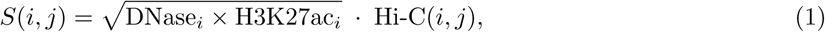

where the activity term is the geometric mean of the DNase accessibility and H3K27ac signal at element *i*, weighted by the Hi-C contact frequency between *i* and *j*. Candidate elements were then ranked by their ABC scores (unnormalized), and high-scoring elements were nominated as putative enhancers of the target gene. For evaluation, we used two genome-wide CRISPRi screens in K562, Gasperini et al. [35] and Morris et al. [36], restricted to the held-out chromosomes 9 and 10. For Gasperini et al., we used all tested pairs from the published results, lifted over to hg38; a pair was positive if expression decreased at empirical FDR *<* 0.1, and pairs tested by several gRNA groups were merged. For Morris et al., we used the pairs as uniformly processed by ENCODE-rE2G [57]. We scored all element–gene pairs 6 kb to 1 Mb from the transcription start site (5,786 pairs, 44 positives, 808 genes; 5,462 distal pairs with 18 positives), using 10 kb contact bins from the measured K562 Hi-C map without a power-law pseudocount. This protocol was selected using ABC computed from measured tracks only, without reference to our model. To place predicted signals on the scale of measured signals, the predicted DNase and H3K27ac tracks were quantile-normalized to the genome-wide distribution of the corresponding measured tracks in the training cell line GM12878 before computing ABC scores. Genomic distance (1*/d*), Hi-C contact alone and ABC computed from the predictions of the Hi-C–only ablation served as baselines. We report AUPRC for distal pairs (≥ 50 kb) and the fraction of positives recovered among the top *K* distal pairs ranked across all genes. Differences in AUPRC were tested with a one-sided paired permutation test in which the two methods’ rank-transformed scores were swapped within each gene with probability 0.5 (10,000 permutations).

For loop anchor classification, we assigned a chromatin state to each anchor based on its local epigenomic signals. For each anchor, we considered a ±5 kb window centered on the anchor and, for each assay, took the maximum signal over this window as the anchor’s representative value. An assay was called *positive* at an anchor if this value exceeded the 90th percentile of the corresponding genome-wide signal distribution. Anchors were then categorized using a priority-based rule. An anchor positive for H3K27me3 was labeled *repressive*. An anchor that was not repressive but positive for H3K4me3 or H3K27ac was labeled *active*. Anchors satisfying neither criterion were labeled *quiescent*. Independently of this assignment, any anchor positive for CTCF was additionally annotated as *CTCF (structural)*, allowing an anchor to carry both a regulatory state and a structural label.

For the held-out cell line K562, loops were called with HiCCUPS using the same parameters on chromosomes 9 and 10 (415 loops). As a reference classification independent of our own rule, we additionally assigned anchors to the classes of the ENCODE registry of candidate cis-regulatory elements [37]: an anchor was a candidate element if DNase and at least one of H3K4me3, H3K27ac and CTCF were high (above the 95th percentile of the genome-wide 2 kb-bin distribution, corresponding to the registry’s *Z >* 1.64), and was then classified as promoter-like (high H3K4me3), enhancer-like (high H3K27ac, low H3K4me3), CTCF-only (high CTCF only) or low-DNase. Because our unit is a 2 kb bin, the registry’s transcription start site distance partitions were collapsed. Labels derived from each method’s predicted tracks were compared with labels derived from measured tracks using accuracy, balanced accuracy and Cohen’s *κ*, with 95% confidence intervals from 1,000 bootstrap resamples and chance levels from label shuffling. The Hi-C–only and sequence-only (Evo 2) models described above served as baselines. Two readouts independent of measured epigenomic data were computed from predicted tracks alone. First, CTCF motifs were identified in anchor sequences with JASPAR MA0139.2 [38] at *p <* 10^−5^ [39]; we report the AUROC of predicted CTCF signal for separating motif-bearing from motif-free anchors, and the fraction of loops with convergent motif orientation among loops whose two anchors were both predicted CTCF-bound. Second, genes whose transcription start site fell within an anchor were assigned ENCODE RNA-seq expression values, and we report the AUROC for separating genes at predicted promoter-like anchors from genes at predicted CTCF-only or low-DNase anchors.

### Identifying cell type-specific motifs

We identified cell type–specific motifs by attributing epigenomic signal variation to individual base pairs. Specifically, we used the DeepLiftShap [40] algorithm implemented in tangermeme [58] to compute attribution scores for each base pair in the DNA sequence, conditioned on the Hi-C contact map of a given cell type. DeepLiftShap was applied to every 320 kb window along the test chromosomes with a stride of 240 kb, excluding windows containing unknown bases. We next obtained a precomputed, genome-wide catalog of motif matches [59]. For each motif, we assessed whether its matching sites were associated with higher attribution scores compared to the local background. A high-attribution site was defined as one whose average attribution score exceeded the chromosome-wide mean by at least five standard deviations. The local background was defined as the flanking region extending ten times the motif length on each side. We then quantified the fold enrichment in the proportion of motif sites with high attribution scores relative to their local background and computed the corresponding p-values using Welch’s z-test. To control for multiple testing, we applied the Benjamini–Hochberg false discovery rate (BH-FDR) procedure to all motifs within each cell type. Motifs enriched in a cell type were defined as those with at least 80 high-attribution sites and a BH-FDR–adjusted p-value below 10^−3^. Cell-type–specific motifs were further defined as those enriched in one cell type but not in any other. We additionally identified cell type–specific motif clusters based on a previously reported motif-similarity analysis [59]. Specifically, cell type-specific motif clusters are clusters that have at least one enriched motif in a cell but no enriched motif in another. Finally, the cell-type–specific motifs and motif clusters were cross-validated using RNA-seq data from ENCODE (accession codes in **Supplementary Table 2**), where transcript-per-million (TPM) values served as an external indicator of motif activity across cell types. For motif clusters, the TPMs of all enriched motifs in the cluster are summed as the TPM for the cluster.

### Resolution enhancement across different species

For resolution enhancement, Evo2HiC takes DNA sequences and low-coverage Hi-C matrices as input to reconstruct high-coverage Hi-C contact maps. The model adopts a U-net–style decoder, similar to the one used for predicting Hi-C contact maps. We fine-tuned Evo2HiC on our pretraining dataset using the same data split, simulating low-coverage inputs through binomial downsampling with a ratio of 1/16. The training set includes only human cell lines. To better leverage DNA sequence information, we trained the model on Hi-C matrices at 2 kb resolution. In addition, we introduced a multi-resolution mean squared error (MSE) loss to mitigate sparsity at this fine resolution, defined as

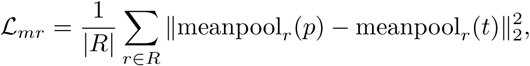

where *p* and *t* denote the predicted and target Hi-C contact maps, respectively; *R* denotes the set of resolutions, and in practice *R* = {2 kb, 4 kb, 8 kb, 10 kb}; meanpool*_r_*(·) denotes average pooling to resolution *r*. We fine-tuned the model with a total batch size of 128 using the Adam optimizer, an initial learning rate of 10^−4^, and 200,000 optimization steps.

We first evaluated model performance on the held-out test cell lines from the pretraining dataset, which include six human and four mouse cell lines. Four metrics were used for evaluation: Peak signal-to-noise ratio (PSNR), Structural similarity index measure (SSIM), and distance-stratified Pearson and Spearman correlation coefficients (PCC and SPC). The distance-stratified PCC and SPC are computed within distance-specific groups of bins and averaged across groups. Following prior work [60], all metrics were computed at 10 kb resolution within a 2 Mb genomic distance. To further assess Evo2HiC’s cross-species generalization, we obtained Hi-C contact maps and DNA sequences from the DNA Zoo dataset [61]. DNA sequences and Hi-C matrices were paired following the DNA Zoo protocol, resulting in 177 sequence–contact map pairs. Using the downsampled low-coverage Hi-C maps and corresponding DNA sequences, we reconstructed the original high-resolution maps and reported distance-stratified SPC and PCC as the evaluation metrics.

As baselines, we included HiCNN [62], HiCARN1, and HiCARN2 [60], following their original training procedures but replacing their datasets with our pretraining data. We further included HiCFoundation [33] using its released resolution enhancement weights without retraining, applied to the same 1/16-downsampled contact maps at 10 kb resolution and evaluated with the same code. HiCFoundation’s fine-tuning data share our biosample split (our 25 test maps are in its test split), but it holds out chromosomes 4, 5, 11 and 14 rather than 9 and 10; on our test chromosomes, HiCFoundation is therefore evaluated on chromosomes seen during its training, which is conservative for Evo2HiC. The Hi-C–only ablation of Evo2HiC uses the same architecture and training procedure without the DNA input.

### Cross-species boundary analysis

TAD boundaries were called at 10 kb resolution on the evaluated scaffold of each species (the first chromosome-scale scaffold of the DNA Zoo assembly) with cooltools [55], using insulation scores with a 100 kb window and retaining boundaries with prominence ≥ 0.2; the same protocol was applied to the ground truth and to every method. TAD-boundary F1 counts a predicted boundary as correct if it lies within three bins of a ground-truth boundary. Boundary precision is the fraction of called boundaries within three bins of a ground-truth boundary, and recall the fraction of ground-truth boundaries with a called boundary within three bins. To isolate the contribution of the sequence input with the model held fixed, we ran the same Evo2HiC checkpoint with every non-N base of the scaffold replaced by an independent random base drawn from the scaffold’s own base composition, keeping the Hi-C input unchanged. For the paired boundary analysis, boundaries called on the Evo2HiC prediction and on a no-DNA reference (random DNA or the Hi-C–only ablation) were matched within one bin and split into boundaries called only with DNA (added) and only without DNA (removed). CTCF motifs were identified on both strands of the evaluated scaffold with the JASPAR MA0139.2 position weight matrix [38] at *p <* 10^−5^, and a boundary was counted as motif-bearing if a match lay within one bin. Sequencing depth was measured as intra-scaffold contacts per 10 kb bin of the ground-truth map, and evolutionary distance as the patristic distance to human in the TimeTree phylogeny; the 23 species without a dated leaf were excluded only from analyses involving divergence time. Paired comparisons across species used two-sided Wilcoxon signed-rank tests, and associations used Spearman correlation. Absolute TAD-boundary F1 values depend on the boundary-calling protocol, matching tolerance and chromosome set, and are therefore not directly comparable with those reported in the HiCFoundation study. Per-species values are provided in Supplementary Data 2.

## Data Availability

All data used in this study are publicly available. Human and mouse reference genomes (hg38 and mm10) were obtained from the UCSC Genome Browser (https://genome.ucsc.edu/). The 149 Hi-C and Micro-C maps used for pretraining and evaluation were obtained from the ENCODE [41] (https://www.encodeproject.org/) and 4D Nucleome [48] (https://data.4dnucleome.org/) data portals; their accession codes, biosources and data splits are listed in Supplementary Data 1. For the comparison in Orca’s setting, Orca’s Micro-C maps were obtained from the 4D Nucleome data portal (4DNFI9GMP2J8 and 4DNFI643OYP9), and its test windows, released predictions and Roadmap Epigenomics peak sets from the Orca repository [29] (https://github.com/jzhoulab/orca). Epigenomic and RNA-seq datasets are listed with their accession codes in Supplementary Tables 1 and 2. Motif annotations were obtained from JASPAR [38] (https://jaspar.elixir.no/) and the motif clustering of Vierstra et al. [59] (https://github.com/vierstralab/motif-clustering). The CRISPRi screens of Gasperini et al. [35] were obtained from the Gene Expression Omnibus (GSE120861), and those of Morris et al. [36] as processed by ENCODE-rE2G [57]. Cross-species Hi-C maps and genome assemblies were obtained from DNA Zoo [61] (https://www.dnazoo.org/), and divergence times from TimeTree (https://timetree.org/). Per-species results are provided as Supplementary Data 2.

## Code Availability

Evo2HiC is accessible at https://github.com/CHNFTQ/Evo2HiC, including the model weights and relevant source code, released under the Apache License 2.0. The tutorial for reproducing figures in this paper is acces-sible at https://github.com/SocraLee/Evo2HiC-Figures. We include detailed methods and implementation steps in the Methods and Supplementary Information to allow for independent replication.

## Acknowledgement

Figure 1 was created using icons adapted from “Various Illustration Vectors” by soco-st, licensed under CC BY 4.0.

## Author Contributions Statement

|  | TF | YL | XW | ZX | SH | GM | JY | HX | AJ | WSN | SW |
| --- | --- | --- | --- | --- | --- | --- | --- | --- | --- | --- | --- |
| Conceptualization | • | • | • |  |  |  |  |  |  | • | • |
| Data curation | • | • |  |  |  |  |  |  |  |  |  |
| Funding acquisition |  |  |  |  |  |  |  |  |  | • | • |
| Investigation | • | • |  |  |  |  |  |  |  |  |  |
| Methodology | • | • | • | • | • | • | • | • | • | • | • |
| Project administration | • | • | • |  |  |  |  |  |  | • | • |
| Software | • | • | • |  |  |  |  |  |  |  |  |
| Resources | • |  |  |  |  |  |  |  |  | • | • |
| Supervision |  |  |  |  |  |  |  |  | • | • |  |
| Visualization | • | • | • | • |  |  |  | • |  |  |  |
| Writing—original draft | • | • |  |  |  |  |  |  |  |  | • |
| Writing—review & editing | • | • | • | • | • | • | • | • | • | • | • |

## Extended Data

**Extended Data Fig. 1:**
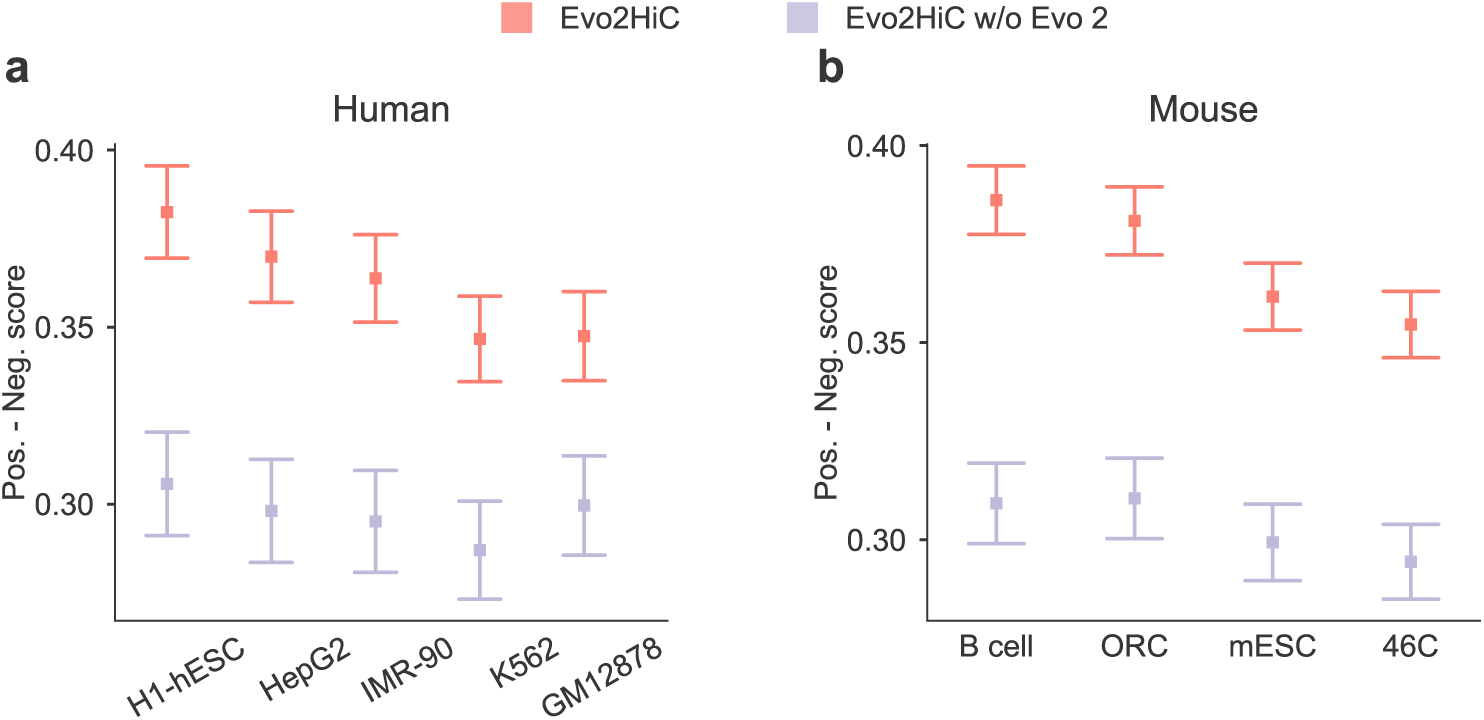
Evaluation of the embedding space of Evo2HiC. **a,b** Comparison between Evo2HiC and the Evo2HiC without Evo 2 on retrieving the most similar Hi-C patch using the corresponding genomic sequence on human **(a)** and mouse **(b)**. y-axis is the gap between positive pairs and negative pairs.

**Extended Data Fig. 2:**
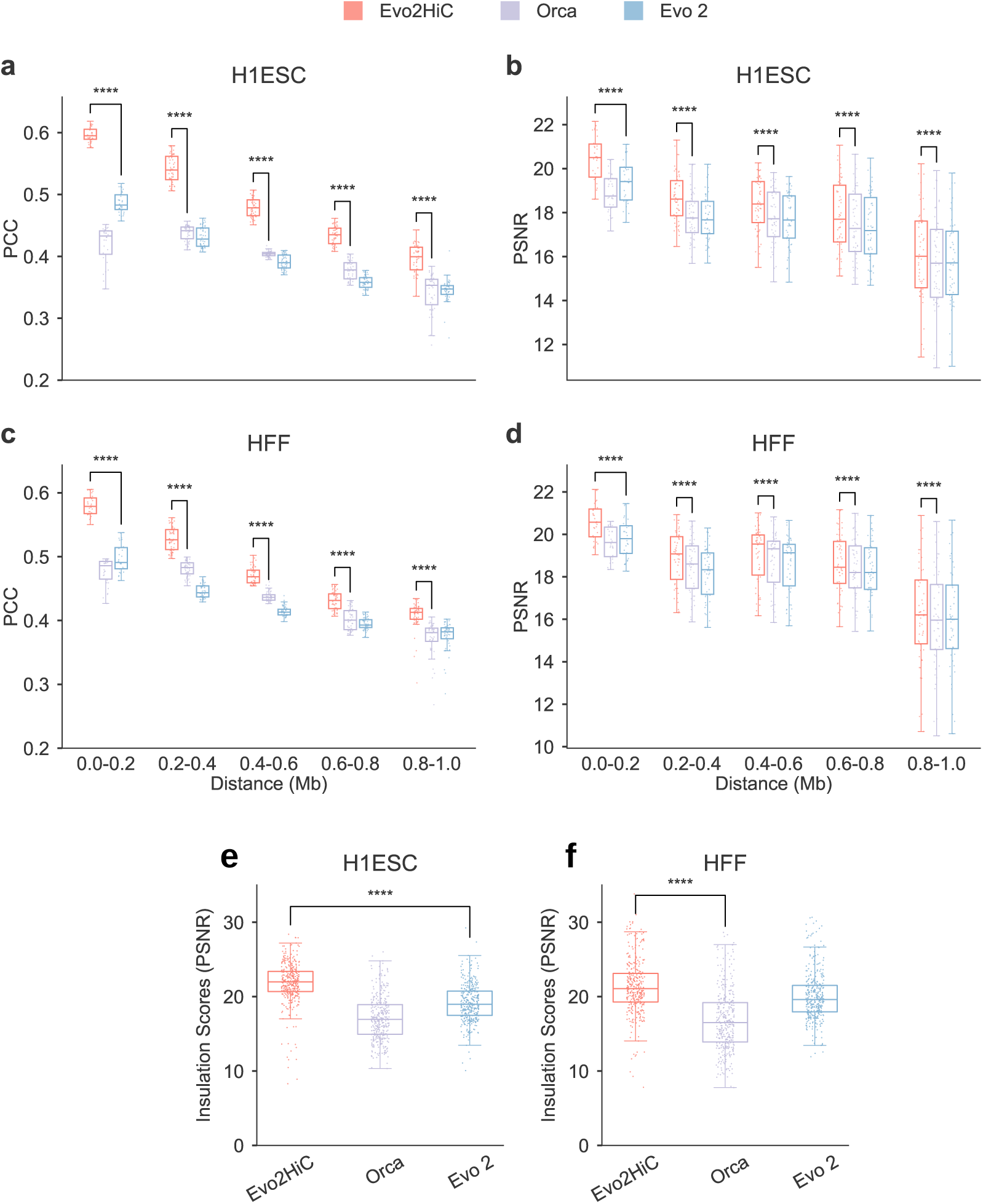
Genome structure prediction from sequence: distance-stratified metrics and insulation score. **a–d**, Comparison among Evo2HiC, Orca, and Evo 2 for predicting Hi-C contact maps from genomic sequence input. Metrics are Pearson correlation (PCC) **(a, c)** and Peak signal-to-noise ratio (PSNR) on Hi-C contact counts **(b, d)**. **a,b** are results on the H1ESC cell line and **c,d** are results on the HFF cell line. **e,f,** Peak signal-to-noise ratio (PSNR) of the insulation score on the H1ESC cell line (**e**) and the HFF cell line (**f** ).

**Extended Data Fig. 3:**
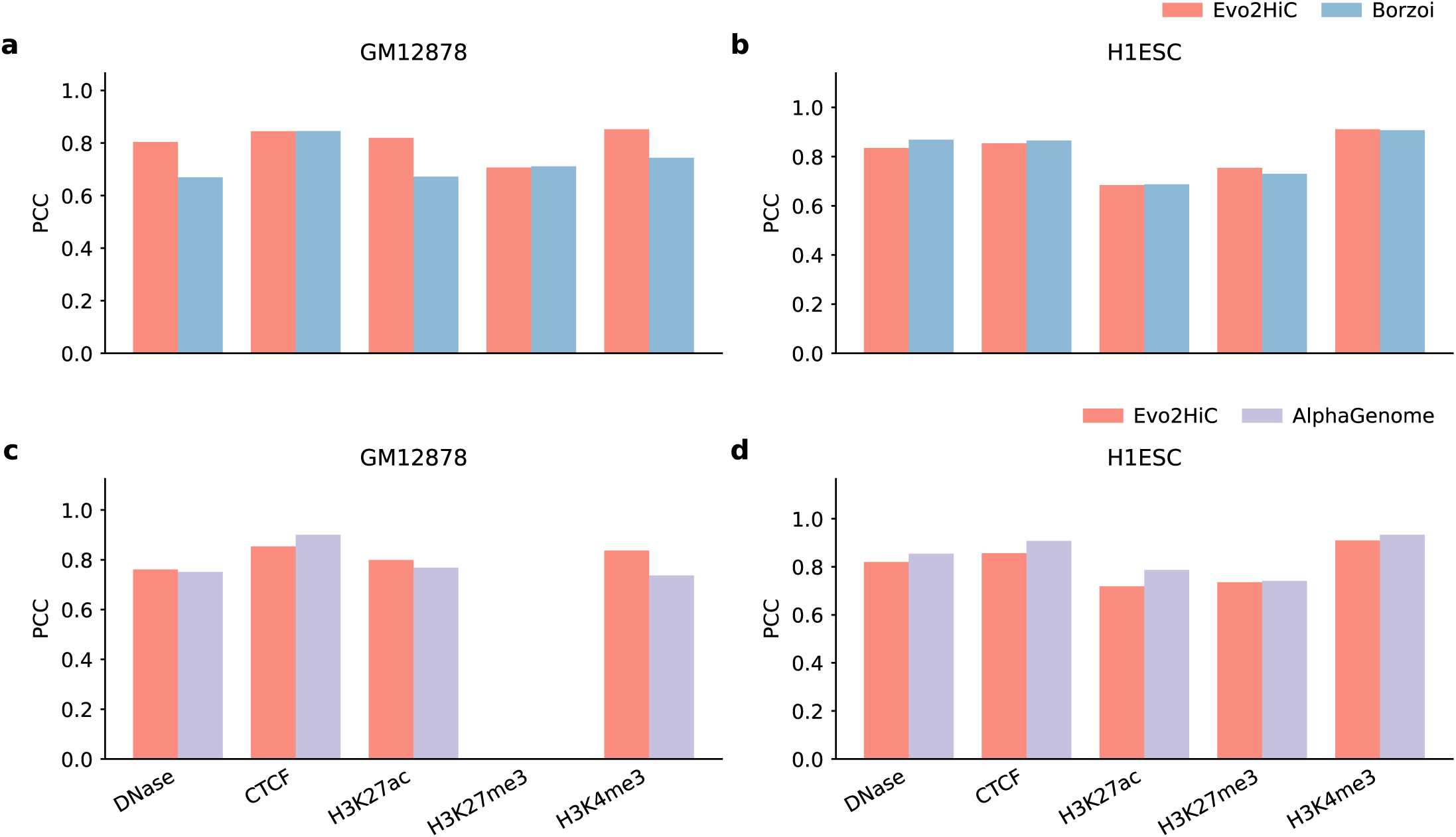
Evaluation of epigenomic signal predictions against Borzoi. **a–d**, Comparison of Evo2HiC against Borzoi on held-out genomic bins. Evo2HiC improves PCC by 5.07% (**a, b**) and SPC by 6.74% (**c, d**) on average. Evaluation is restricted to intersection of Evo2HiC and Borzoi’s test fold with no training/validation overlap. K562 was excluded as it is in the training set for Borzoi but held-out for Evo2HiC. Statistical significance was determined using per-chromosome Williams’ tests combined via Fisher’s method.

**Extended Data Fig. 4:**
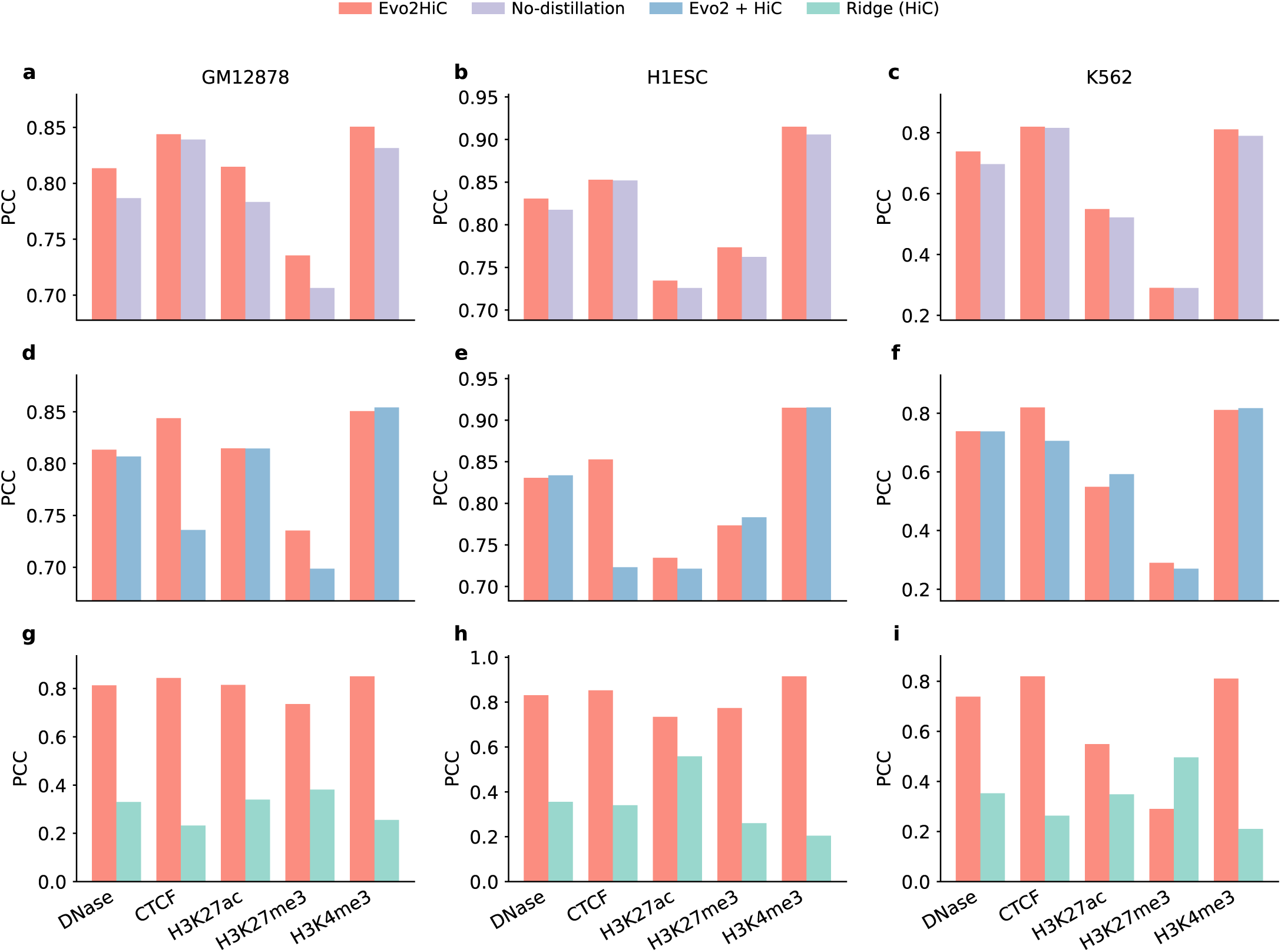
Evaluation on predicting epigenomic signals. **a–c**, Against a no-distillation variant, Evo2HiC achieves an average absolute PCC improvement of 2.2%, 0.9%, and 1.9% on GM12878 (a), H1ESC (b), and K562 (c), respectively. **d–f**, Against an Evo2 + HiC variant, Evo2HiC yields an average absolute PCC improvement of 3.0%, 2.6%, and 1.7% respectively. **g–i**, Against the HiC-based Ridge regression baseline, Evo2HiC yields an average absolute PCC improvement of 50.4%, 47.8%, and 30.8%, respectively. All improvements are statistically significant (*p <* 1 *×* 10*^−^*^4^) under Williams’ test combined via Fisher’s method.

**Extended Data Fig. 5:**
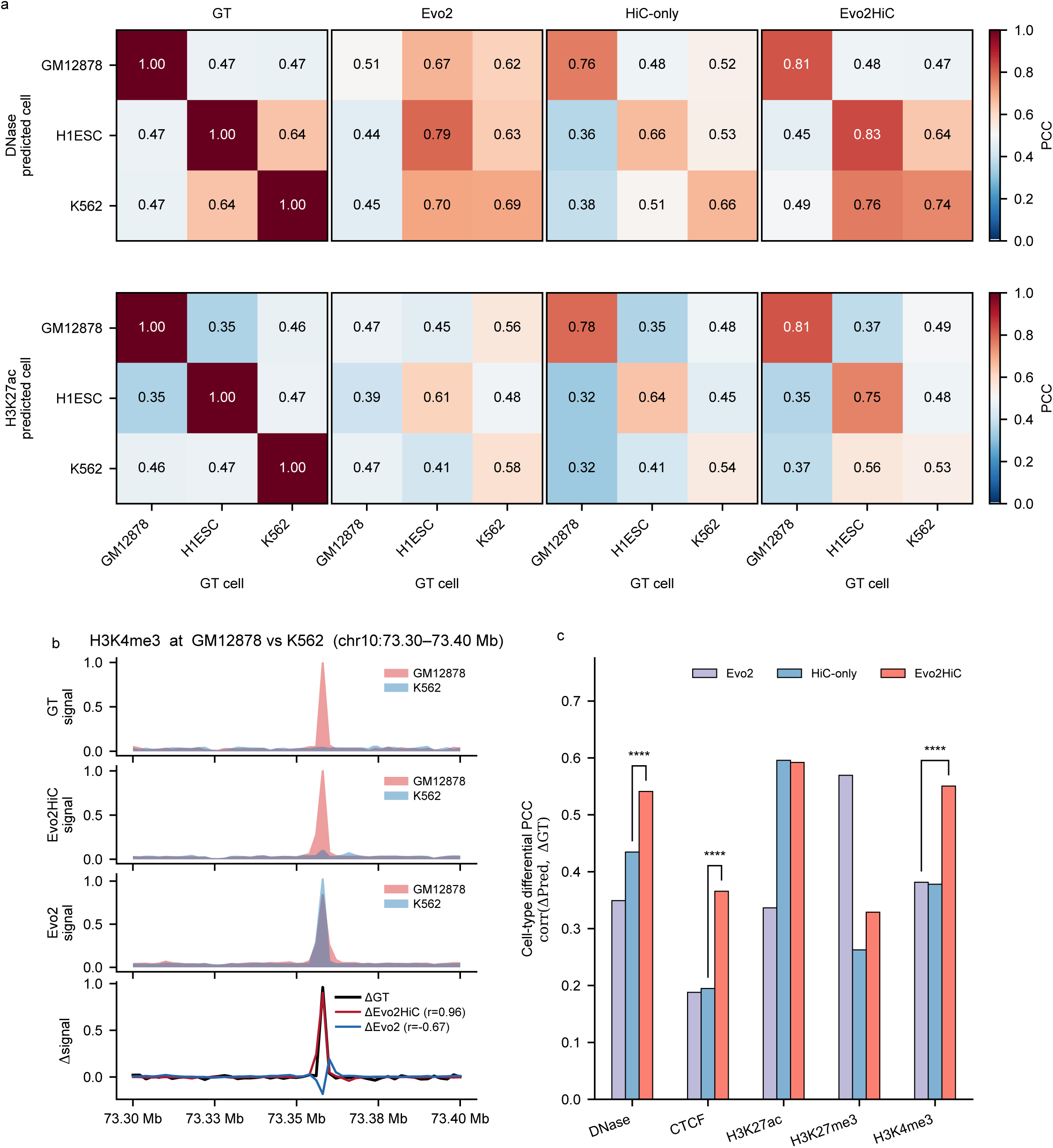
Cross-cell-type evaluation. **a**, Cross-cell-type Pearson correlation matrices comparing predicted and experimental tracks across three cell lines for representative tracks. Evo2HiC successfully restores the cell-type-specific diagonal patterns observed in the experimental ground truth (GT). **b**, A representative genomic locus illustrating the cell-type differential signal (Δ). While Evo2 produces cell-agnostic predictions, Evo2HiC accurately captures the cell-type-specific peak imbalance. **c**, Cell-type differential analysis. Correlation between predicted and experimental between-cell differences (correlation between ΔPred and ΔGT) further supports the cross-cell-type

**Extended Data Fig. 6:**
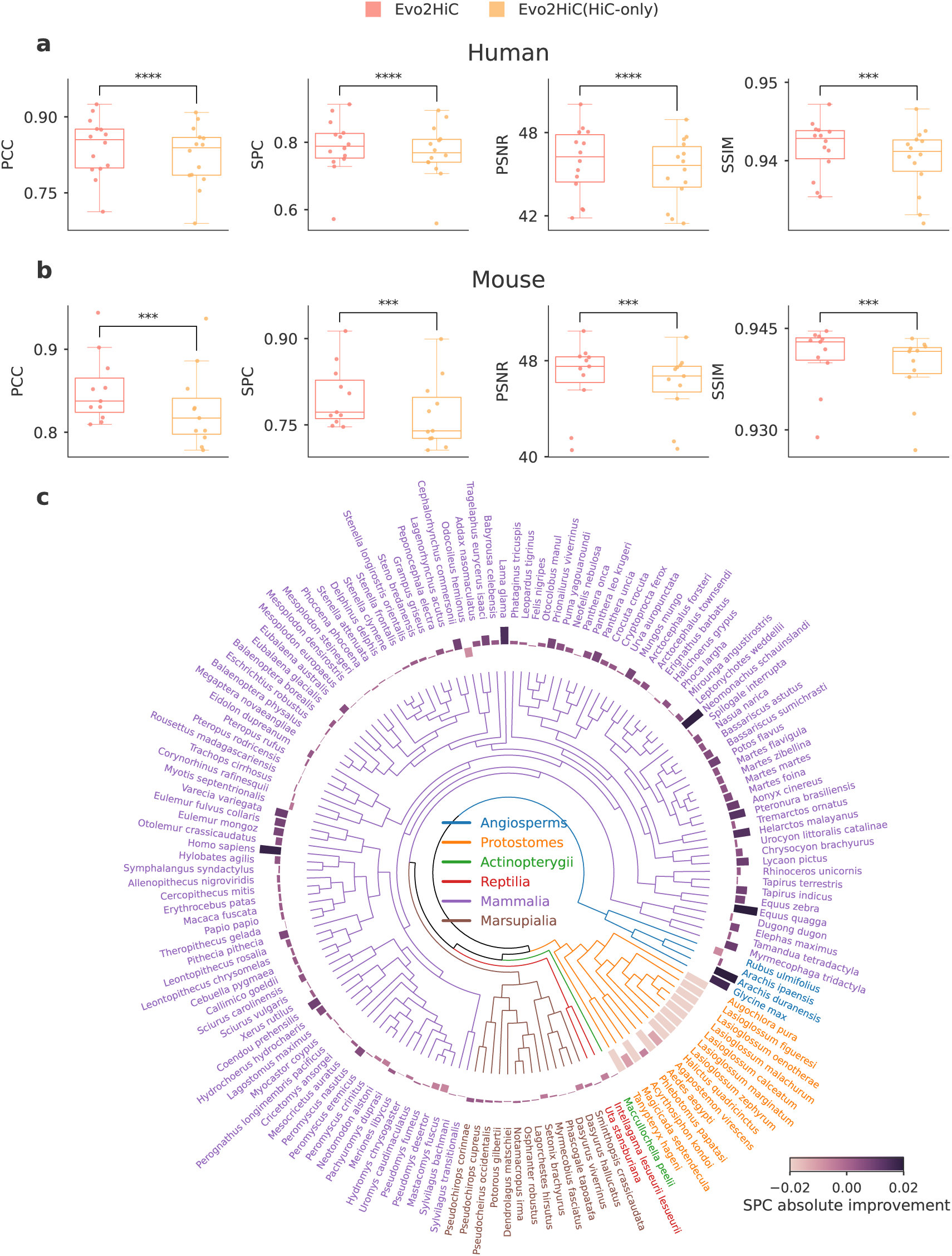
Ablation study against to HiC-only variant. **a, b**, Integrating the sequence prior improves all evaluation metrics over a HiC-only ablation on held-out human (**a**) and mouse (**b**) chromosomes. **c**, Across 177 evaluated species, the sequence prior improves predictions in 116 species. Specifically, the performance gain exhibits a clear phylogenetic decay: it remains robust in closely related mammals (80% improved) but attenuates in evolutionarily distant lineages, reflecting the bias of human-centric training distribution.

**Extended Data Fig. 7:**
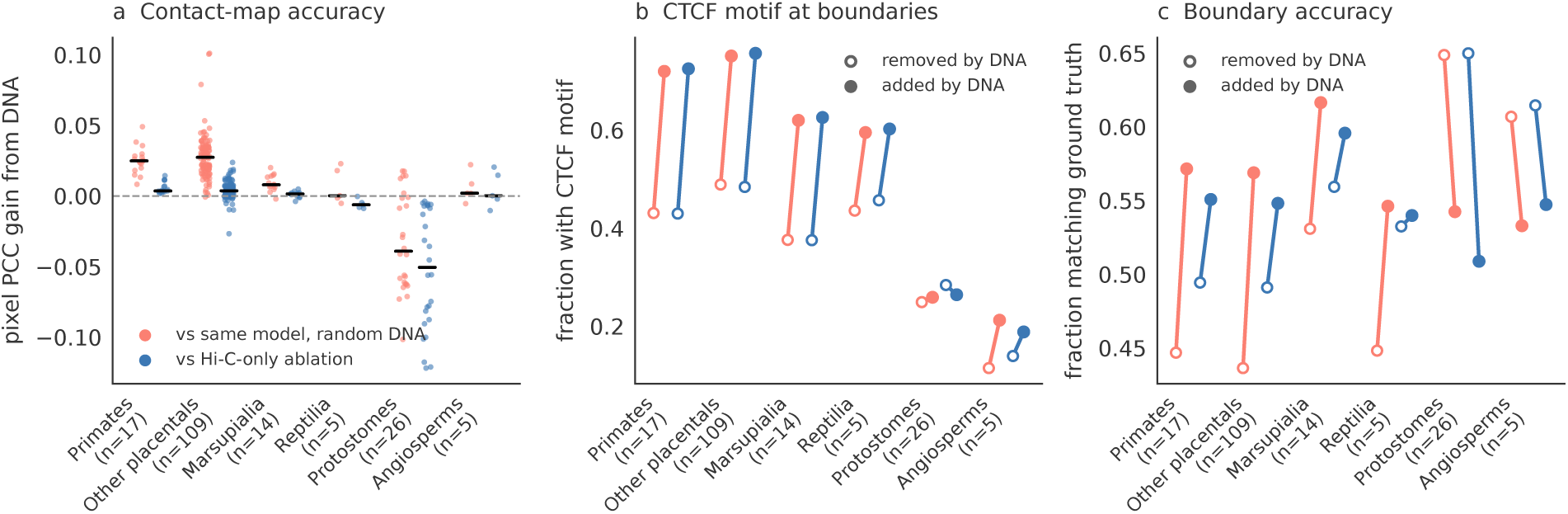
Contribution of the sequence prior across species, paired DNA versus no-DNA comparisons. Each species is compared with two no-DNA references: the same Evo2HiC checkpoint run on random DNA of matched base composition (red), and the Hi-C–only ablation (blue). **a**, Gain in pixel-level PCC from the sequence input, per species (lines: medians). **b**, Fraction of TAD boundaries (paper protocol) that carry a CTCF motif, for boundaries called only with DNA (added, filled) versus only without DNA (removed, open); means over species. **c**, Fraction of added and removed boundaries within three bins of a ground-truth boundary. Placental mammals are split into primates and other placentals; the single fish species is omitted from the plot. Values, counts and paired tests are given in **Supplementary Tables 15 and 16**.

**Extended Data Table 1:** Comparison with Orca in Orca’s setting. Evo2HiC trained on Orca’s Micro-C target representation with Orca’s auxiliary epigenomic peak sets, evaluated with Orca’s released predictions on its 417 test windows (chromosomes 9 and 10, 1 Mb at 4 kb), against Orca’s target matrices and with Orca’s validity mask. Orca’s reported metric is the pooled correlation of log(observed/expected) contacts; published values are 0.776/0.762 (H1ESC) and 0.740/0.718 (HFF). The remaining rows are complementary metrics [30] (observed/expected correlation, mean squared error and distance-stratified correlation computed with the published implementation; insulation score correlation computed with cooltools, 100 kb window) and TAD-boundary metrics. *p*: one-sided paired Wilcoxon signed-rank test across windows (Evo2HiC better than Orca). The reported Evo2HiC checkpoints were selected on batch-normalization-recalibrated chromosome-8 validation MSE; the test windows were used once.

| Metric | H1ESC |  |  | HFF |  |  |
| --- | --- | --- | --- | --- | --- | --- |
| | Evo2HiC | Orca | $p$ | Evo2HiC | Orca | $p$ |
| Pooled PCC, log(O/E) (Orca’s metric) | 0.7701 | 0.7759 | – | 0.7289 | 0.7367 | – |
| Pooled SPC, log(O/E) (Orca’s metric) | 0.7530 | 0.7613 | – | 0.7108 | 0.7159 | – |
| Insulation score PCC | 0.9397 | 0.8873 | $6.5 \times 10^{-50}$ | 0.9168 | 0.8543 | $5.3 \times 10^{-46}$ |
| Observed/expected PCC | 0.6420 | 0.4797 | $8.1 \times 10^{-59}$ | 0.4559 | 0.4204 | $3.4 \times 10^{-13}$ |
| Mean squared error (lower better) | 0.0047 | 0.0066 | $3.2 \times 10^{-38}$ | 0.00086 | 0.00088 | $3.5 \times 10^{-4}$ |
| Distance-stratified PCC | 0.5300 | 0.5129 | $8.8 \times 10^{-6}$ | 0.4448 | 0.4382 | 0.18 |
| TAD-boundary $F_1$ | 0.7285 | 0.7157 | 0.14 | 0.6938 | 0.6518 | $3.3 \times 10^{-4}$ |
| Boundary PR-AUC | 0.7815 | 0.7618 | 0.052 | 0.7269 | 0.6921 | 0.0038 |

**Extended Data Table 2:** Comparison with HiCFoundation. a. , Epigenomic prediction. PCC between predicted and measured signals, averaged over chromosomes 9 and 10. HiCFoundation uses its released epigenomic weights; both models take the same Hi-C maps as input. K562 is held out for both models. **b**, Hi-C resolution enhancement. Mean over maps (human, mouse) or species (DNA Zoo) of distance-stratified PCC and SPC, SSIM and PSNR at 10 kb within 2 Mb. Wins: number of maps or species in which Evo2HiC is better than HiCFoundation; *p*: one-sided paired Wilcoxon signed-rank test in the direction of the better method. HiCFoundation is evaluated on chromosomes included in its training (Methods).

**a**
| Cell line | Method | DNase | CTCF | H3K27ac | H3K27me3 | H3K4me3 |
| --- | --- | --- | --- | --- | --- | --- |
| GM12878 | Evo2HiC | **0.814** | **0.844** | **0.815** | **0.735** | **0.851** |
|  | HiCFoundation | 0.679 | 0.548 | 0.743 | 0.458 | 0.625 |
| H1ESC | Evo2HiC | **0.831** | **0.853** | **0.734** | **0.774** | **0.915** |
|  | HiCFoundation | 0.563 | 0.516 | 0.391 | 0.512 | 0.671 |
| K562 | Evo2HiC | **0.738** | **0.820** | **0.549** | 0.290 | **0.811** |
|  | HiCFoundation | 0.539 | 0.418 | 0.524 | **0.452** | 0.534 |

**b**
| Dataset | Metric | Evo2HiC | HiCFoundation | HiCARN2 | Wins | $p$ |
| --- | --- | --- | --- | --- | --- | --- |
| Human (14 maps) | PCC | 0.840 | 0.841 | 0.811 | 5/14 | 0.12 |
|  | SPC | 0.788 | 0.784 | 0.755 | 11/14 | 0.015 |
| | SSIM | 0.942 | 0.940 | 0.939 | 14/14 | $6.1 \times 10^{-5}$ |
| | PSNR | 45.90 | 45.04 | 44.60 | 14/14 | $6.1 \times 10^{-5}$ |
| Mouse (11 maps) | PCC | 0.852 | 0.855 | 0.817 | 0/11 | $4.9 \times 10^{-4}$ |
|  | SPC | 0.799 | 0.799 | 0.756 | 6/11 | 0.45 |
|  | SSIM | 0.941 | 0.939 | 0.937 | 10/11 | 0.0015 |
|  | PSNR | 46.65 | 46.26 | 45.50 | 10/11 | 0.0049 |
| DNA Zoo (177 species) | PCC | 0.372 | 0.377 | 0.356 | 127/177 | $1.3 \times 10^{-5}$ |
|  | SPC | 0.324 | 0.329 | 0.315 | 113/177 | 0.0037 |

**Extended Data Table 3:** Enhancer nomination and loop-anchor classification in unseen cellular contexts. a. , Enhancer nomination in the held-out cell line K562 on the held-out chromosomes 9 and 10, distal element–gene pairs (*≥* 50 kb; 5,462 pairs, 18 CRISPRi-validated positives from the genome-wide screens of Gasperini et al. and Morris et al.). Recall@*K*: fraction of positives among the top *K* pairs ranked across all genes. *p*: one-sided paired permutation test of the AUPRC difference with Evo2HiC (Methods). Evo2HiC and Hi-C–only: ABC from the predicted DNase and H3K27ac tracks, quantile-normalized to the measured tracks of the training cell line GM12878 (no K562 data used). **b**, Loop-anchor classification under the ENCODE cCRE classes. Agreement between anchor labels derived from predicted and from measured tracks (promoter-like, enhancer-like, CTCF-only, low-DNase). 95% confidence intervals from 1,000 bootstrap resamples; chance is the accuracy under label shuffling. **c**, Validation of loop-anchor labels without measured epigenomic data. Motif AUROC: separation of anchors with and without a CTCF motif (JASPAR MA0139.2, *p <* 10*^−^*^5^) by the maximum predicted (or measured) CTCF signal. Convergent: fraction of loops with convergent CTCF motif orientation among loops whose two anchors are both called CTCF-positive (25% expected by chance). Expression AUROC: separation of genes at anchors called promoter-like from genes at anchors called CTCF-only or low-DNase, using ENCODE RNA-seq TPM.

| Score | AUPRC | $p$ | Recall@25 | Recall@50 | Recall@100 |
| --- | --- | --- | --- | --- | --- |
| Genomic distance | 0.039 | 0.017 | 0.00 | 0.11 | 0.28 |
| Hi-C contact | 0.043 | 0.006 | 0.00 | 0.06 | 0.33 |
| ABC, Hi-C-only | 0.123 | 0.12 | 0.22 | 0.22 | 0.33 |
| ABC, Evo2HiC | 0.221 | – | 0.33 | 0.33 | 0.39 |

**b**
| Cell line (anchors) | Method | Accuracy [95% CI] | Balanced acc. | $\kappa$ [95% CI] | Chance | $F_1$ promoter / enhancer / CTCF |
| --- | --- | --- | --- | --- | --- | --- |
| GM12878 (1,238) | Evo2HiC | 0.739 [0.716, 0.764] | 0.634 | 0.588 [0.552, 0.627] | 0.401 | 0.79 / 0.30 / 0.53 |
|  | Hi-C-only | 0.674 [0.645, 0.698] | 0.514 | 0.482 [0.445, 0.518] |  | 0.70 / 0.04 / 0.39 |
|  | Sequence-only | 0.578 [0.549, 0.603] | 0.468 | 0.334 [0.291, 0.371] |  | 0.55 / 0.15 / 0.41 |
| H1ESC (643) | Evo2HiC | 0.751 [0.715, 0.782] | 0.572 | 0.598 [0.543, 0.643] | 0.378 | 0.74 / 0.22 / 0.42 |
|  | Hi-C-only | 0.663 [0.627, 0.700] | 0.465 | 0.452 [0.403, 0.505] |  | 0.62 / 0.00 / 0.33 |
|  | Sequence-only | 0.726 [0.692, 0.759] | 0.562 | 0.550 [0.495, 0.599] |  | 0.75 / 0.31 / 0.38 |
| K562 (703) | Evo2HiC | 0.623 [0.589, 0.657] | 0.515 | 0.451 [0.404, 0.499] | 0.312 | 0.67 / 0.16 / 0.44 |
|  | Hi-C-only | 0.561 [0.525, 0.597] | 0.425 | 0.351 [0.304, 0.398] |  | 0.63 / 0.00 / 0.31 |
|  | Sequence-only | 0.585 [0.548, 0.620] | 0.454 | 0.390 [0.337, 0.441] |  | 0.61 / 0.04 / 0.45 |

**c**
| Cell line | Tracks used | CTCF motif AUROC | Convergent (both anchors CTCF+) | Expression AUROC |
| --- | --- | --- | --- | --- |
| GM12878 | Evo2HiC predicted | 0.797 | 0.714 | 0.852 |
|  | Hi-C-only predicted | 0.763 | 0.703 | 0.812 |
|  | Sequence-only predicted | 0.753 | 0.688 | 0.793 |
|  | Measured | 0.803 | 0.713 | 0.814 |
| H1ESC | Evo2HiC predicted | 0.848 | 0.763 | 0.731 |
|  | Hi-C-only predicted | 0.787 | 0.783 | 0.691 |
|  | Sequence-only predicted | 0.821 | 0.732 | 0.722 |
|  | Measured | 0.844 | 0.771 | 0.719 |
| K562 | Evo2HiC predicted | 0.848 | 0.661 | 0.715 |
|  | Hi-C-only predicted | 0.768 | 0.679 | 0.686 |
|  | Sequence-only predicted | 0.769 | 0.654 | 0.754 |
|  | Measured | 0.813 | 0.675 | 0.781 |

**Supplementary Figure 1:**
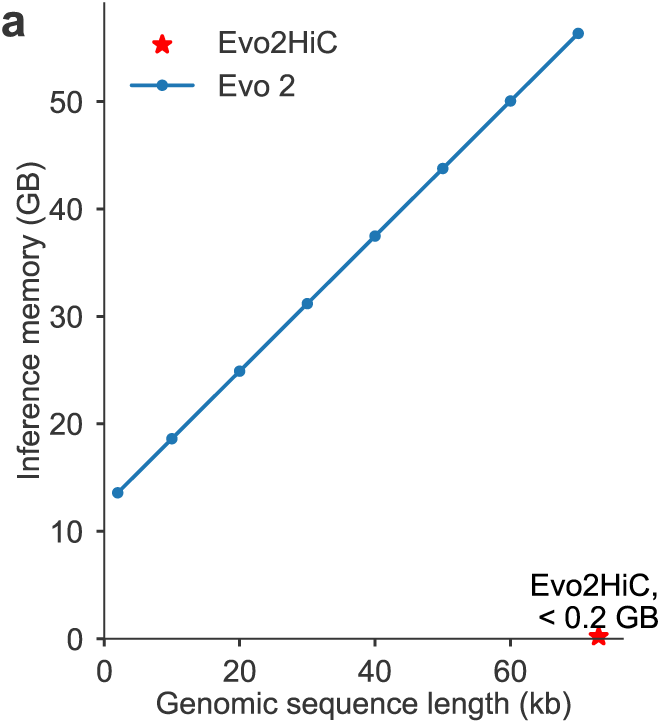
Comparison of the inference memory. Memory cost for the inference of one genomic sequence with different lengths between Evo 2 and Evo2HiC.

**Supplementary Figure 2:**
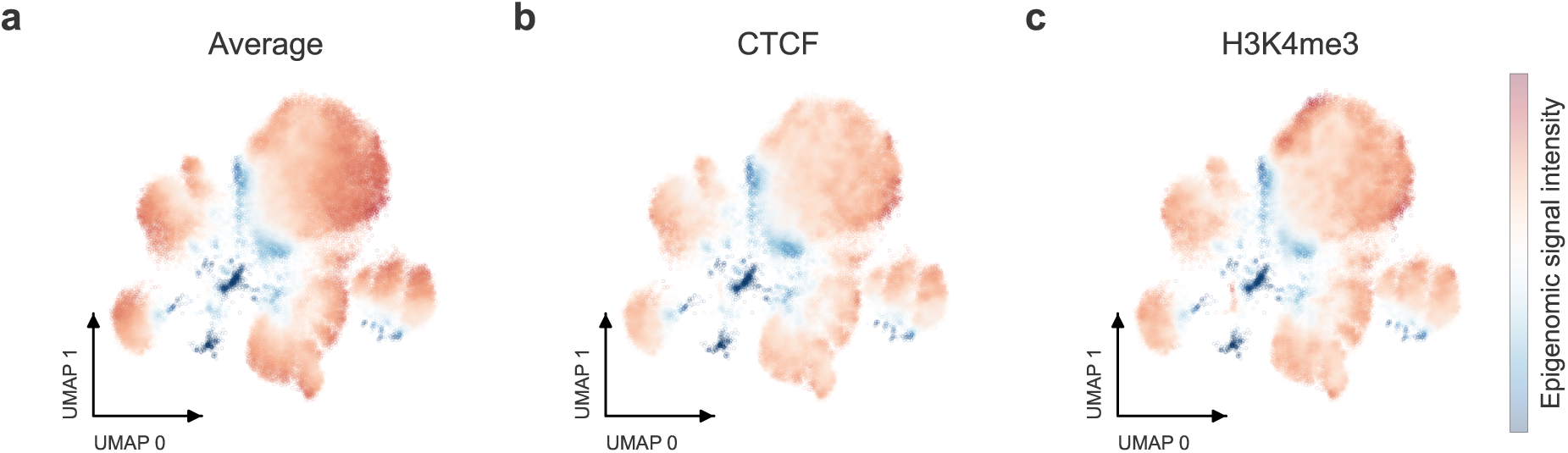
UMAP visualization. UMAP visualization of the embeddings of genomic sequences by Evo2HiC. **a** is the average epigenomic signal over CTCF, DNase, H3K27ac, H3K27me3, and H3K4me3. **b** is the epigenomic signal of CTCF and **c** is H3K4me3.

**Supplementary Figure 3:**
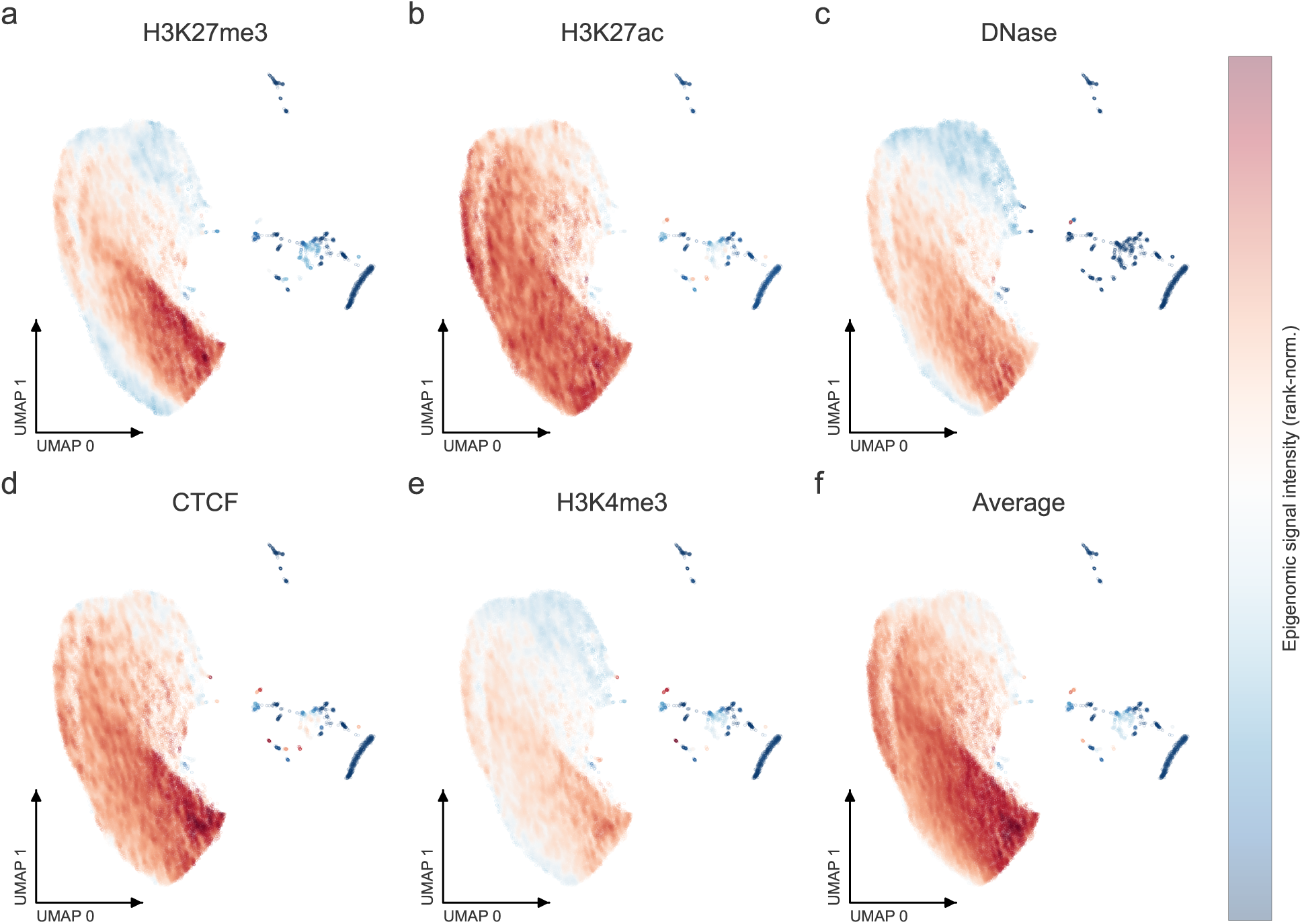
UMAP visulization of Evo 2. Visualization of the embeddings of genomic sequences by Evo 2, colored by epigenomic signals of H3K27me3 (a), H3K27ac (b), DNase (c), CTCF (d), H3K4me3 (e), and average signal over all assays (f).

**Supplementary Figure 4:**
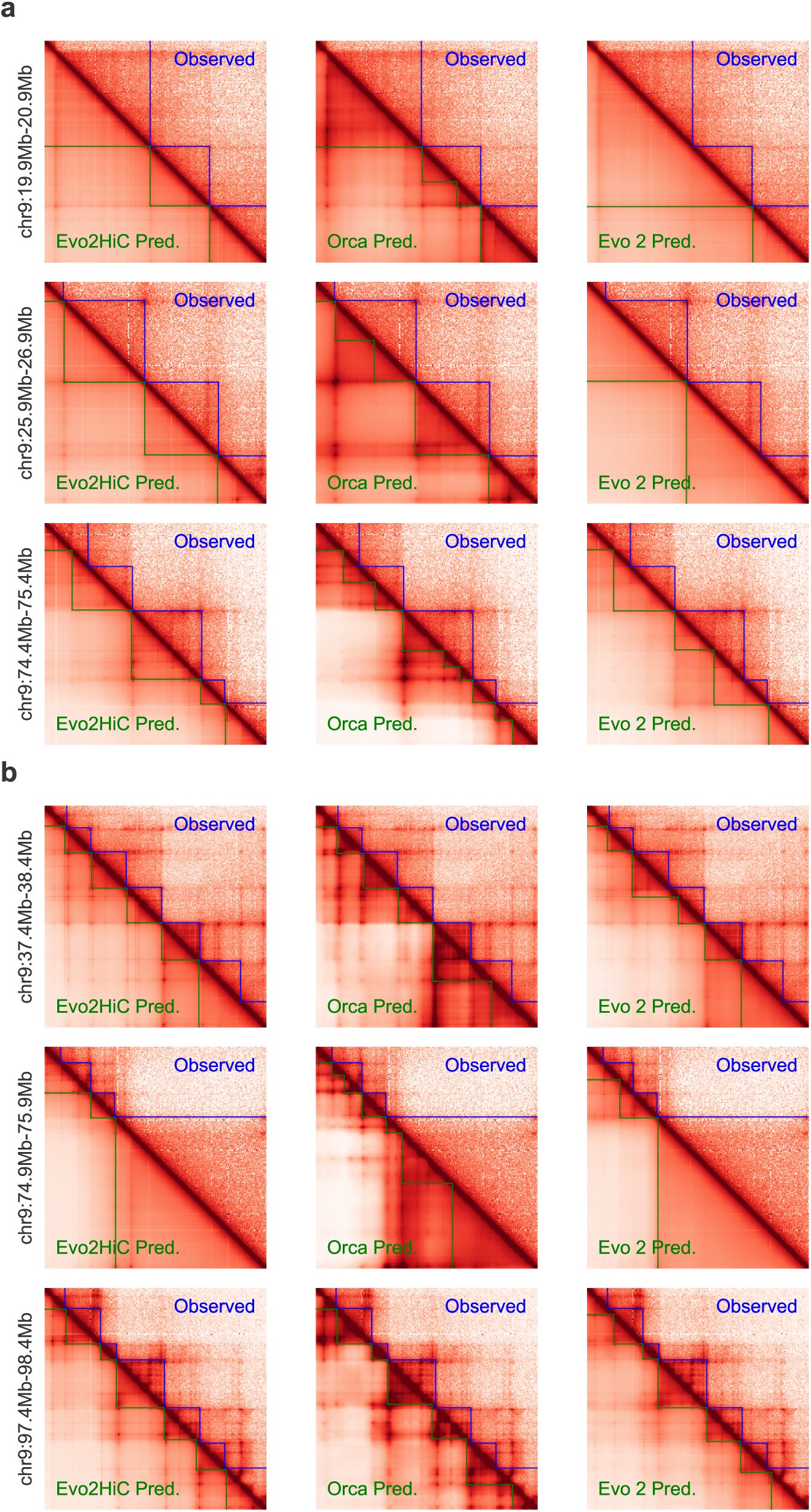
Visualization of genome structure prediction from sequence. **a,b**, Visualization of predicted and observed Hi-C contact maps for the H1ESC cell line (**a**) and the HFF cell line (**b**). In each visualization, the lower-left corner shows the predicted matrices from Evo2HiC, Orca, and Evo 2, respectively, while the top-right corner shows the observed matrix.

**Supplementary Figure 5:**
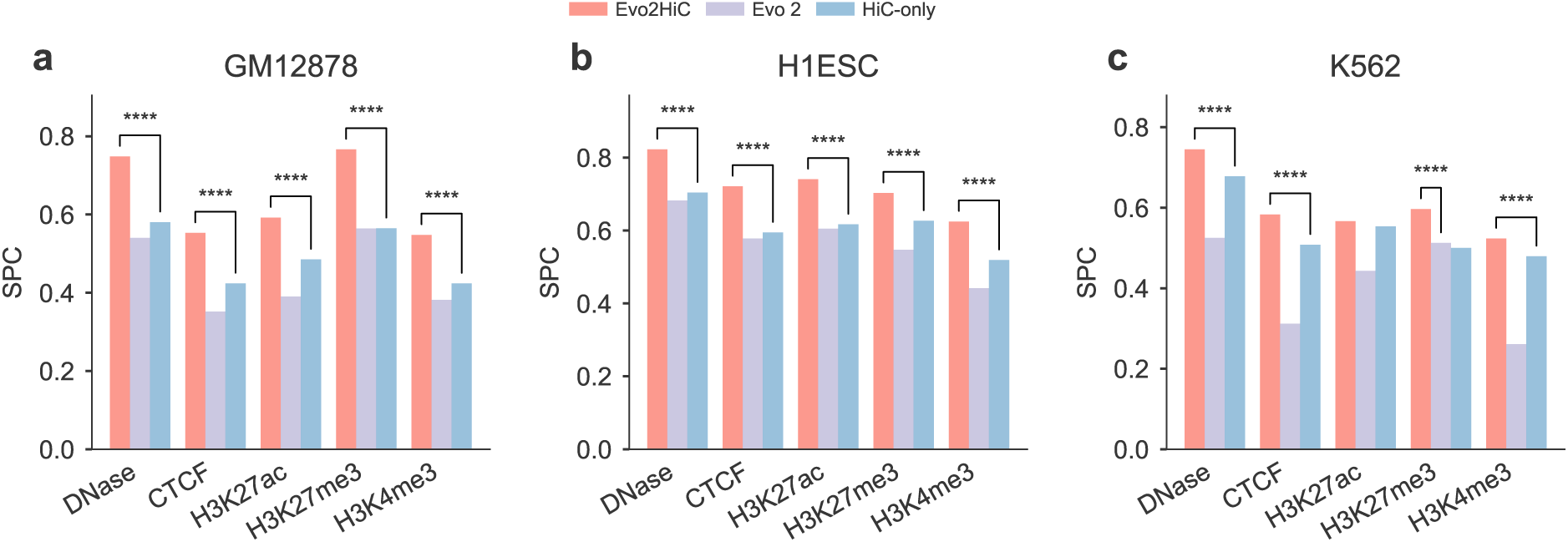
Evaluation on predicting epigenomic signals. **a,b**, Comparison on predicting epigenomic signals in the cross chromosome setting on GM12878 (**a**) and H1ESC (**b**). **c,** Comparison on predicting epigenomic signals in the cross cell line and cross chromosome setting on K562.

**Supplementary Figure 6:**
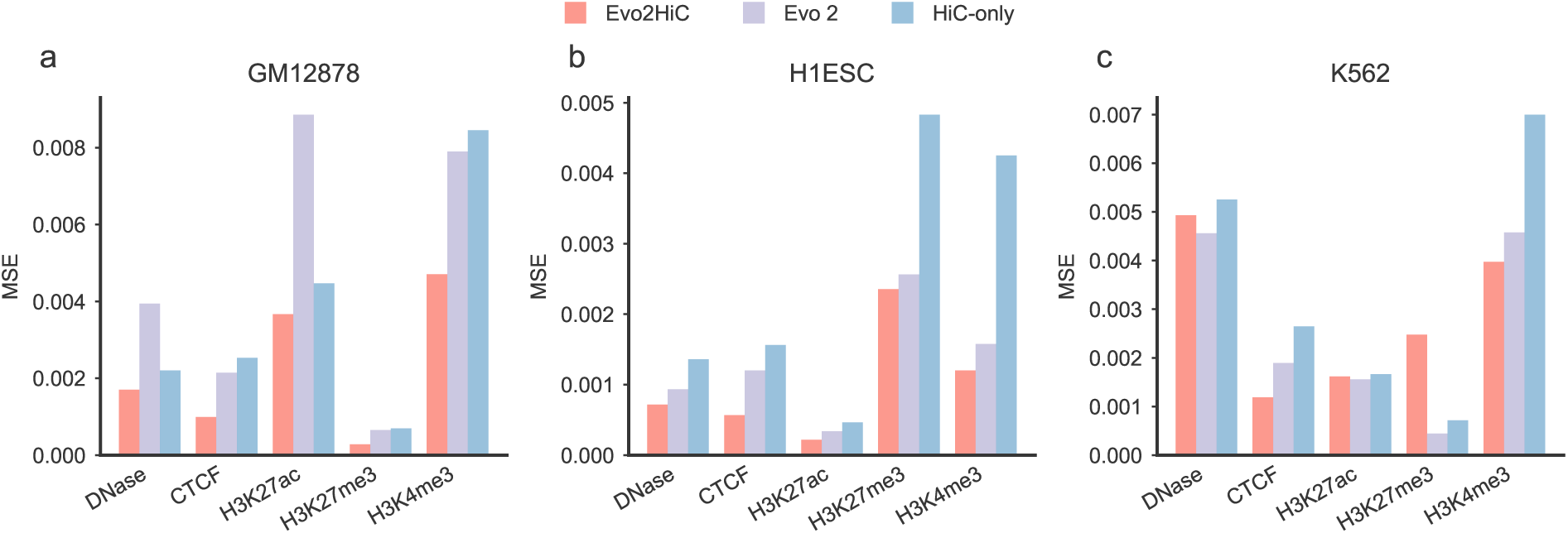
Comparison on predicting epigenomic signals. **a,b**, Comparison on predicting epigenomic signals in the cross chromosome setting on GM12878 (**a**) and H1ESC (**b**). **c,** Comparison on predicting epigenomic signals in the cross cell line and cross chromosome setting on K562. All differences are significant, as determined by Wilcoxon one-sided test.

**Supplementary Figure 7:**
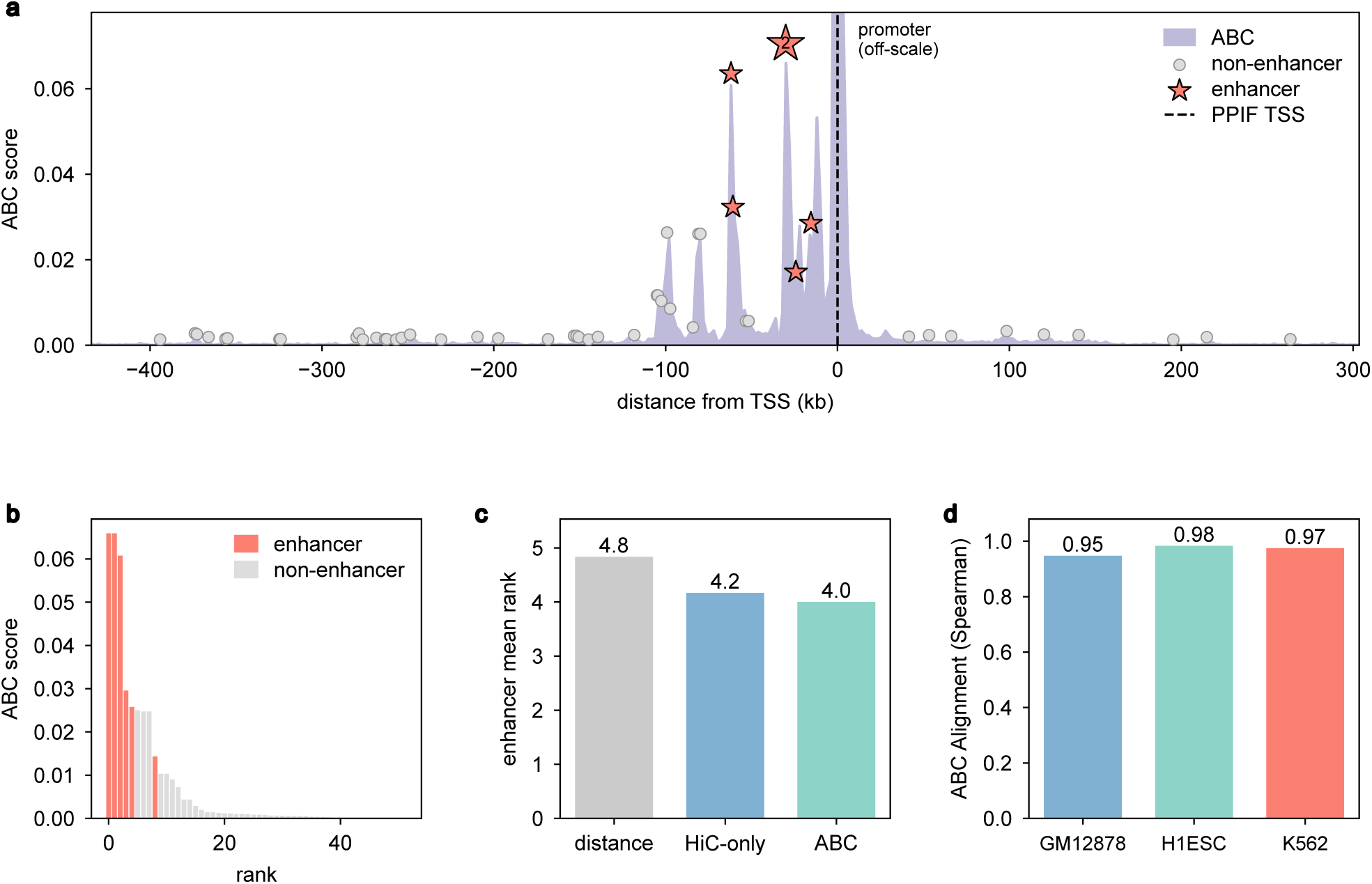
Activity-by-Contact (ABC) analysis for enhancer nomination. **a**, ABC score across the candidate region for *PPIF* in GM12878, spanning the 52 candidate elements. **b**, The 52 candidate elements ranked by ABC score, with validated enhancers highlighted in orange-red. **c**, Mean rank of validated enhancers under three ranking strategies—ABC, Hi-C contact only, and genomic distance; the ABC score yields the lowest (best) mean rank. **d**, Spearman correlation between ABC scores derived from Evo2HiC’s predicted signals and those computed from experimentally measured signals across three cell lines, all exceeding 0.95, indicating strong agreement. CRISPRi-FlowFISH data for *PPIF* in GM12878 are from Nasser et al. [63], as uniformly processed by ENCODE-rE2G [57].

**Supplementary Figure 8:**
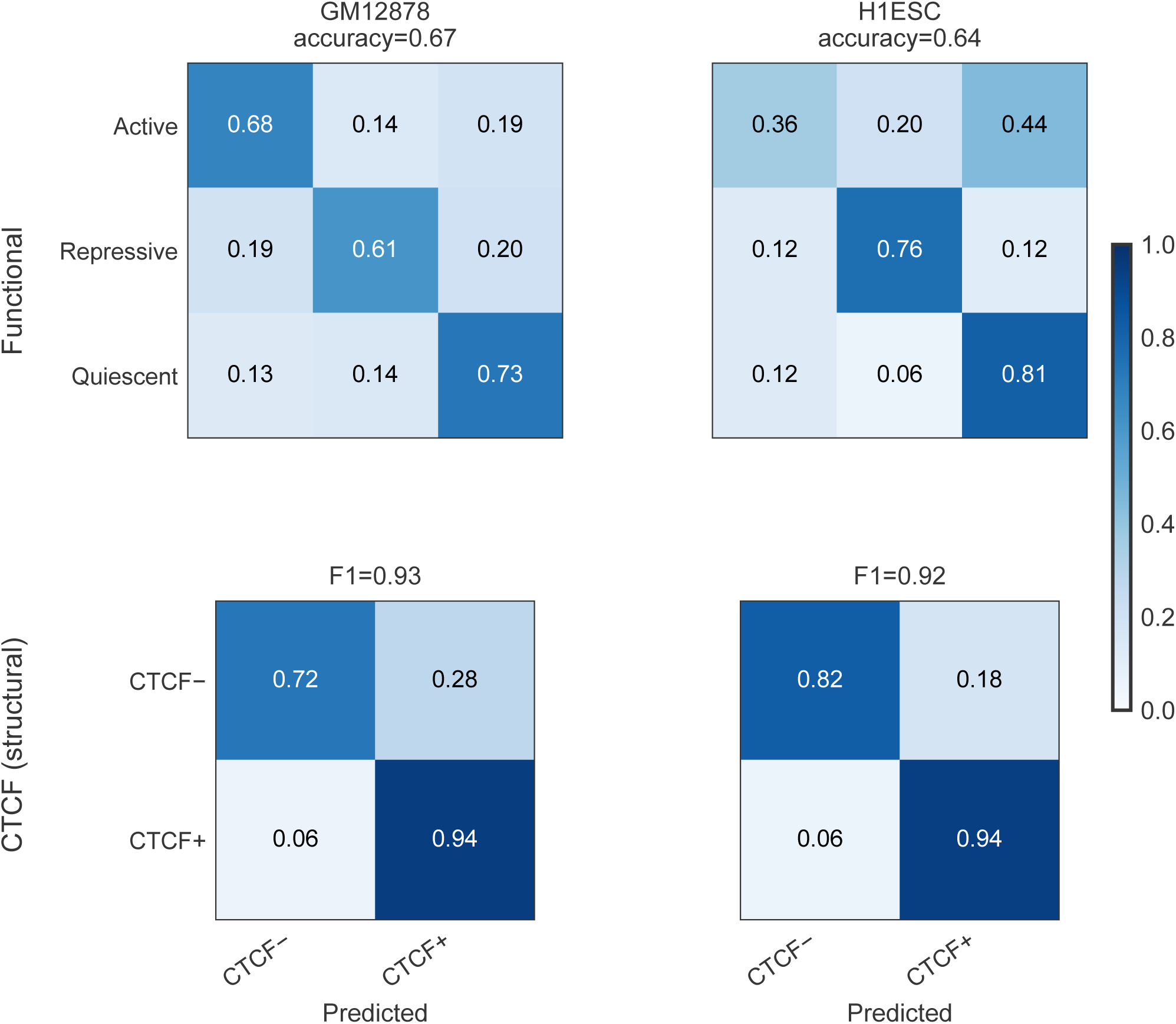
Consistency of loop anchor classification between predicted and measured epigenomic signals. Confusion matrices comparing loop anchor labels assigned from Evo2HiC’s predicted epigenomic profiles against those assigned from experimentally measured signals. Top row, classification by regulatory function (active, repressive, quiescent). Bottom row, classification by CTCF status (CTCF-positive vs. CTCF-negative). High agreement along the diagonals indicates that anchor types inferred from predicted profiles closely match those derived from measured data.

**Supplementary Figure 9:**
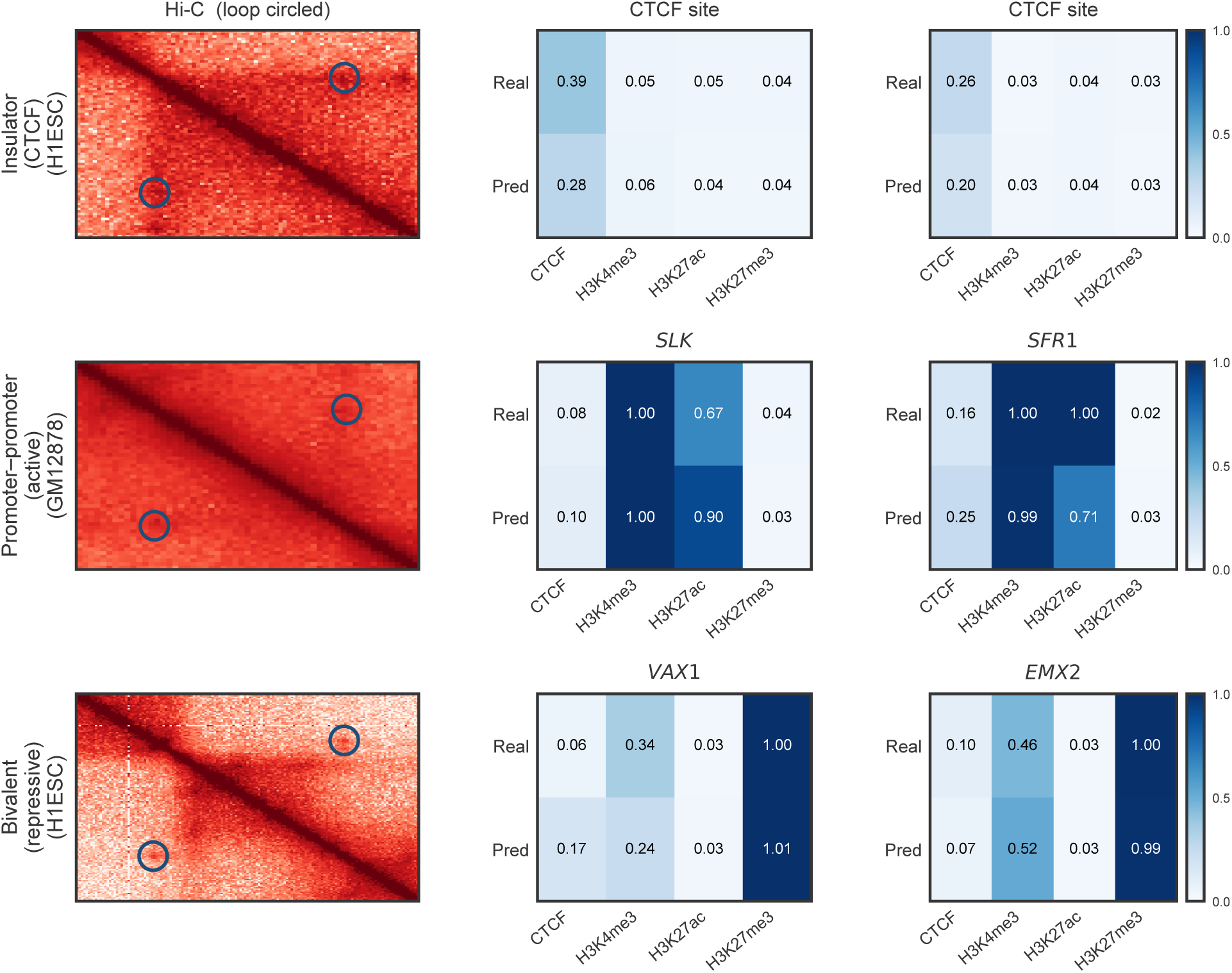
Representative chromatin loops with distinct epigenomic signatures at their anchors. Each row shows a representative loop, with Evo2HiC’s predicted epigenomic profiles displayed at both anchors. The clearly separable signatures across the three loop types are consistent with their expected regulatory biology.

**Supplementary Figure 10:**
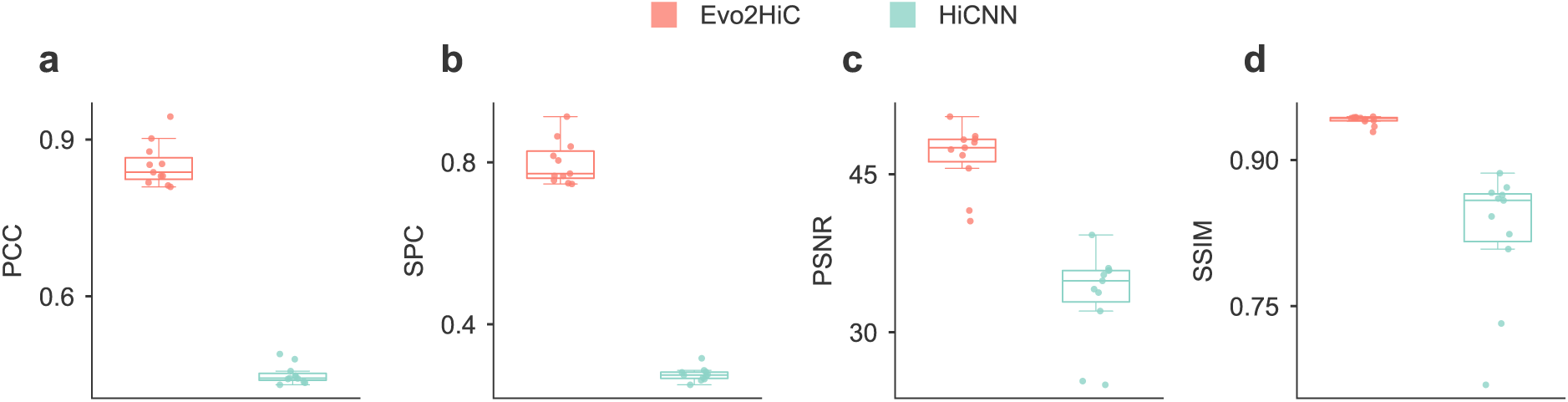
Comparison on resolution enhancement in mouse across four metrics between Evo2HiC and HiCNN.

**Supplementary Figure 11:**
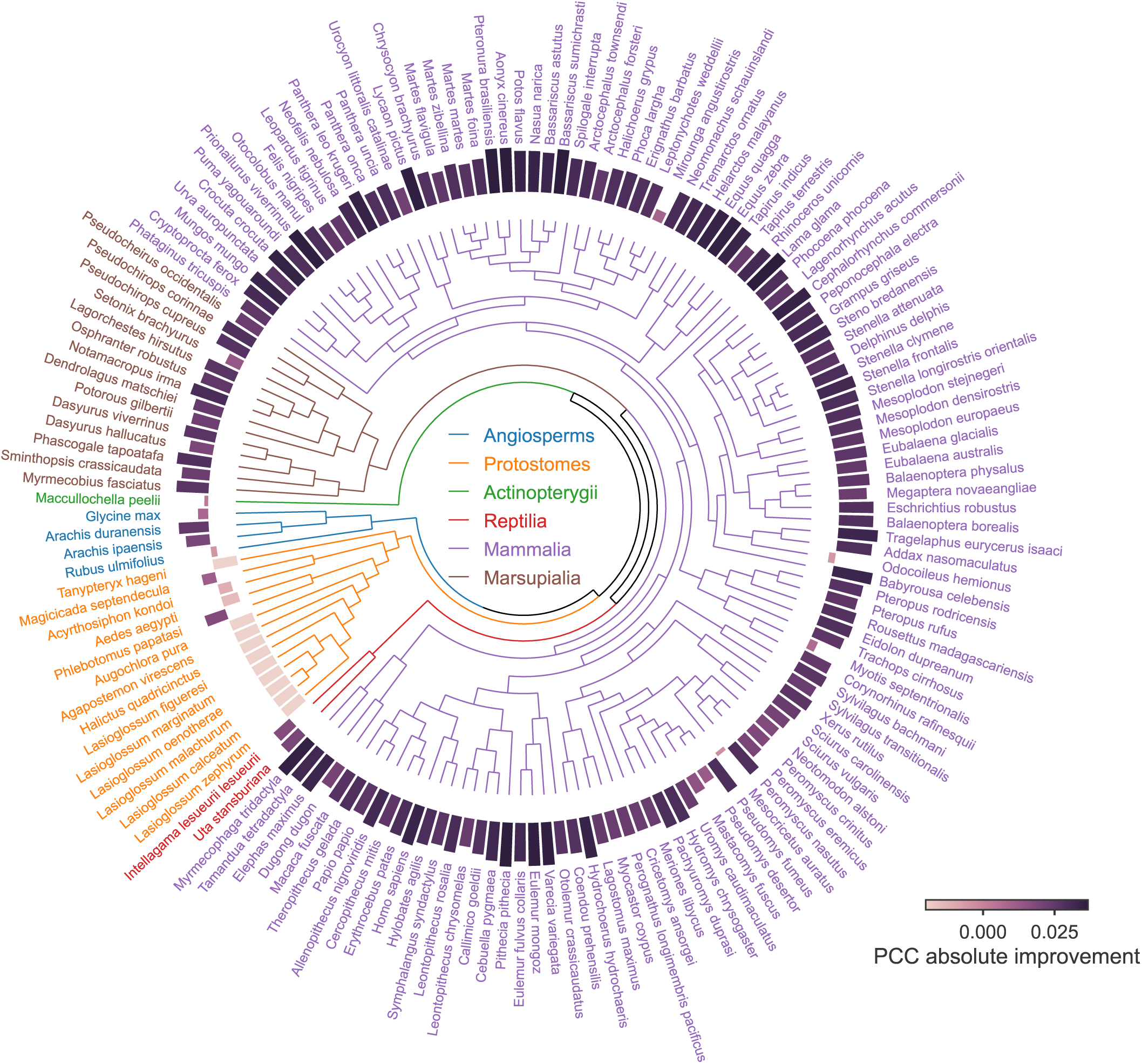
Comparison on resolution enhancement on 177 species using DNA Zoo. Bar heights indicate the relative improvement over HiCARN2 by Evo2HiC in terms of Pearson correlation.

**Supplementary Figure 12:**
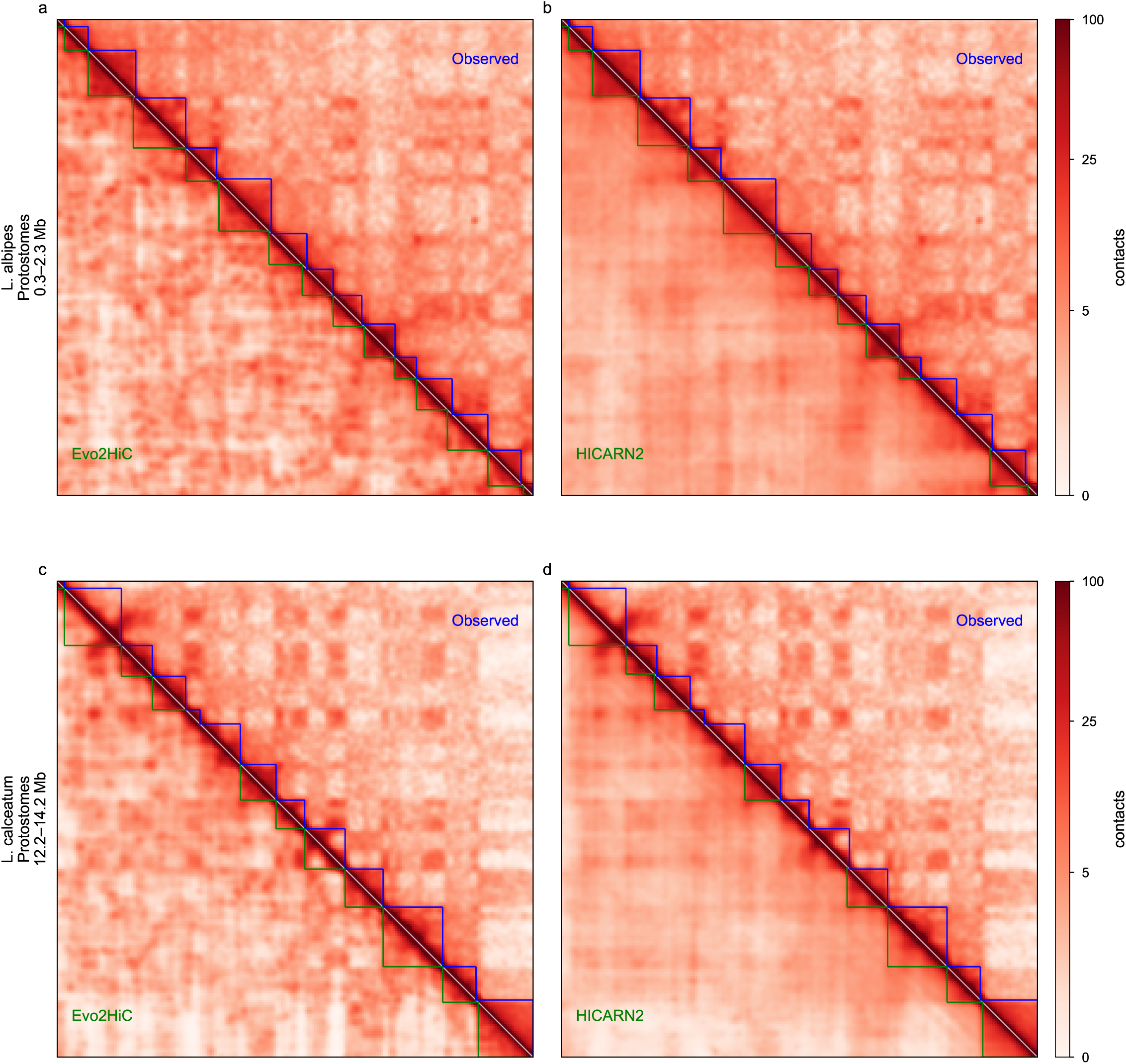
Evo2HiC resolves fine-grained chromatin structures missed by baseline models. **a–d**, Observed (upper-right) versus predicted (lower-left) Hi-C heatmaps and TAD boundaries (blue/green brackets) for 2-Mb regions in *Protostomes*. Evo2HiC (**a, c**) recovers fine-grained features with higher structural fidelity, which further benefits downstream analyses.

**Supplementary Figure 13:**
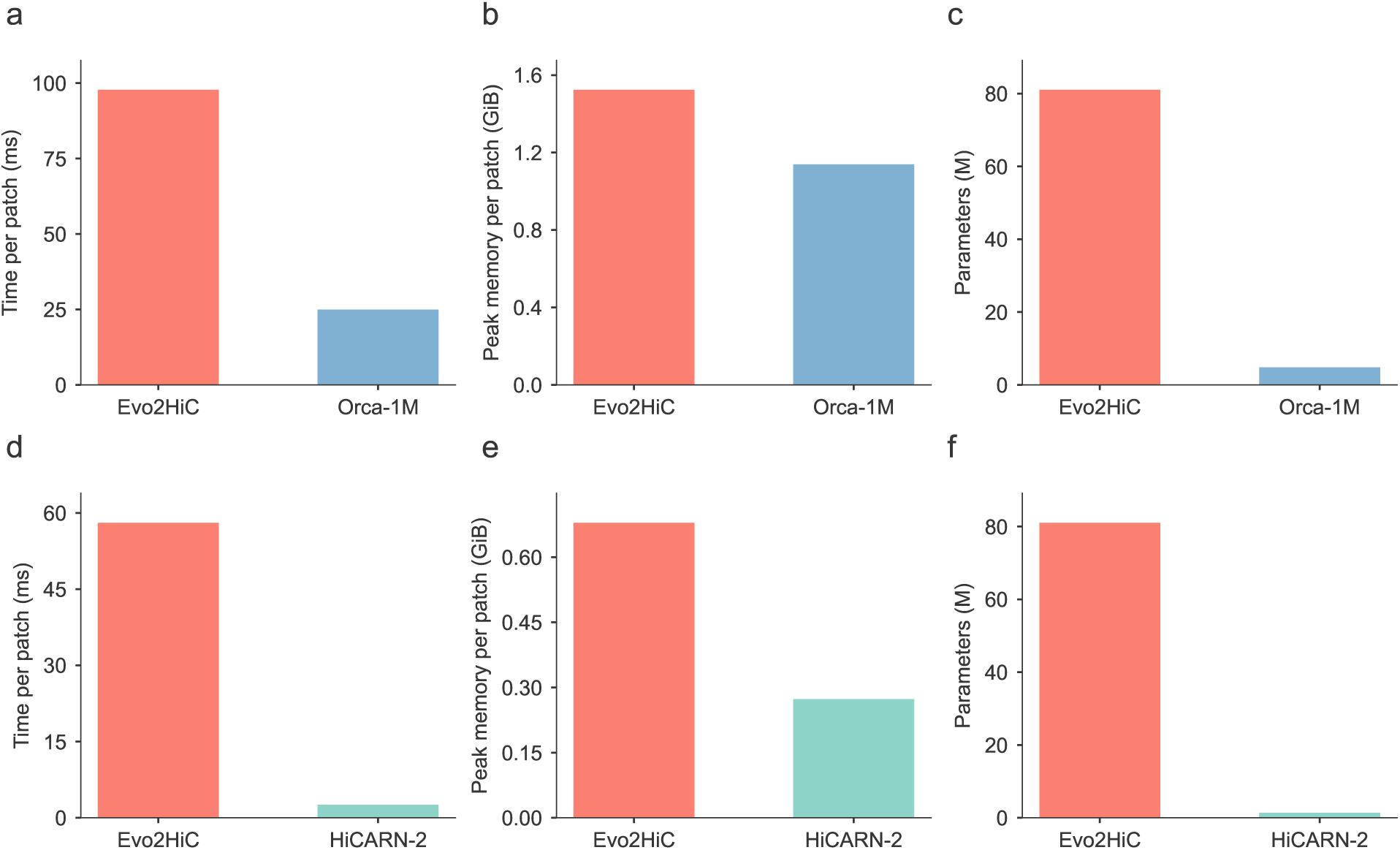
Comparison on inference cost and memory. **a-c**, inference cost comparison between Evo2HiC and Orca on Hi-C contact matrix prediction of one 1 Mb *×* 1 Mb patch prediction. **d-f**, inference cost comparison between Evo2HiC and HiCARN2 on Hi-C resolution enhancement of one 320 kb *×* 320 kb patch prediction.

**Supplementary Figure 14:**
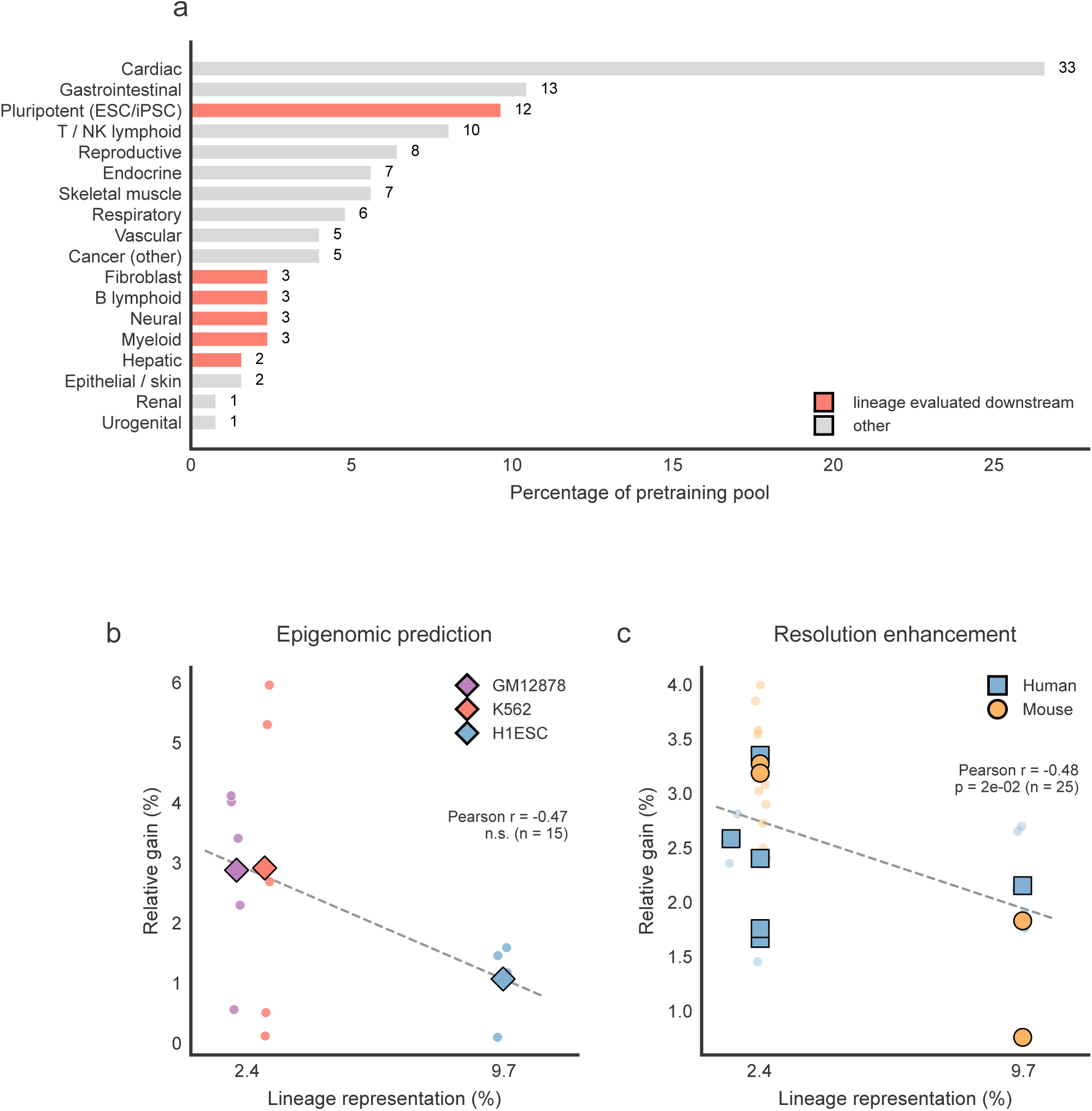
Downstream performance gain is not driven by representation in the pretraining data. **a**, Distribution of the pretraining data pool grouped by lineage, with lineages relevant to the downstream evaluation highlighted. **b**, Performance gain of Evo2HiC in epigenomic prediction versus the representation of the corresponding cell type in the pretraining data; the association is not significant (n.s.). **c**, Performance gain in Hi-C resolution enhancement versus representation in the pretraining data, where a negative correlation is observed. Together, these results indicate that downstream performance gains are primarily governed by factors such as task difficulty and other effects, rather than by the representation in the pretraining data distribution.

**Supplementary Figure 15:**
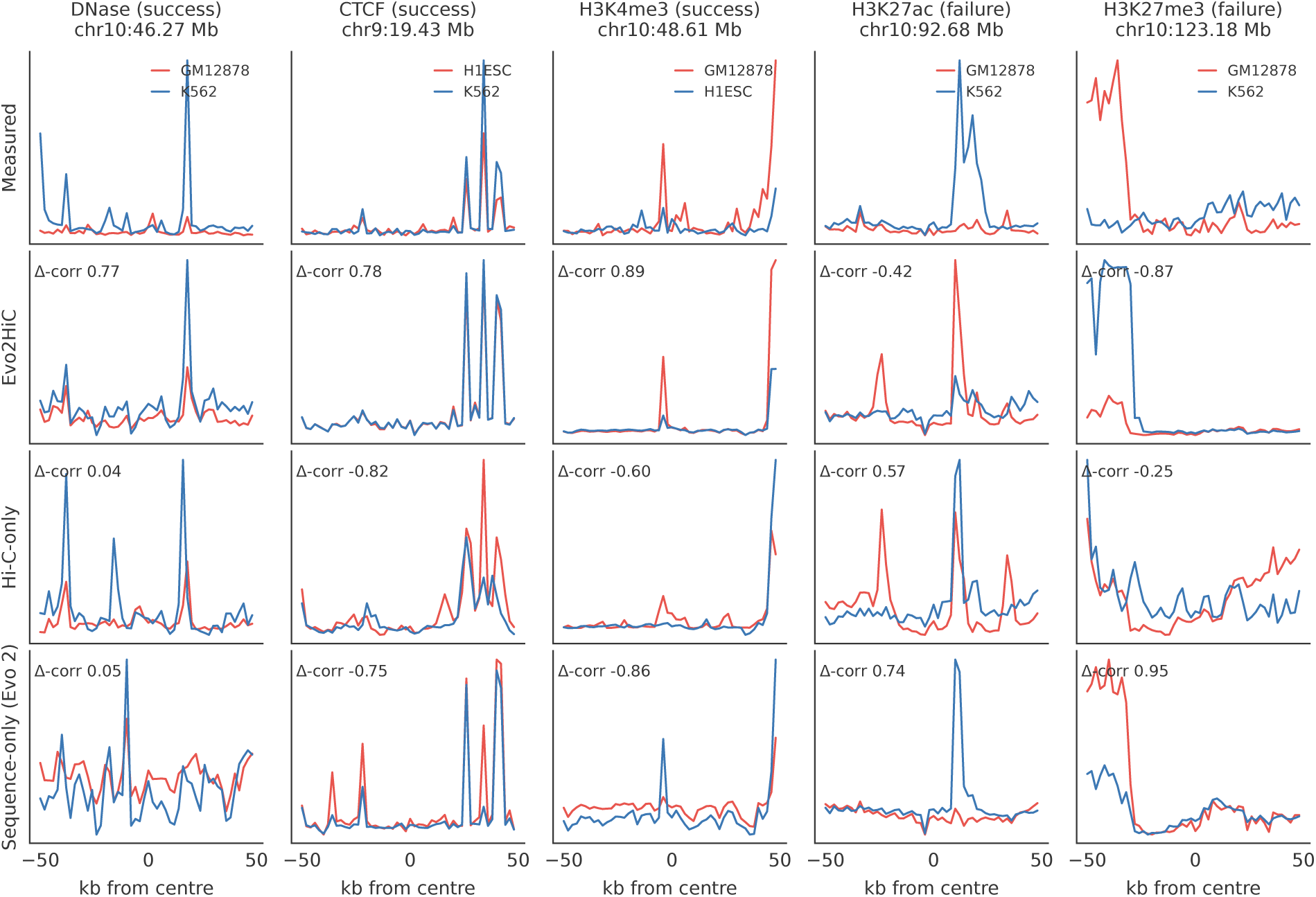
Representative loci for the cell type-differential analysis, including unsuccessful assays. For each assay, one 100 kb window is shown for a pair of cell lines, with measured signals (top) and the predictions of Evo2HiC, the Hi-C–only baseline and the sequence-only (Evo 2) baseline (rows 2–4) for both cell lines. Δ-corr is the correlation between the predicted and measured between-cell differences within the window. Windows were selected by a fixed rule among windows in the top 10% of measured cell-type difference amplitude: for successes (DNase, CTCF, H3K4me3), the window where the Δ-corr of Evo2HiC most exceeds that of the better baseline; for failures (H3K27ac, H3K27me3), the window where the Δ-corr of the sequence-only baseline most exceeds that of Evo2HiC. Per-assay effect sizes are given in **Supplementary Table 13**.

**Supplementary Figure 16:**
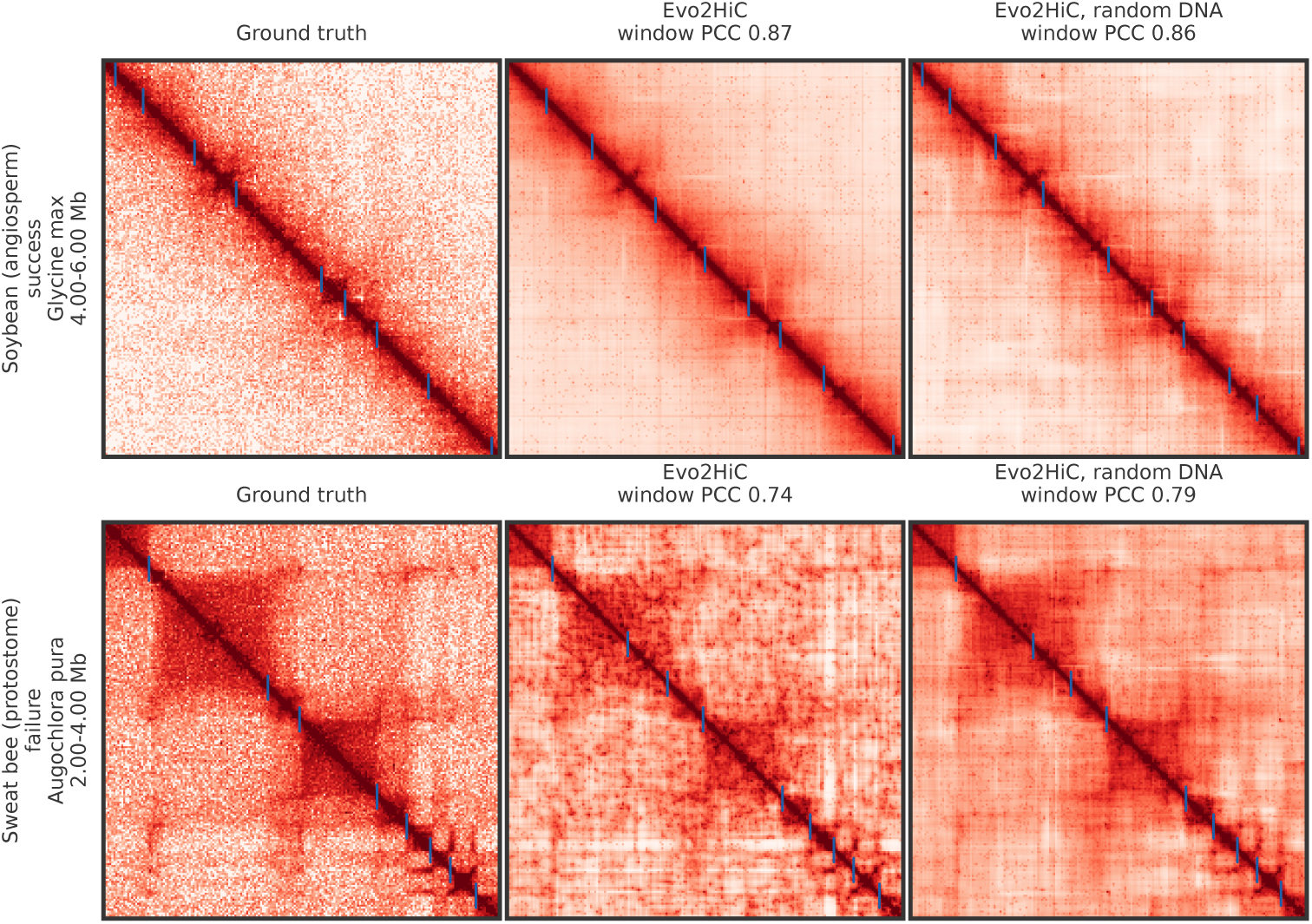
Distant-clade examples of the sequence prior. Ground-truth contact map, Evo2HiC prediction and the prediction of the same Evo2HiC model run on random DNA, with TAD boundaries (paper protocol) marked in blue. Species were selected by a fixed rule: among protostomes and angiosperms whose Evo2HiC pixel PCC is at or above the clade median, the species with the largest gain (soybean, *Glycine max*, +0.022) and the largest loss (sweat bee, *Augochlora pura*, *−*0.102) relative to random DNA. The 2 Mb window shown is the one whose local gain is closest to the species’ median window gain. In the sweat bee, the sequence input degrades the domain structure that the Hi-C input alone recovers.

**Supplementary Table 1:** Accession codes of the datasets used for predicting epigenomic profiles.

| **Assay** | **GM12878** | **H1ESC** | **K562** |
| --- | --- | --- | --- |
| Hi-C | 4DNFI1UEG1HD | 4DNFIQYQWPF5 | 4DNFITUOMFUQ |
| DNase | ENCFF960FMM | ENCFF573NKX | ENCFF414OGC |
| CTCF | ENCFF644EEX | ENCFF332TNJ | ENCFF534ITO |
| H3K27ac | ENCFF469WVA | ENCFF919FBG | ENCFF381NDD |
| H3K27me3 | ENCFF486WAD | ENCFF321JUJ | ENCFF242ENK |
| H3K4me3 | ENCFF280PUF | ENCFF493QWY | ENCFF911JVK |

**Supplementary Table 2:** Accession codes of the RNA-seq datasets used for validating cell type-specific motifs.

| **Assay** | **GM12878** | **H1ESC** | **K562** |
| --- | --- | --- | --- |
| RNA-seq | ENCFF345SHY | ENCFF910OBU | ENCFF928NYA |

**Supplementary Table 3:** Statistical tests for Fig 3 (Wilcoxon signed-rank, one-sided *greater*, Evo2HiC *>* comparator.

| Panel | Cell / scope | Comparison (left vs. right) | $n$ | $p$ |
| --- | --- | --- | --- | --- |
| Fig 3a | H1ESC, 0.0–0.2 Mb diagonals | Evo2HiC vs. Orca | 45 | $2.84 \times 10^{-14}$ |
| Fig 3a | H1ESC, 0.2–0.4 Mb diagonals | Evo2HiC vs. Orca | 50 | $8.88 \times 10^{-16}$ |
| Fig 3a | H1ESC, 0.4–0.6 Mb diagonals | Evo2HiC vs. Orca | 50 | $8.88 \times 10^{-16}$ |
| Fig 3a | H1ESC, 0.6–0.8 Mb diagonals | Evo2HiC vs. Orca | 50 | $8.88 \times 10^{-16}$ |
| Fig 3a | H1ESC, 0.8–1.0 Mb diagonals | Evo2HiC vs. Evo 2 | 50 | $8.88 \times 10^{-16}$ |
| Fig 3b | H1ESC per-window IS-PCC | Evo2HiC vs. Orca | 376 | $4.49 \times 10^{-15}$ |
| Fig 3e | HFF, 0.0–0.2 Mb diagonals | Evo2HiC vs. Orca | 45 | $9.13 \times 10^{-11}$ |
| Fig 3e | HFF, 0.2–0.4 Mb diagonals | Evo2HiC vs. Orca | 50 | $8.88 \times 10^{-16}$ |
| Fig 3e | HFF, 0.4–0.6 Mb diagonals | Evo2HiC vs. Orca | 50 | $8.88 \times 10^{-16}$ |
| Fig 3e | HFF, 0.6–0.8 Mb diagonals | Evo2HiC vs. Orca | 50 | $8.88 \times 10^{-16}$ |
| Fig 3e | HFF, 0.8–1.0 Mb diagonals | Evo2HiC vs. Orca | 50 | $8.88 \times 10^{-16}$ |
| Fig 3f | HFF per-window IS-PCC | Evo2HiC vs. Orca | 378 | $1.69 \times 10^{-14}$ |

**Supplementary Table 4:** Statistical tests for Fig 4 (Williams’ overlapping-correlation test, applied per chromosome and then Fisher-combined across chr 9 and chr 10; one-sided *greater*, Evo2HiC PCC *>* HiC-only PCC; 2 kb bins, *n* = 136 097 bins per cell-track combined across the two chromosomes). The test is skipped ΔPCC *≤* 0.

| Panel | Cell / track | PCC (Evo2HiC vs. HiC-only) | $p$ (Fisher-combined) |
| --- | --- | --- | --- |
| Fig 4a | GM12878 / DNase | 0.814 vs. 0.756 | $\leq 10^{-300}$ |
| Fig 4a | GM12878 / CTCF | 0.844 vs. 0.548 | $\leq 10^{-300}$ |
| Fig 4a | GM12878 / H3K27ac | 0.815 vs. 0.782 | $\leq 10^{-300}$ |
| Fig 4a | GM12878 / H3K27me3 | 0.735 vs. 0.483 | $\leq 10^{-300}$ |
| Fig 4a | GM12878 / H3K4me3 | 0.851 vs. 0.723 | $\leq 10^{-300}$ |
| Fig 4b | H1ESC / DNase | 0.831 vs. 0.657 | $\leq 10^{-300}$ |
| Fig 4b | H1ESC / CTCF | 0.853 vs. 0.526 | $\leq 10^{-300}$ |
| Fig 4b | H1ESC / H3K27ac | 0.734 vs. 0.622 | $\leq 10^{-300}$ |
| Fig 4b | H1ESC / H3K27me3 | 0.773 vs. 0.459 | $\leq 10^{-300}$ |
| Fig 4b | H1ESC / H3K4me3 | 0.915 vs. 0.667 | $\leq 10^{-300}$ |
| Fig 4c | K562 / DNase | 0.738 vs. 0.664 | $\leq 10^{-300}$ |
| Fig 4c | K562 / CTCF | 0.820 vs. 0.481 | $\leq 10^{-300}$ |
| Fig 4c | K562 / H3K27ac | 0.549 vs. 0.541 | $2.65 \times 10^{-168}$ |
| Fig 4c | K562 / H3K27me3 | 0.290 vs. 0.323 | – (skip) |
| Fig 4c | K562 / H3K4me3 | 0.811 vs. 0.625 | $\leq 10^{-300}$ |

**Supplementary Table 5:** Statistical tests for Fig 5f (per-motif two-proportion *z*-test on foreground vs. background rate, one-sided *greater*, Benjamini–Hochberg FDR within each cell line; *n* = 460 motifs per cell).

| Cell | Motif | $q$ |
| --- | --- | --- |
| GM12878 | CTCF_MA0139.1 | 0.0 |
| H1ESC | CTCF_MA0139.1 | 0.0 |
| K562 | CTCF_MA0139.1 | 0.0 |
| GM12878 | GLI2_MA0734.1 | $7.82 \times 10^{-1}$ |
| H1ESC | GLI2_MA0734.1 | $3.30 \times 10^{-4}$ |
| K562 | GLI2_MA0734.1 | $9.96 \times 10^{-1}$ |
| GM12878 | IRF2_MA0051.1 | 0.0 |
| H1ESC | IRF2_MA0051.1 | $7.37 \times 10^{-1}$ |
| K562 | IRF2_MA0051.1 | $2.77 \times 10^{-1}$ |
| GM12878 | PRDM1_MA0508.2 | $3.99 \times 10^{-15}$ |
| H1ESC | PRDM1_MA0508.2 | 1.00 |
| K562 | PRDM1_MA0508.2 | $4.58 \times 10^{-1}$ |
| GM12878 | RARG_NR_6 | $2.56 \times 10^{-1}$ |
| H1ESC | RARG_NR_6 | $9.63 \times 10^{-12}$ |
| K562 | RARG_NR_6 | $2.22 \times 10^{-1}$ |
| GM12878 | SPIB_MA0081.1 | $1.64 \times 10^{-8}$ |
| H1ESC | SPIB_MA0081.1 | 1.00 |
| K562 | SPIB_MA0081.1 | $3.54 \times 10^{-1}$ |
| GM12878 | YY1_MA0095.2 | 1.00 |
| H1ESC | YY1_MA0095.2 | $3.60 \times 10^{-4}$ |
| K562 | YY1_MA0095.2 | $6.32 \times 10^{-1}$ |

**Supplementary Table 6:** Statistical tests for Fig 6 (Wilcoxon signed-rank, one-sided *greater*, Evo2HiC vs. HiCARN2; paired across per-chromosome metric values). The saturated values 6.10 *×* 10*^−^*^5^ and 4.88 *×* 10*^−^*^4^ are the lower bounds for *n* = 14 and *n* = 11 respectively. Because every paired difference is positive, so the Wilcoxon *W* saturates at its maximum.

| Panel | Species | Metric | $n$ (chroms) | $p$ |
| --- | --- | --- | --- | --- |
| Fig 6a | Human | PCC | 14 | $6.10 \times 10^{-5}$ |
| Fig 6a | Human | SPC | 14 | $6.10 \times 10^{-5}$ |
| Fig 6a | Human | PSNR | 14 | $6.10 \times 10^{-5}$ |
| Fig 6a | Human | SSIM | 14 | $6.10 \times 10^{-5}$ |
| Fig 6b | Mouse | PCC | 11 | $4.88 \times 10^{-4}$ |
| Fig 6b | Mouse | SPC | 11 | $4.88 \times 10^{-4}$ |
| Fig 6b | Mouse | PSNR | 11 | $4.88 \times 10^{-4}$ |
| Fig 6b | Mouse | SSIM | 11 | $4.88 \times 10^{-4}$ |

**Supplementary Table 7:** Statistical tests for Extended Data Fig. 3 (Williams’ test on overlapping correlations using the Steiger SE, Fisher-combined across chr 9 and chr 10; one-sided *greater*, Evo2HiC vs. Borzoi; *n* = 28 792 chr-9 bins). The test is skipped when Δ*PCC <* 0.

| Panel / metric | Cell | Track | $p$ |
| --- | --- | --- | --- |
| ED Fig. 3 PCC | GM12878 | DNase | $\leq 10^{-300}$ |
| ED Fig. 3 PCC | GM12878 | CTCF | – (skip) |
| ED Fig. 3 PCC | GM12878 | H3K27ac | $\leq 10^{-300}$ |
| ED Fig. 3 PCC | GM12878 | H3K27me3 | – (skip) |
| ED Fig. 3 PCC | GM12878 | H3K4me3 | $\leq 10^{-300}$ |
| ED Fig. 3 PCC | H1ESC | DNase | – (skip) |
| ED Fig. 3 PCC | H1ESC | CTCF | – (skip) |
| ED Fig. 3 PCC | H1ESC | H3K27ac | – (skip) |
| ED Fig. 3 PCC | H1ESC | H3K27me3 | $2.54 \times 10^{-24}$ |
| ED Fig. 3 PCC | H1ESC | H3K4me3 | $4.62 \times 10^{-7}$ |
| ED Fig. 3 SPC | GM12878 | DNase | $1.90 \times 10^{-181}$ |
| ED Fig. 3 SPC | GM12878 | CTCF | $1.38 \times 10^{-50}$ |
| ED Fig. 3 SPC | GM12878 | H3K27ac | – (skip) |
| ED Fig. 3 SPC | GM12878 | H3K27me3 | $1.93 \times 10^{-7}$ |
| ED Fig. 3 SPC | GM12878 | H3K4me3 | $7.13 \times 10^{-128}$ |
| ED Fig. 3 SPC | H1ESC | DNase | $5.36 \times 10^{-5}$ |
| ED Fig. 3 SPC | H1ESC | CTCF | – (skip) |
| ED Fig. 3 SPC | H1ESC | H3K27ac | $6.43 \times 10^{-8}$ |
| ED Fig. 3 SPC | H1ESC | H3K27me3 | $4.51 \times 10^{-19}$ |
| ED Fig. 3 SPC | H1ESC | H3K4me3 | – (skip) |

**Supplementary Table 8:** Statistics for Extended Data Fig. 4, summarised per (ablation *×* cell). Each cell has 5 tracks; Williams’ 1-sided test *×* 2 chromosomes + Fisher’s method (one-sided *Evo2HiC PCC > baseline PCC*; *n* = 136 097 bins per tested cell-track). *p* range = min/max over tested tracks. Test is not performed when ΔPCC *≤* 0.

| Ablation | Cell | Tested | Gate-skipped | $p$ range |
| --- | --- | --- | --- | --- |
| No-distillation | GM12878 | 5/5 | 0 | $\leq 10^{-300}$ to $3.117 \times 10^{-23}$ |
| No-distillation | H1ESC | 5/5 | 0 | $1.052 \times 10^{-219}$ to $1.139 \times 10^{-4}$ |
| No-distillation | K562 | 5/5 | 0 | $\leq 10^{-300}$ to $1.745 \times 10^{-5}$ |
| Evo2 + HiC | GM12878 | 4/5 | 1 (H3K4me3) | $\leq 10^{-300}$ to $5.768 \times 10^{-5}$ |
| Evo2 + HiC | H1ESC | 2/5 | 3 (DNase, H3K27me3, H3K4me3) | $\leq 10^{-300}$ to $3.632 \times 10^{-80}$ |
| Evo2 + HiC | K562 | 3/5 | 2 (H3K27ac, H3K4me3) | $\leq 10^{-300}$ to $1.420 \times 10^{-3}$ |
| Ridge (HiC) | GM12878 | 5/5 | 0 | $\leq 10^{-300}$ to $\leq 10^{-300}$ |
| Ridge (HiC) | H1ESC | 5/5 | 0 | $\leq 10^{-300}$ to $\leq 10^{-300}$ |
| Ridge (HiC) | K562 | 4/5 | 1 (H3K27me3) | $\leq 10^{-300}$ to $\leq 10^{-300}$ |

**Supplementary Table 9:** Statistics for Extended Data Fig. 5c (Δ-signal Pearson), one row per track. The figure draws a significance bracket only where Evo2HiC has the highest mean SpecPCC among Evo2HiC / Evo 2 / HiC-only on that track; otherwise (gate not passed) no test is performed. For passed tracks, per-pair Williams’ Z (1-sided, pre-specified direction *r*_Evo2HiC_ *> r*_baseline_) across 3 unordered cell-pairs are Fisher-combined. *n* = 136 097 bins per cell-pair. Source: result/crosscell/williams_pvals.tsv.

| Track | Runner-up baseline | SpecPCC (E2H vs. RU) | Per-pair Williams Z | $p$ (Fisher) |
| --- | --- | --- | --- | --- |
| DNase | HiC-only | 0.541 vs. 0.435 | +78.6 / +79.5 / +21.6 | $\leq 10^{-300}$ |
| CTCF | HiC-only | 0.366 vs. 0.195 | +103.9 / +16.2 / +69.7 | $\leq 10^{-300}$ |
| H3K27ac | HiC-only | 0.592 vs. 0.596 | – (skip) | n.s. |
| H3K27me3 | Evo 2 | 0.329 vs. 0.569 | – (skip) | n.s. |
| H3K4me3 | Evo 2 | 0.551 vs. 0.381 | +120.2 / +82.3 / +17.0 | $\leq 10^{-300}$ |

**Supplementary Table 10:** Statistical tests for Extended Data Fig. 6 (Wilcoxon signed-rank, one-sided *greater*, Evo2HiC vs. Evo2HiC(HiC-only) paired tests).

| Panel | Species | Metric | $n$ (chroms) | $p$ |
| --- | --- | --- | --- | --- |
| ED Fig. 6a | Human | PCC | 14 | $6.10 \times 10^{-5}$ |
| ED Fig. 6a | Human | SPC | 14 | $6.10 \times 10^{-5}$ |
| ED Fig. 6a | Human | PSNR | 14 | $6.10 \times 10^{-5}$ |
| ED Fig. 6a | Human | SSIM | 14 | $6.10 \times 10^{-5}$ |
| ED Fig. 6b | Mouse | PCC | 11 | $4.88 \times 10^{-4}$ |
| ED Fig. 6b | Mouse | SPC | 11 | $4.88 \times 10^{-4}$ |
| ED Fig. 6b | Mouse | PSNR | 11 | $9.77 \times 10^{-4}$ |
| ED Fig. 6b | Mouse | SSIM | 11 | $4.88 \times 10^{-4}$ |

**Supplementary Table 11:** Hi-C contact matrices prediction performance in Orca’s evaluation setting. Performance at 1 Mb / 4 kb resolution on held-out test chromosomes (chr9, chr10) in log(O/E) space. Pearson (PCC) and Spearman (SPC) correlations are computed globally across all 417 windows following the Orca evaluation protocol.

| Method | H1-ESC |  | HFF |  |
| --- | --- | --- | --- | --- |
|  | PCC | SPC | PCC | SPC |
| Orca | 0.776 | 0.762 | 0.740 | 0.718 |
| Evo2HiC | 0.672 | 0.679 | 0.581 | 0.607 |

**Supplementary Table 12:**
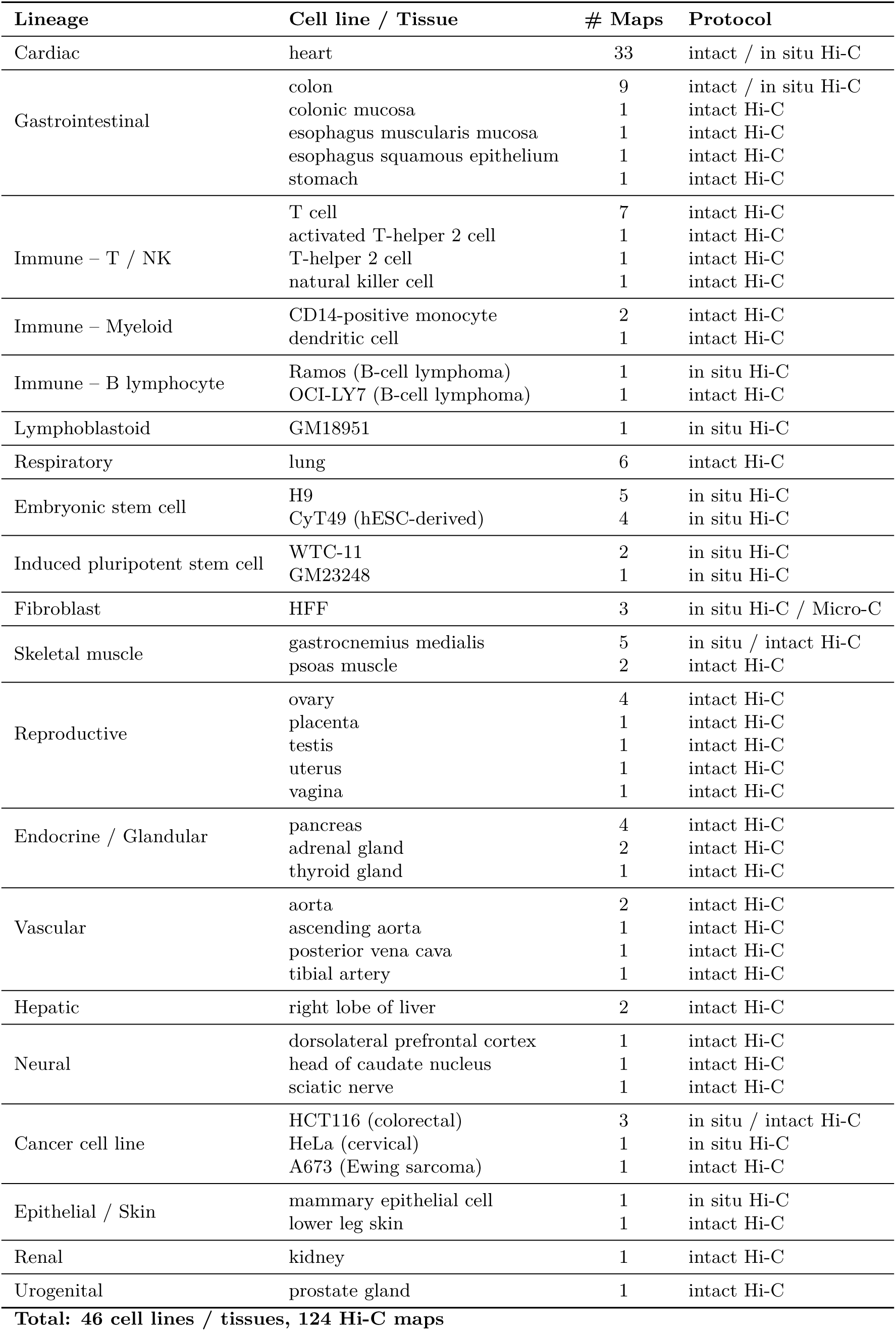
Cell lines and tissues used for pretraining. All 124 human Hi-C maps from 46 cell lines/tissues in the training split of are listed, grouped by lineage.

**Supplementary Table 13:**
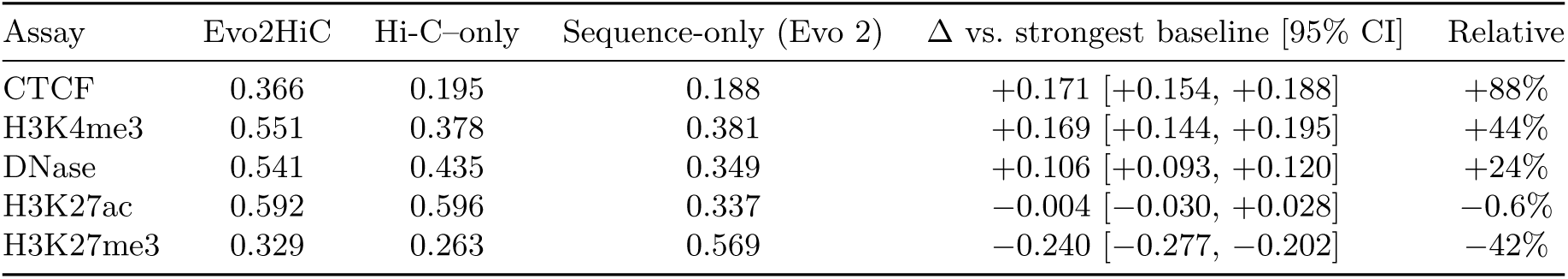
Per-assay effect sizes of the cell type-differential analysis. Correlation between predicted and measured between-cell signal differences over all 2 kb bins of chromosomes 9 and 10, averaged over the three pairs of GM12878, H1ESC and K562 (K562 held out). Δ: Evo2HiC minus the strongest baseline for that assay; 95% confidence interval from a paired block bootstrap over genomic bins (100 kb blocks, 2,000 resamples).

| Assay | Evo2HiC | Hi-C-only | Sequence-only (Evo 2) | $\Delta$ vs. strongest baseline [95% CI] | Relative |
| --- | --- | --- | --- | --- | --- |
| CTCF | 0.366 | 0.195 | 0.188 | +0.171 [+0.154, +0.188] | +88% |
| H3K4me3 | 0.551 | 0.378 | 0.381 | +0.169 [+0.144, +0.195] | +44% |
| DNase | 0.541 | 0.435 | 0.349 | +0.106 [+0.093, +0.120] | +24% |
| H3K27ac | 0.592 | 0.596 | 0.337 | -0.004 [-0.030, +0.028] | -0.6% |
| H3K27me3 | 0.329 | 0.263 | 0.569 | -0.240 [-0.277, -0.202] | -42% |

**Supplementary Table 14:** Loop-anchor classification under the rule-based classes (CTCF, active, repressive, quiescent). Agreement between anchor labels derived from predicted and from measured tracks; 95% confidence intervals from 1,000 bootstrap resamples; chance is the accuracy under label shuffling. The rule assigns H3K27me3-positive anchors to the repressive class first, so it depends on H3K27me3, the assay with the weakest held-out-cell prediction.

| Cell line | Method | Accuracy [95% CI] | Balanced accuracy [95% CI] | Chance |
| --- | --- | --- | --- | --- |
| GM12878 | Evo2HiC | 0.652 [0.627, 0.679] | 0.656 [0.628, 0.683] | 0.292 |
|  | Hi-C-only | 0.562 [0.536, 0.590] | 0.553 [0.524, 0.583] |  |
|  | Sequence-only | 0.476 [0.448, 0.504] | 0.473 [0.442, 0.502] |  |
| H1ESC | Evo2HiC | 0.658 [0.621, 0.692] | 0.654 [0.616, 0.690] | 0.295 |
|  | Hi-C-only | 0.557 [0.519, 0.596] | 0.495 [0.459, 0.530] |  |
|  | Sequence-only | 0.571 [0.532, 0.608] | 0.548 [0.508, 0.586] |  |
| K562 | Evo2HiC | 0.496 [0.461, 0.532] | 0.547 [0.504, 0.585] | 0.344 |
|  | Hi-C-only | 0.381 [0.347, 0.417] | 0.423 [0.384, 0.462] |  |
|  | Sequence-only | 0.498 [0.461, 0.535] | 0.456 [0.413, 0.500] |  |

**Supplementary Table 15:** Cross-species analysis (DNA Zoo, 177 species). a, b. , Contribution of the genome sequence, measured against two references without it: **a**, the same Evo2HiC checkpoint run with the genome replaced by random DNA of matched base composition; **b**, the separately trained Hi-C–only ablation. Clades are listed in order of divergence from human. Values are means over species (Evo2HiC/ reference); wins are species in which Evo2HiC is higher; *p*: two-sided paired Wilcoxon signed-rank test over species. Boundaries called and precision refer to TAD boundaries at the paper’s calling threshold; precision is the fraction of called boundaries within three bins of a ground-truth boundary. **c**, Comparison with other methods, by clade and over all species (means; wins of Evo2HiC against each method). Pixel PCC: distance-stratified correlation at 10 kb within 2 Mb. TAD-boundary *F*1: cooltools boundaries with prominence *≥* 0.2, a prediction being correct within three bins of a ground-truth boundary. Species without ground-truth boundaries or a random-DNA run are omitted from the corresponding counts. Per-species values, sequencing depth and divergence times are provided in **Supplementary Data 2**.

**a**
| Clade | $n$ | Pixel PCC | | | Boundaries called | | Boundary precision | | | TAD-boundary $F_1$ |
| --- | --- | --- | --- | --- | --- | --- | --- | --- | --- | --- |
| | | Evo2HiC/ random DNA | wins | $p$ | Evo2HiC/ random DNA | | Evo2HiC/ random DNA | wins | $p$ | Evo2HiC/ random DNA |
| Primates | 17 | 0.353 / 0.327 | 17/17 | $2 \times 10^{-5}$ | 498 / 707 | | 0.655 / 0.557 | 17/17 | $2 \times 10^{-5}$ | 0.566 / 0.575 |
| Other placental mammals | 109 | 0.374 / 0.346 | 108/109 | $1 \times 10^{-19}$ | 481 / 670 | | 0.669 / 0.567 | 108/109 | $2 \times 10^{-19}$ | 0.595 / 0.596 |
| Marsupials | 14 | 0.343 / 0.334 | 13/14 | $2 \times 10^{-4}$ | 1888 / 2483 | | 0.709 / 0.638 | 14/14 | $1 \times 10^{-4}$ | 0.575 / 0.603 |
| Reptiles | 5 | 0.297 / 0.290 | 3/5 | 0.62 | 958 / 1432 |  | 0.611 / 0.535 | 5/5 | 0.06 | 0.519 / 0.562 |
| Ray-finned fish | 1 | 0.236 / 0.233 | 1/1 | – | 122 / 170 |  | 0.672 / 0.659 | 1/1 | – | 0.491 / 0.588 |
| Protostomes | 26 | 0.392 / 0.424 | 6/26 | $4 \times 10^{-4}$ | 1213 / 1485 | | 0.732 / 0.759 | 7/23 | 0.02 | 0.684 / 0.731 |
| Angiosperms | 5 | 0.489 / 0.483 | 4/5 | 0.31 | 316 / 337 |  | 0.700 / 0.706 | 2/5 | 0.62 | 0.614 / 0.644 |

**b**
| Clade | $n$ | Pixel PCC | | | Boundaries called | | Boundary precision | | | TAD-boundary $F_1$ |
| --- | --- | --- | --- | --- | --- | --- | --- | --- | --- | --- |
| | | Evo2HiC/ Hi-C-only | wins | $p$ | Evo2HiC/ Hi-C-only | | Evo2HiC/ Hi-C-only | wins | $p$ | Evo2HiC/ Hi-C-only |
| Primates | 17 | 0.353 / 0.347 | 17/17 | $3 \times 10^{-4}$ | 498 / 603 | | 0.655 / 0.612 | 14/17 | $3 \times 10^{-3}$ | 0.566 / 0.587 |
| Other placental mammals | 109 | 0.374 / 0.369 | 91/109 | $1 \times 10^{-12}$ | 481 / 555 | | 0.669 / 0.627 | 94/109 | $1 \times 10^{-14}$ | 0.595 / 0.603 |
| Marsupials | 14 | 0.343 / 0.342 | 10/14 | 0.05 | 1888 / 2323 | | 0.709 / 0.668 | 12/14 | $5 \times 10^{-3}$ | 0.575 / 0.612 |
| Reptiles | 5 | 0.297 / 0.302 | 0/5 | 0.06 | 958 / 1059 |  | 0.611 / 0.608 | 2/5 | 0.81 | 0.519 / 0.544 |
| Ray-finned fish | 1 | 0.236 / 0.236 | 1/1 | – | 122 / 128 |  | 0.672 / 0.664 | 1/1 | – | 0.491 / 0.493 |
| Protostomes | 26 | 0.392 / 0.444 | 0/26 | $3 \times 10^{-8}$ | 967 / 827 | | 0.739 / 0.791 | 2/22 | $3 \times 10^{-6}$ | 0.684 / 0.690 |
| Angiosperms | 5 | 0.489 / 0.484 | 3/5 | 0.62 | 316 / 351 |  | 0.700 / 0.707 | 1/5 | 0.62 | 0.614 / 0.662 |

**c**
| Clade | Mean |  |  | Evo2HiC wins against |  |
| --- | --- | --- | --- | --- | --- |
|  | Evo2HiC | HiCARN2 | HiCFoundation | HiCARN2 | HiCFoundation |
| *Pixel PCC* |  |  |  |  |  |
| Primates | 0.353 | 0.323 | 0.347 | 17/17 | 17/17 |
| Other placental mammals | 0.374 | 0.346 | 0.369 | 107/109 | 87/109 |
| Marsupials | 0.343 | 0.318 | 0.336 | 14/14 | 12/14 |
| Reptiles | 0.297 | 0.285 | 0.286 | 5/5 | 4/5 |
| Ray-finned fish | 0.236 | 0.233 | 0.179 | 1/1 | 1/1 |
| Protostomes | 0.392 | 0.431 | 0.453 | 9/26 | 5/26 |
| Angiosperms | 0.489 | 0.481 | 0.504 | 3/5 | 1/5 |
| All species | 0.372 | 0.356 | 0.377 | 156/177 | 127/177 |
| *TAD-boundary $F_1$* |  |  |  |  |  |
| Primates | 0.566 | 0.545 | 0.535 | 13/17 | 13/17 |
| Other placental mammals | 0.595 | 0.572 | 0.554 | 81/109 | 80/109 |
| Marsupials | 0.575 | 0.560 | 0.536 | 10/14 | 10/14 |
| Reptiles | 0.519 | 0.505 | 0.405 | 3/5 | 4/5 |
| Ray-finned fish | 0.491 | 0.448 | 0.395 | 1/1 | 1/1 |
| Protostomes | 0.684 | 0.673 | 0.626 | 13/26 | 20/26 |
| Angiosperms | 0.614 | 0.615 | 0.553 | 2/5 | 5/5 |
| All species | 0.601 | 0.582 | 0.556 | 123/177 | 133/177 |

**Supplementary Table 16:** Boundaries added and removed by the sequence input. TAD boundaries (paper protocol) called on the Evo2HiC prediction and on a no-DNA reference were split into those called only with DNA (added) and only without DNA (removed). CTCF motif: fraction of boundaries within one bin of a JASPAR MA0139.2 match (*p <* 10*^−^*^5^); accuracy: fraction within three bins of a ground-truth boundary. Wins: species in which the added set exceeds the removed set; *p*: paired Wilcoxon signed-rank test over species. For reference, in mammals 62% of ground-truth boundaries and 24% of all bins carry a CTCF motif (protostomes: 26% and 10%).

| Group | $n$ | CTCF motif (added / removed) | wins | $p$ | accuracy (added / removed) | wins | $p$ |
| --- | --- | --- | --- | --- | --- | --- | --- |
| *no-DNA reference: same model on random DNA* |  |  |  |  |  |  |  |
| Primates | 17 | 0.72 / 0.43 | 17/17 | $2 \times 10^{-5}$ | 0.57 / 0.45 | 17/17 | $2 \times 10^{-5}$ |
| Glires | 23 | 0.77 / 0.56 | 23/23 | $2 \times 10^{-7}$ | 0.58 / 0.47 | 21/23 | $1 \times 10^{-6}$ |
| Laurasiatheria | 75 | 0.75 / 0.47 | 75/75 | $5 \times 10^{-14}$ | 0.57 / 0.43 | 72/75 | $9 \times 10^{-14}$ |
| Afrotheria/Xenarthra | 4 | 0.73 / 0.45 | 4/4 | 0.12 | 0.53 / 0.46 | 3/4 | 0.25 |
| Mammalia (all) | 126 | 0.75 / 0.48 | 126/126 | $2 \times 10^{-22}$ | 0.57 / 0.44 | 119/126 | $6 \times 10^{-22}$ |
| Marsupialia | 14 | 0.62 / 0.38 | 14/14 | $1 \times 10^{-4}$ | 0.62 / 0.53 | 13/14 | $4 \times 10^{-4}$ |
| Reptilia | 5 | 0.60 / 0.44 | 5/5 | 0.062 | 0.55 / 0.45 | 5/5 | 0.062 |
| Protostomes | 23 | 0.26 / 0.25 | 15/23 | 0.47 | 0.54 / 0.65 | 9/23 | 0.072 |
| Angiosperms | 5 | 0.21 / 0.11 | 4/5 | 0.12 | 0.53 / 0.61 | 2/5 | 0.31 |
| *no-DNA reference: Hi-C-only ablation* |  |  |  |  |  |  |  |
| Primates | 17 | 0.73 / 0.43 | 17/17 | $2 \times 10^{-5}$ | 0.55 / 0.49 | 12/17 | 0.023 |
| Glires | 23 | 0.79 / 0.55 | 23/23 | $2 \times 10^{-7}$ | 0.57 / 0.54 | 16/23 | 0.086 |
| Laurasiatheria | 75 | 0.75 / 0.47 | 75/75 | $5 \times 10^{-14}$ | 0.55 / 0.49 | 62/75 | $1 \times 10^{-8}$ |
| Afrotheria/Xenarthra | 4 | 0.73 / 0.44 | 4/4 | 0.12 | 0.50 / 0.48 | 2/4 | 0.88 |
| Mammalia (all) | 126 | 0.75 / 0.48 | 126/126 | $2 \times 10^{-22}$ | 0.55 / 0.49 | 98/126 | $2 \times 10^{-11}$ |
| Marsupialia | 14 | 0.63 / 0.38 | 14/14 | $1 \times 10^{-4}$ | 0.60 / 0.56 | 9/14 | 0.091 |
| Reptilia | 5 | 0.60 / 0.46 | 5/5 | 0.062 | 0.54 / 0.53 | 2/5 | 0.81 |
| Protostomes | 22 | 0.26 / 0.28 | 15/22 | 0.59 | 0.51 / 0.65 | 1/22 | $3 \times 10^{-4}$ |
| Angiosperms | 5 | 0.19 / 0.14 | 4/5 | 0.12 | 0.55 / 0.61 | 1/5 | 0.12 |

## Supplementary Data

**Supplementary Data 1.** Hi-C and Micro-C maps used in this study (one row per map; 149 maps). Columns give the organism, biosource, experiment type, number of biological replicates, data split (pretraining, or test maps held out for evaluation), release year, lab, data portal (ENCODE or 4DN), dataset and file accession codes, and the number of non-diagonal reads (millions).

**Supplementary Data 2.** Per-species results on the 177 DNA Zoo species (one row per species). Columns give the species, its clade, the patristic distance to *Homo sapiens* (million years; empty when no dated leaf is available), the sequencing depth of the evaluated map (contacts per 10 kb bin), the evaluated scaffold length, the number of 10 kb bins and of ground-truth TAD boundaries; the distance-stratified pixel-level PCC and the TAD-boundary *F*_1_ of Evo2HiC, its Hi-C–only ablation, Evo2HiC run with random DNA, HiCARN2 and HiCFoundation; and the number of boundaries called, the boundary precision and the boundary recall of Evo2HiC, Evo2HiC run with random DNA and the Hi-C–only ablation. Metrics are defined in Methods and summarized by clade in Supplementary Table 15.

